# Mechanical model of muscle contraction. 4. Theoretical calculations of the tension during the isometric tetanus plateau and the tension exerted at the end of phase 1 of a length step

**DOI:** 10.1101/2019.12.16.878843

**Authors:** S. Louvet

**Affiliations:** Institut P’. Physique et Mécanique des Matériaux. Université de Poitiers, Futuroscope Chasseneuil, France

## Abstract

Hypothesis 4 presented in accompanying Paper 2 states that the lever of a myosin II head in working stroke (WS) moves in a fixed plane, the orientation of the lever being defined by the angle θ. From this conjecture can be deduced the hypothesis 5 developed in accompanying Paper 3: the distribution of θ is identical and uniform in each half-sarcomere (hs) of a muscle fiber stimulated under isometric conditions. We propose a sixth hypothesis that establishes a linear relationship between the θ angle and the motor moment (*ℳ*) exerted on the lever. These three hypotheses lead to calculations of the tension during isometric tetanus plateau (T0) and the tension applied at the end of phase 1 of a length step when the only internal actions are the forces of elastic origin produced by the myosin heads in WS (T1_Elas_). However, the T1_Elas_ values are higher than those observed experimentally. The model introduces the presence of viscosity as the seventh hypothesis. The internal actions resulting from the coupling of the elasticity of the WS heads and the viscosity make it possible to explain all the observed phenomena that contribute to the phase 1 of a length step. An adequate adjustment between the theoretical tension from the model (T1) and the tension representative of the end of phase 1 exposed in examples from the physiological literature is proven (r^2^ > 98%). Other parameters such as stiffness (e), compliance (C) and strain (Y) are deduced; their investigation enables the construction of an analytical “nanoscope” by means of which the uniform density of θ is explored. The equations for T0, T1, e, C and Y explain and predict the influence of factors such as the duration of phase 1, the initial length of the sarcomere, the concentration of calcium, the presence of an inhibitor, the tension rise to the isometric tetanus plateau, relaxation after tetanization or shortening at constant speed. The results obtained during a slack-test are indicated by the model, the slack of the fiber being interpreted as an event of purely viscous origin.

## INTRODUCTION

### Perturbation by a length step

After being isometrically tetanized, the stimulated fiber is shortened or elongated rapidly by a length step (ΔL). Then the temporal evolution of the tension is observed. Four phases are distinguished, numbered from 1 to 4; see Table 1 in [1] and Table 4 in [2].

**Table 1.** Reference values for parameters related to the calculation of T1 concerning some fibers isolated from vertebrate muscles.

|  | From Fig 13<br>in Ford 1977 | From Fig 3B in<br>Brunello 2009 | From Figs 9C and 9D<br>in Linari 2004 | From Fig 1C in<br>Caremani 2008 |
| --- | --- | --- | --- | --- |
| Fig No. in<br>Paper 4 and<br>Suppl S4.J | 3B, 6<br>J8, J9, J10, J14 | 5<br>J13 | 4 | J11 |
| Animal | Rana T <sup>(1)</sup> | Rana E <sup>(2)</sup> | Man | Rabbit |
| Muscle | TAM <sup>(3)</sup> | TAM <sup>(3)</sup> | muscle<br>slow / fast | Psoas |
| $\Gamma$ | 2.5 °C | 4 °C | 12 °C | 12.4 °C |
| pCa | 4.5 | 4.5 | 4.5 | 4.5 |
| $\tau_{p1}$ | 200 μs <sup>(4)</sup> | 90 μs <sup>(5)</sup> | 110 μs <sup>(4)</sup> | 110 μs <sup>(4)</sup> |
| T0 | 245 kPa | 200 kPa | 66 kPa / 109 kPa | 156 kPa |
| L0 | 6 mm | 5 mm | 3.15 mm / 3.51 mm | 4.71 mm |
| $N_{hs}$ | 5500 | 4800 | 2800 / 3225 | 3300 |
| $L0_s$ | 2.2 μm | 2.14 μm | 2.4 μm / 2.45 μm | 2.47 μm |
| $\delta X_{Max}$ | 11.5 nm | 11 nm | 11 nm / 11.5 nm | 13 nm |
| $\delta X_T$ | 8 nm | 8 nm | 8 nm | 8.5 nm |
| $\delta X_{z1}$ | 4 nm | 3 nm | 3.5 nm / 4 nm | 5 nm |
| $Bz1_{min}$ | -3 nm | -1.7 nm | -2.6 nm / -3.4 nm | -3.4 nm |
| $Bz1_{Max}$ | -4.8 nm | -4.6 nm | -4.2 nm / -4.4 nm | -5.3 nm |
| $Bz2_{min}$ | -6.8 nm | -4.6 nm | -6.6 nm / -8.3 nm | -8.2 nm |
| $Bz2_{Max}$ | -17 nm | -24 nm | -16 nm / -15 nm | -17.7 nm |
| $\chi_{z1}$ | 0.133 nm<sup>-1</sup> | 0.133 nm<sup>-1</sup> | 0.143 nm<sup>-1</sup> / 0.133 nm<sup>-1</sup> | 0.113 nm<sup>-1</sup> |
| $\chi_{z2}$ | 0.062 nm<sup>-1</sup> | 0.067 nm<sup>-1</sup> | 0.067 nm<sup>-1</sup> / 0.057 nm<sup>-1</sup> | 0.054 nm<sup>-1</sup> |
| $K_{z1}$ | 1.35 | 2.05 | 1.34 / 1.185 | 1.31 |
| $K_{z2}$ | 1.7 | 2.8 | 1.67 / 1.38 | 1.61 |
| $q_{z1}$ | 1.99 | 1.84 | 1.99 / 2.07 | 2 |
| $q_{z2}$ | 1.9 | 1.76 | 1.89 / 1.97 | 1.91 |
| $r^2$ | 99.9% | 99.8% | 99.7% / 99.8% | 97.6% |

<sup>(1)</sup> *Rana Temporaria* <sup>(2)</sup> *Rana Esculenta* <sup>(3)</sup> *Tibialis Anterior* Muscle
<sup>(4)</sup> Length step <sup>(5)</sup> Sinusoidal oscillations at 4 kHz

### Phase 1 of a length step (ΔL<0)

The first transitional period of the length step called as “phase 1” is the shortening over a time lapse (τ_p1_), generally less than 0.2 ms [2.3], where the tension drops suddenly and linearly from T0 to T1, the minimum value of the tension reached at the end of phase 1 (Figs 1a and 1b). An index k is assigned to each length step (ΔL_k_) to which the minimum tension at the end of phase 1 (T1_k_) corresponds; see Fig 1b with 3 examples of length steps with index, k, (k+1) and (k+2), respectively. The index 0 corresponds to the case of the isometric tetanus plateau with ΔL_0_=0 and T1_0_=T0. For each step k, the shortening (ΔL_k_) is carried out during τ_p1_ with a constant shortening velocity (V_k_) but variable from one step to another (Fig 1a) since:

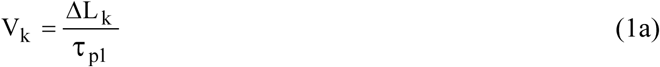

where τ_p1_ is the duration of the phase 1 identical for all length steps.

**Fig 1.**
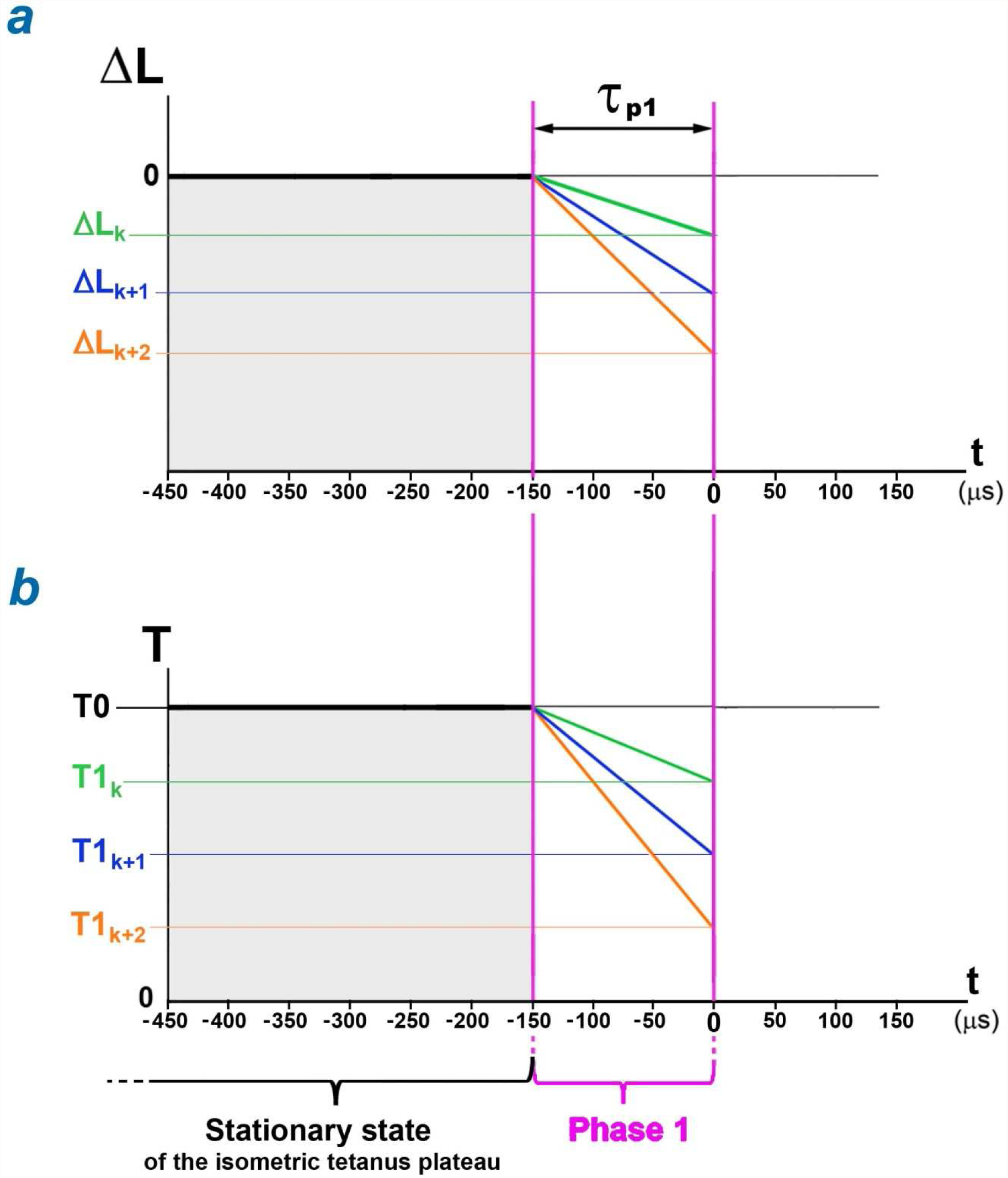
Phase 1 of a length step. (a) Length difference (ΔL) for a muscle fiber as a function of time (t), the stimulated fiber being shortened at constant speed during τ_p1_ according to three negative length steps, ΔL_k_, ΔL_k+1_ and ΔL_k+2_ with index, k, (k+1), (k+2), respectively. (b) Instantaneous tension (T) exerted at the fiber extremity during the three length steps; at the start of phase 1 corresponding to the end time of the isometric tetanus plateau, the tension drops abruptly and linearly during τ_p1_ from T0 to each of the three minimum values T1_k_, T1_k+1_ and T1_k+2_.

ΔL_k_ is trivially the sum of the shortenings of all half-sarcomeres (hs) of the myofibril as:

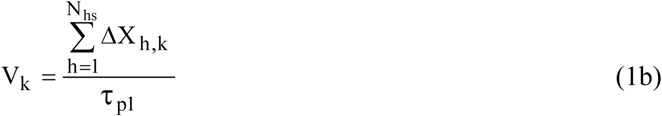

where ΔX_h,k_ is the shortening of hs n° h following the length step with index k; N_hs_ is the number of hs per myofibril.

In the rest of the paper, the k index of the length step will no longer be mentioned.

The average hs shortening 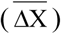 corresponding to the shortening ΔL of the fiber is:

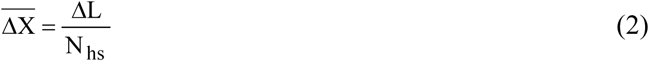

Our objective is to calculate tensions T0 and T1.

## Methods

### Hypothesis 6: Linearity between the motor-moment and the angular position of the lever

A myosin II head is traditionally modelled by 3 material segments articulated between them: the motor domain (S1a), the lever (S1b) and the rod (S2). The mechanical condition of a WS head is characterized by 4 conditions: 1/ the rigidity of S1a, S1b and S2; 2/ the strong bond between S1a and the actin molecule; 3/ a motor-moment (*ℳ*) exerted on S1b, a moment which induces a traction of the myosin filament via S2 and consequently the shortening of the hs; 4/ the displacement of the lever S1b in a fixed plane, the orientation of S1b in this plane being defined with the angle θ bounded by the 2 limits θ_up_ and θ_down_ corresponding to the classical *up* and *down* positions. The first 3 conditions were set in 1993 by I. Rayment [4]. Condition 4 is also suggested in the same article, but hypothesis 4 introduced in accompanying Paper 2 specifies its geometry. We formulate hypothesis 6 which states that the motor-moment *ℳ* is an affine function of θ (Fig 2; red line):

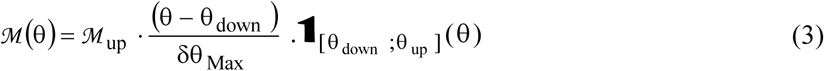

where *ℳ*_up_ is the maximum moment corresponding to the angle θ_up_; δθ_Max_ is the angular range between θ_up_ and θ_down_, two angles whose values are calculated, respectively, in (12a) and (12b) in Paper 2; **1** is the symbol characterizing the indicator function defined in (A2b) in Supplement S1.A of Paper 1.

**Fig 2.**
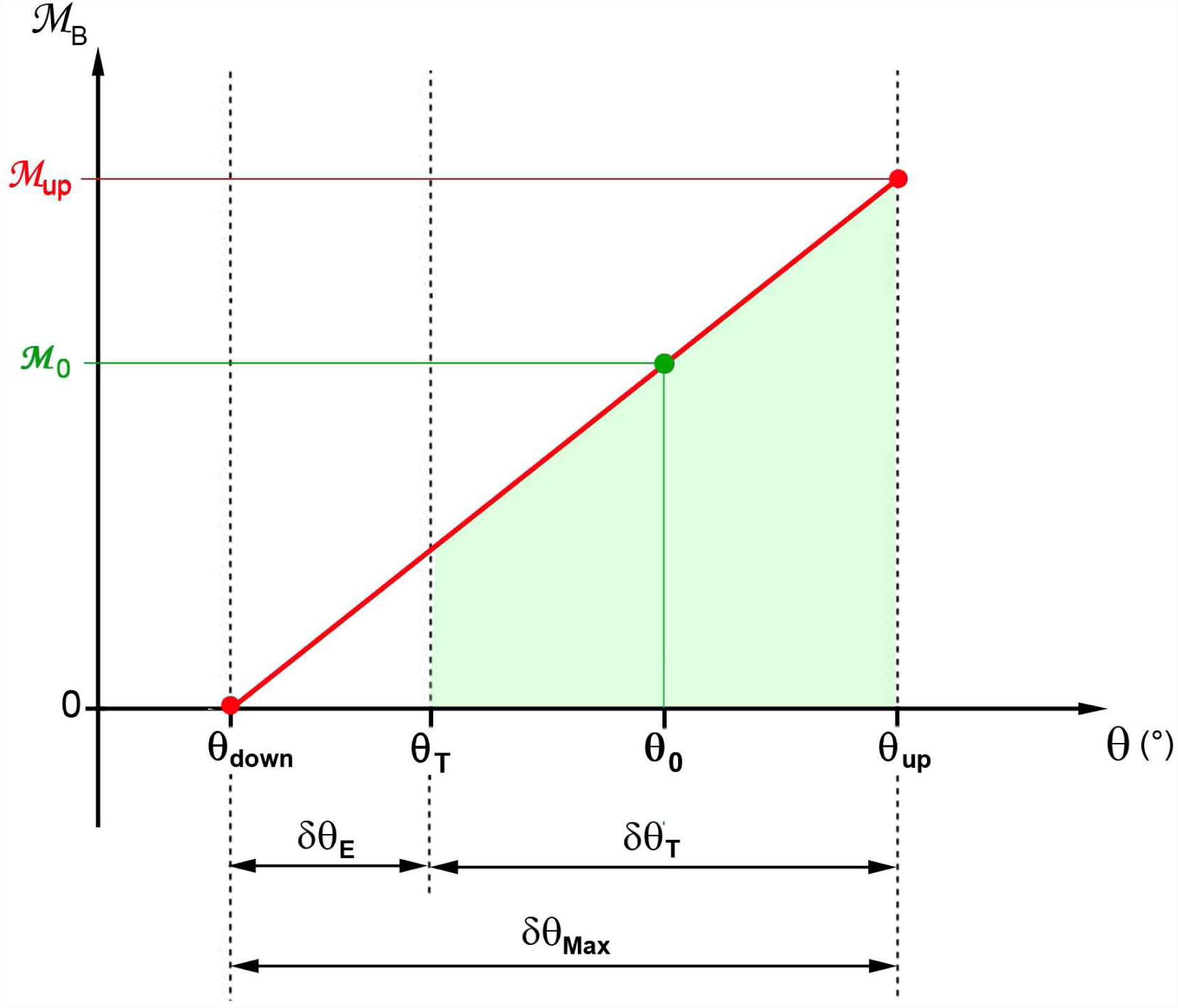
Affine relationship between the motor-moment *ℳ* and the θ angle of lever S1b in a half-sarcomere on the right. Linearity is represented by a red line. The green trapezoidal area whose base is the extent δθ_T_ characterizes the domain where the orientation θ of the Λ_0_ levers belonging to the Λ_0_ WS heads is distributed uniformly between θ_T_ and θ_up_. The rectangular triangle based on the δθ_E_ range is empty because all WS heads with a lever orientation between θ_down_ and θ_T_ have come off during the rise to tetanus plateau.

Several researchers [5,6,7,8,9] have previously enunciated this conjecture without formalizing it. The motor-moment (*ℳ*) derives from a potential elastic energy stored in the β-sheet element of the motor domain of the myosin head II; see paragraph C.3 of Supplement S2.C of Paper 2.

### Isometric tetanus plateau and calculation of T0

An isolated muscle fiber at rest is stimulated to a fixed length by pulses until fused tetanus. The measured tension increases and then reaches a quasi-constant maximum value (T0) which characterizes a stationary equilibrium (Fig 1b) called as “isometric tetanus plateau”. The plateau duration can be longer than the second [10]. During the tetanus plateau, the spatial density of θ is observed over an angular range (δθ_T_) between 40° and 50°, postulated as uniform by various authors [11,12,13,14,15], result found after geometric modelling of a hs in Paper 3 and theorized with hypothesis 5. The maximum variation of θ (δθ_Max_) between the two limits θ_up_ and θ_down_ relating to the two positions *up* and *down* is equal to 70°. The range δθ_T_ is framed by the two angles θ_T_ and θ_up_. During the isometric tetanus plateau, there is an interval (δθ_E_) of about 20° between θ_T_ and θ_down_ where no head is found in WS state (Fig 2). This absence is explained by the slow detachment of the myosin heads in the range δθ_E_ during the tension rise to the tetanus plateau; see accompanying Paper 1 for further details.

The angle θ_0_ is defining as located at the middle of δθ_T_ (Fig 2):

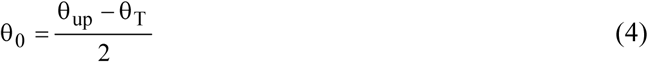

For example, in support of the data of the Table 1 in Paper 3, the values of θ_T_ and θ_0_ are −21° (+21°) and +3.5° (−3.5°) in a hs on the right (left), respectively.

With hypothesis 5 of uniform density of θ between θ_T_ and θ_up_ and by introducing equality (4) in equation (3), we obtain the average moment (*ℳ*_0_) generated by all the motor-moments present during an isometric tetanus plateau (Fig 2):

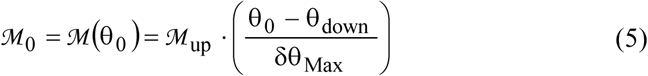

The study in Fig 2 provides:

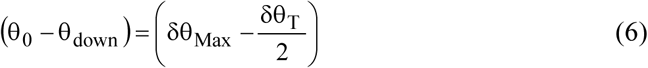

And equality (5) is rewritten with (6):

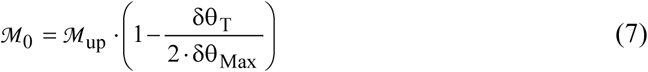

The average moment *ℳ*_0_ depends on 3 parameters: *ℳ*_up_ is the maximum motor moment; δθ_T_ is the range of the uniform law of the random variable Θ associated with the angle θ during the isometric tetanus plateau; δθ_Max_ is the maximum variation of θ during the WS.

T0 is calculated in (I18) in Supplement S4.I. With (6), T0 is equal to:

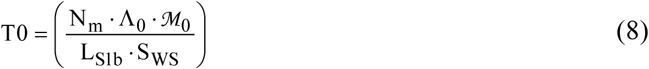

where N_m_ is the number of myofibrils of the fiber; Λ_0_ is the number of identical and constant WS myosin heads per hs during the tetanus plateau in isometric conditions; L_S1b_ is the length of the lever; S_WS_ is a characteristic parameter of the myosin head defined in equations (13) and (14) of Paper 2 whose value is near 0.95.

### Linear relationship between ΔX and Δθ

In the linear domain defined by the terminals θ_up_ and θ_down_, the relationship between the hs displacement (ΔX) and the lever rotation (Δθ) is established in (19) in Paper 2 and duplicated below:

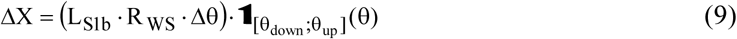

where R_WS_ is another characteristic parameter of the WS myosin head; R_WS_ is determined with equalities (13) and (14) in Paper 2 and its value is equal to S_WS_.

From (9) are deduced the equivalence relationships between angular and linear ranges developed in Supplement S4.I; the equations (I22), (I23), (I24) and (I27) are reproduced below:

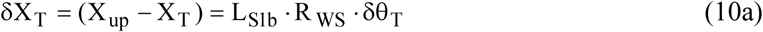

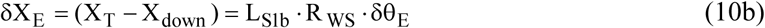

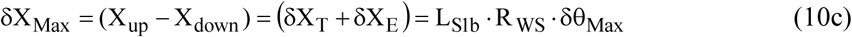

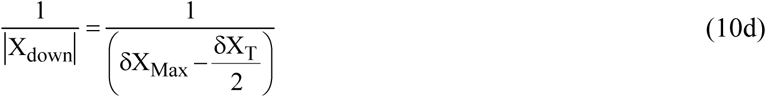

where X_up_, X_down_, X_T_, X_E_ are the abscissa corresponding to the angles θ_up_, θ_down_, θ_T_, θ_E_ (see Fig I1).

### Characterization of T1_Elas_ with internal actions calculated in the absence of viscosity

We consider the hs shortenings inferior in modulus at δX_Max_. In this case, the linear interval [-δX_Max_;0] is separated into 2 zones (Fig 3) defined in paragraph I.6 of Supplement S4.I:

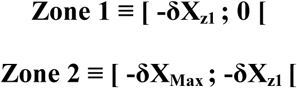

where δX_z1_ is ranged between δX_E_ and (2·δX_E_); δX_E_ and δX_Max_ are two linear ranges specified in (10b) and (10c).

**Fig 3.**
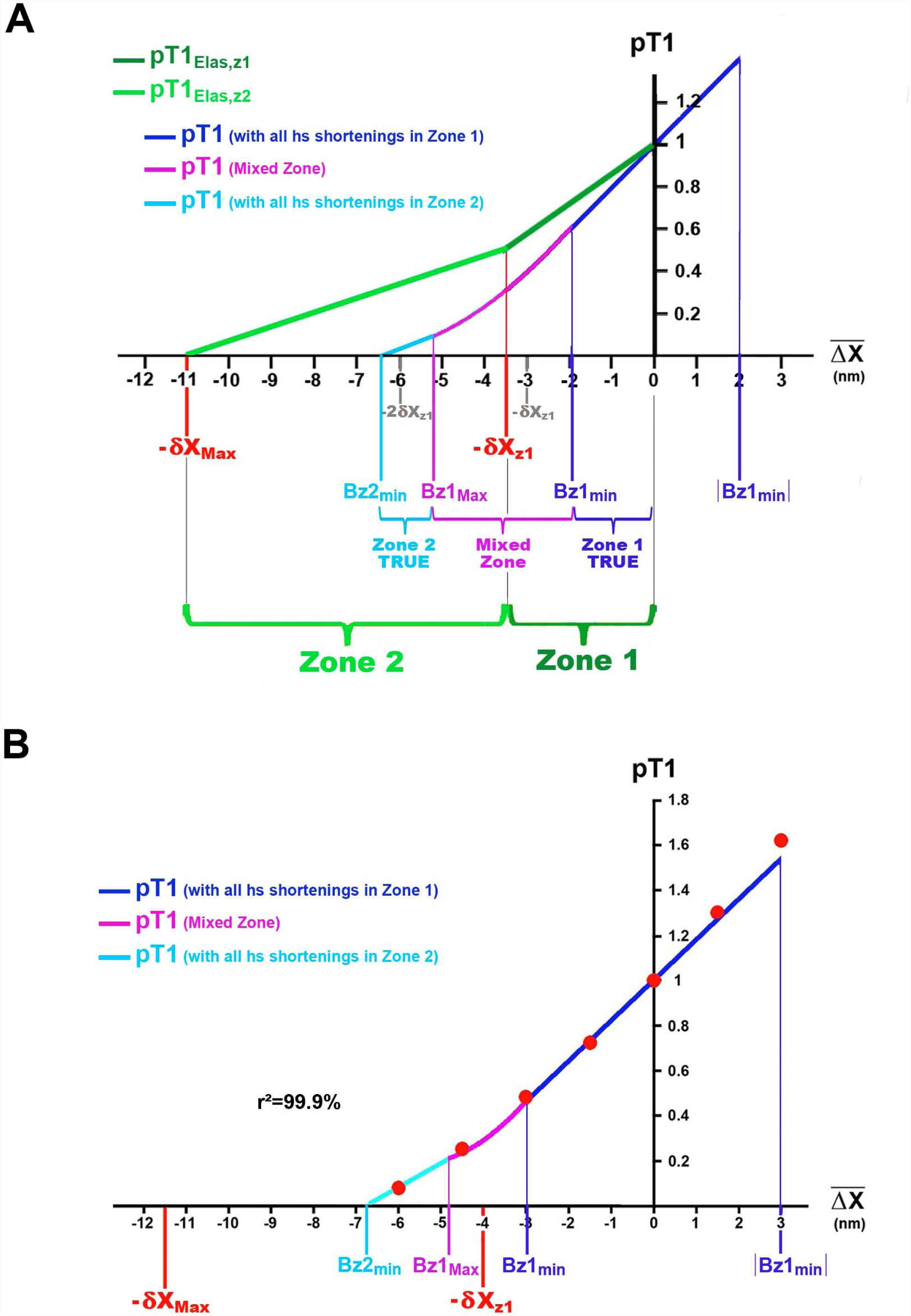
Evolution of pT1, the relative tension of the fiber at the end of phase 1 of a length step as a function of the average hs shortening 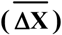. (A) Theoretical model: pT1_Elas_ is calculated with expressions (10) and (13) in Zone 1 (dark green segment) and Zone 2 (light green segment); pT1 is determined using equations (20) and (21) in Zone 1 True (dark blue segment), in the Mixed Zone (mauve parabolic arc) and in Zone 2 true (sky blue segment). (B) Application of the model to a fiber isolated from the *tibialis anterior* muscle of *rana Temporaria*; the red dots are from Fig 13 in [2], reproduced in Fig J8 of Supplement S4.J.

When the internal actions are only the linking forces and moments, equations (I52a) and (I53a) of sub-paragraph I.5.4 of Supplement S4.I provide the relative tension (pT1_Elas_) exerted at the fiber extremities at the end of phase 1; pT1_Elas_ is calculated in Zones 1 and 2 according to:

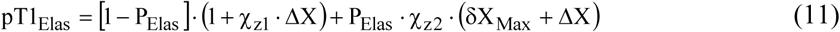

where ΔX is the same shortening of all hs in the fiber; with (2), ΔX is equal to:

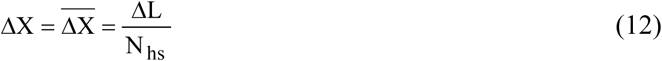

where P_Elas_ is a weight as a function of ΔX, equal to 0 or 1, such that :

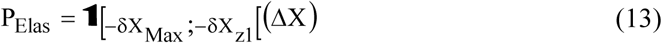

where χ_z1_ and χ_z2_ are two slopes of elastic origin equal to:

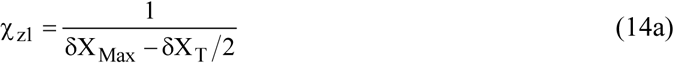

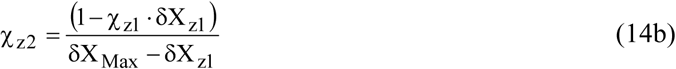

With this modeling, the fiber behaves like a linear-elastic or Hookean spring with stiffness χ_z1_ or χ_z2_ depending on the zone where ΔX is located. The relationship (11) is represented in Fig 3A by two straight line segments coloured in dark and light green in Zones 1 and 2, respectively.

If the tensions measured in reality are compared with the theoteritical values determined with equation (11), the experimental tensions are generally much lower. An example is provided in Fig J9 of paragraph J. 9 of Supplement S4.J where all the points reproduced are from Fig 19 in [2]. The dark blue experimental dots associated with τ_p1_=0.2 ms are displayed well below the two light green segments representative of pT1_Elas_ determined with (11).

The linking actions related to WS heads are derived from an elastic potential energy. By definition, these actions depend only on the distance covered and not on the time taken to cover it. In the same figure J9, the light green dots associated with τ_p1_=1 ms are located above the dark blue dots and are close to the two light green segments representative of pT1_Elas_. This observation suggests the presence of other braking forces depending on the duration of phase 1 and therefore on the velocity according to equality (1b). Hypothesis 7 introduces viscosity as the candidate for the position.

### Characterization of T1 with internal actions calculated in the presence of viscosity

All the following acronyms, equalities and equations are explained in Supplement S4.J. A viscous parameter (ν), common to all hs of the fiber and specific to each experiment, is introduced:

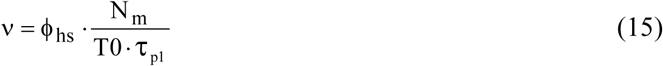

where ϕ_hs_ is the proportionality coefficient in the presence of viscosity, characteristic and common to both massive sets of a hs, the M-disk and Z-disk; N_m_ is the number of myofibrils of the fiber.

The value of ν calculated experimentally at 2°C is very low, in the order of 10^−8^ nm^−1^.

q_z1_ and q_z2_ are two parameters constitutive of the presence of viscosity depending on whether the hs shortening (ΔX) is in Zone 1 or Zone 2 defining as:

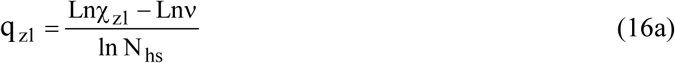

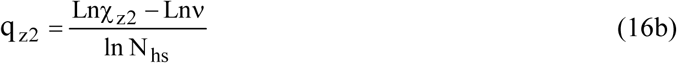

where Ln is the symbol for the natural logarithm; χ_z1_ and χ_z2_ are the two slopes of elastic origin formulated in (14a) and (14b); N_hs_ is the number of hs per myofibril.

The two parameters q_z1_ and q_z2_ are linked to each other by:

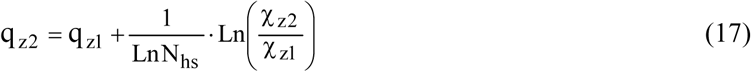

Relatively and respectively to q_z1_ for Zone 1 and q_z2_ for Zone 2, the two coefficients K_z1_ and K_z2_ are assigned as:

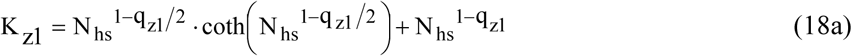

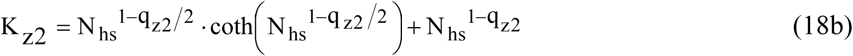

where coth is the symbol of the hyperbolic cotangent with coth(x) = (e^2x^+1) / (e^2x^-1).

A representation of K_z1_ (K_z2_) as a function of q_z1_ (q_z2_) is given in Fig J3 of Supplement S4.J. Parameter K is a decreasing function of q such that:

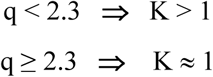

Four abscissa characteristic of the presence of viscosity (Bz1_min_, Bz1_Max_, Bz2_min_, Bz2_Max_) are defined from the four variables, q_z2_, q_z2_, K_z1_, K_z2_, that is:

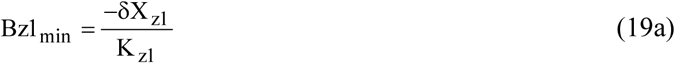

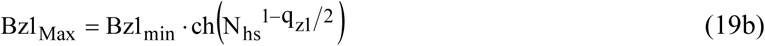

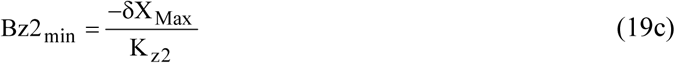

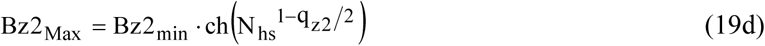

where ch is the symbol of the hyperbolic cosine with ch(x) = (e^x^+e^−x^)/2.

Three of these four abscissa are shown in Fig 3A, the calculations being made with the data displayed in the green column of Table 1. They are used to delimit three zones, Zone 1 True, Zone 2 True and Mixed Zone, defined in the following sub-section.

When the internal forces consist of the linking actions of elastic origin on the one hand and the forces due to viscosity on the other hand, the relative tension (pT1) exerted at the extremities of the fiber at the end of phase 1 is formulated according to (J48), equation reproduced below:

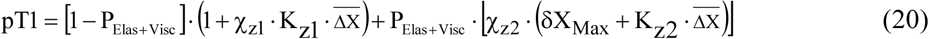

where 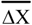 is the average shortening calculated in (2); K_z1_ and K_z2_ are determined in (18a) and (18b) and are two multiplying coefficients greater than 1 relative to the two slopes of elastic origin, χ_z1_ and χ_z2_;

P_Elas+Visc_ is a weight as function of 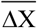, varying between 0 and 1, such that:

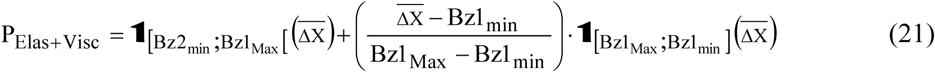

In the absence of viscosity, i.e. K_z1_= K_z2_ = 1 with q_z2_ ≥ 2.3, relationships (20) and (21) becomes (11) and (13), respectively.

### Zone 1 True, Zone 2 True and Mixed Zone

The introduction of viscosity forces into the equations implies that the shortenings of the hs of a myofibril are no longer equal; see series of inequalities in (J14). The variability of the lengths of the hs depends on the coefficients q_z1_ and q_z2_. Figure J4 shows the progression of the 1000 shortenings of the 1000 hs composing a myofibril for 3 values of q_z1_ (or q_z2_).

Following these remarks, equation (20) presents 3 different shapes depending on whether the shortening 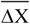 belongs to one of the following three intervals (Fig 3A):

**1/ Zone** 1 **True** 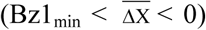: the shortenings of all hs are in Zone 1 and the relationship between pT1 and 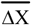 is a straight line segment of slope (χ_z1_ K_z1_) traced in dark blue.

**2/ Mixed Zone** 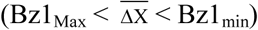: the shortenings of the most distal hs have passed into Zone 2 while those of the proximal hs are still in Zone 1; the relationship between pT1 and 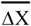 is a convex parabolic arc traced in violet.

**3/ Zone 2 True 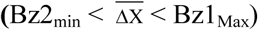**: the shortenings of all hs are in Zone 2 and the relationship between pT1 and 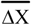 is a straight line segment of slope (χ_z2_ K_z2_) traced in light blue.

The value of Bz2_min_ delivered in (19c) is the value for which the tension pT1 is cancelled (Figs 3A and 3B). Consecutively the tension T1 is zero or negative for shortenings below Bz2_min_.

### Zone 1 Enlarged

The curvature of the arc of the parabola in Mixed Zone (Figs 3A and 3B; purple line) is little between −δX_z1_ and Bz1_min_. The equation (20) determined for the Zone 1 True remains valid for the entire Zone 1, to which it is possible to add a zone of elongations with the interval [0; |Bz1_min_|]; see paragraph J.5 of Supplement S4.J. So the tension at the end of phase 1 is formulated between −δX_z1_ and |Bz1_min_|:

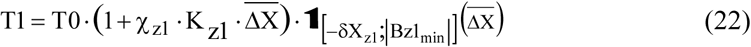

Zone 1 Enlarged is displayed in Fig J5a to Supplement S4.J.

### Stiffness and strain

The stiffness (e_0_) is the slope of the straight line for equation (22):

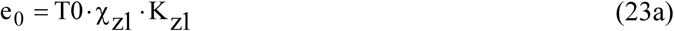

The strain (Y_0_) is the absolute value of 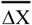 for which the tension T1 calculated in (22) is cancelled:

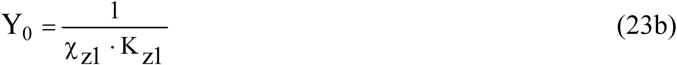

The following equality is noted:

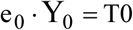

### Instant tetanus tension (T0_i_) and instant tension at the end of phase 1 (T1_i_)

A fiber at rest is tetanized in isometric conditions. When rising to the isometric tetanus plateau (T0_c_), the instant tension [T0i(t)] varies from 0 to T0_c_. If, during the rise, a series of length steps is practiced, the instantaneous tension collected at the end of phase 1 [T1_i_(t+τ_p1_)] for each step is calculated in Zone 1 Enlarged from the relationship (22) reformatted according to:

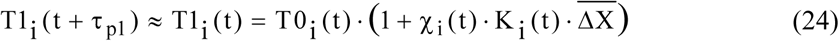

where χ_i_ is the instant coefficient of elastic origin stiffness evaluated from (14a):

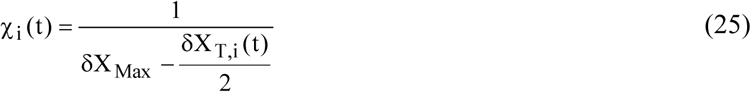

where δX_T,i_ is the instant linear range associated with the instant angular range (δθ_T,i_) over which the orientation of the levers belonging to the WS heads is uniformly distributed at time t.

and where K_i_ is the instant multiplier coefficient determined according to (18a):

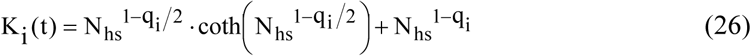

where q_i_ is the instant parameter, constitutive of the presence of viscosity in the instantaneous experiment; q_i_ is calculated according to the equation (J57):

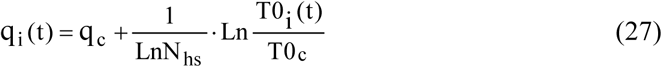

where q_c_ is the characteristic parameter of the presence of viscosity in the control experiment for which the tension of the isometric tetanus plateau is equal to T0_c_.

Equations (24) to (27) are demonstrated in paragraph J.11 of Supplement S4.J. These equations also apply to time-independent experimental series, such as force step series or length step series at different intracellular concentrations of calcium, inorganic phosphate or inhibitor of cross-bridges. In these cases, T0_i_ is referred to as “intermediate tetanus tension”.

Following expressions (23a) and (23b), the instantaneous normalized stiffness (e_i_/e0) and the instantaneous strain (Y_i_) are defined from equation (24):

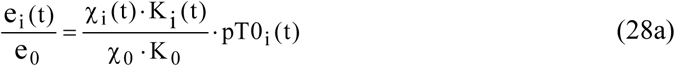

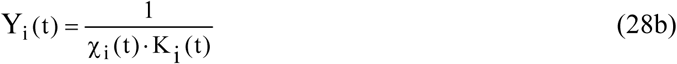

where the constants χ_0_ and K_0_ are equivalent to χ_z1_ and K_z1_.

In support of (28a) and (28b), the instantaneous slope (χ_i_) is calculated relative to e_i_/e_0_ and Y_i_:

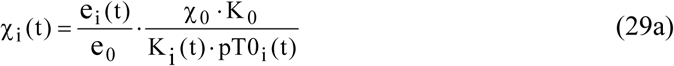

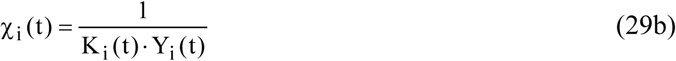

The stroke size (δX_Max_) is a constant, so the slope χ_i_ depends only on the instant linear range (δX_T,i_) according to (25). Relatively to e_i_/e_0_ and Y_i_ with (29a) and (29b), we obtain:

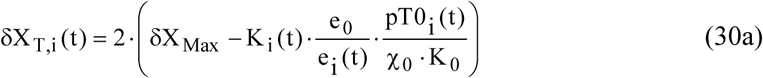

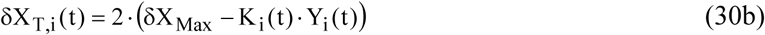

With the linear relationship (9), the corresponding instant angular range (δθ_T,i_) is equal to :

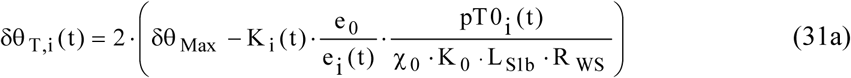

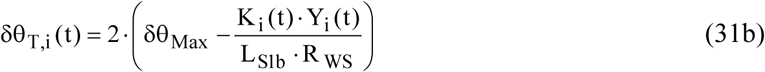

A hs of the fiber is thus interpreted as a controlled motor system that responds to a perturbation between two particular equilibrium states:

Equilibrium 1 with pT0_i_=0 and δθ_T, i_=0 : characteristics of the rest or total relaxation where no cross-bridge is present

Equilibrium 2 with pT0_i_=1 and δθ_T,i_=δθ_T_ : determining conditions of the isometric tetanus plateau

Thanks to the expressions of δθ_T,i_ formulated in (31a) and (31b), we have an theoretical “nanoscope” that allows to study the evolution of the uniform density of the θ angle of the levers belonging to the WS heads between the two equilibrium states. The adequate response of δθ_T,i_ as a function of pT0_i_ is broken down into a non-stationary phase with critical regime followed by a stationary phase with stable state representative of one of the two equilibrium states which acts as a setpoint to respect. The response is based on 4 equations formulated from (J76) to (J79) in sub-paragraph J.10.7 of Supplement S4.J relating to various experiments, such as the introduction of a cross-bridge inhibitor, the tension rise to the isometric tetanus plateau, relaxation and shortening at constant velocity.

### Objective

The purpose of the paper is to compare the previous equations with the experimental points of the T1/ΔX relationship presented in figures from the physiological literature.

### Statistics

A linear regression is performed between the measured tension values (points recorded on the figures of the articles listed) and the theoretical tension values calculated at the same velocity. The regression line goes through the origin, such that:

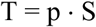

where T characterizes the values of the theoretical tensions and S those of the experimental tensions; p is the slope of the regression line equal to:

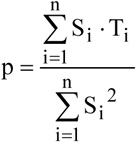

The determination coefficient is defining as:

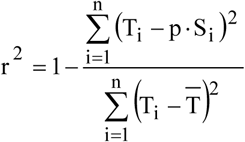

### Algorithmic

The sequence of computer programs is written under Visual Basic 6.

In a first step, the relationships developed in Supplement S4.J were verified, particularly the equality (J18) which attests to the equivalence between the two terms R_γ,n_ and Q_q,n_ (Fig J2). Equality (J18) is used to validate the calculation of the multiplying coefficients K_z1_ and K_z2_ given in (18a) and (18b) as well as the relationship between pT1 and 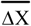 formulated in (20).

Equations (11) to (31b) were put into algorithms to obtain the plots of the relationships 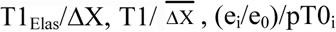, Y_i_/pT0_i_, δθ_T,i_/pT0_i_ and δX_T,i_/pT0_i_.

### Adequacy between experimental points and theoretical layout

For each curve, the adjustment is made visually by the trial and error method by searching for determination coefficient (r^2^) closest to 1 and allocating values to the data relating to myosin heads that are compatible with those in the literature and with those used in the calculations of Papers 1 to 6.

## Results

An example of the T1 calculation according to the theoretical equations (20) and (21) is given in paragraph J.6 of Supplement S4.J using the data displayed in the violet column of Table 1. The result of the modelling is shown in Fig 3B where the red dots come from Fig 13 in [2].

### Influence of the duration of phase 1 (τ_p1_)

A demonstration is provided in paragraph J.9 of Supplement S4.J and is illustrated in Fig J9, the points of which are taken from Fig 19 in [2].

### Influence of the intermediate tetanus tension (T0_i_)

A complete analysis where T0_i_ is determined from the length of the sarcomere ranging from 2.2 µ m to 3.25 µm is undertaken in paragraph J.10 of Supplement S4.J; see Figs J10a and J10b whose points are collected, respectively, from Figs 6 and 11 in [7].

The equations of the model are tested with another example: T0_i_ is evaluated on the basis of the calcium concentration (pCa) in slow and fast fibers collected from human muscle.

1/ With an intracellular calcium concentration (pCa_0_) equal to 4.5, length step series imposed on a slow fiber and a fast fiber serve as control experiments. Data for this pCa level are presented in the dark blue column of Table 1.

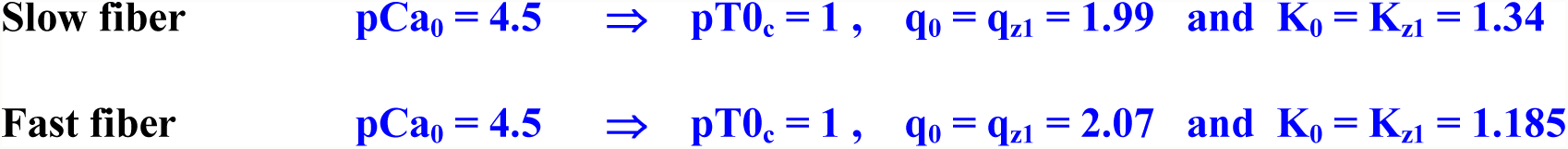

In support of equation (20), the two 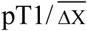 plots for the two types of human fibers in control experiments appear as a blue line on Figs 4a and 4b.

**Fig 4.**
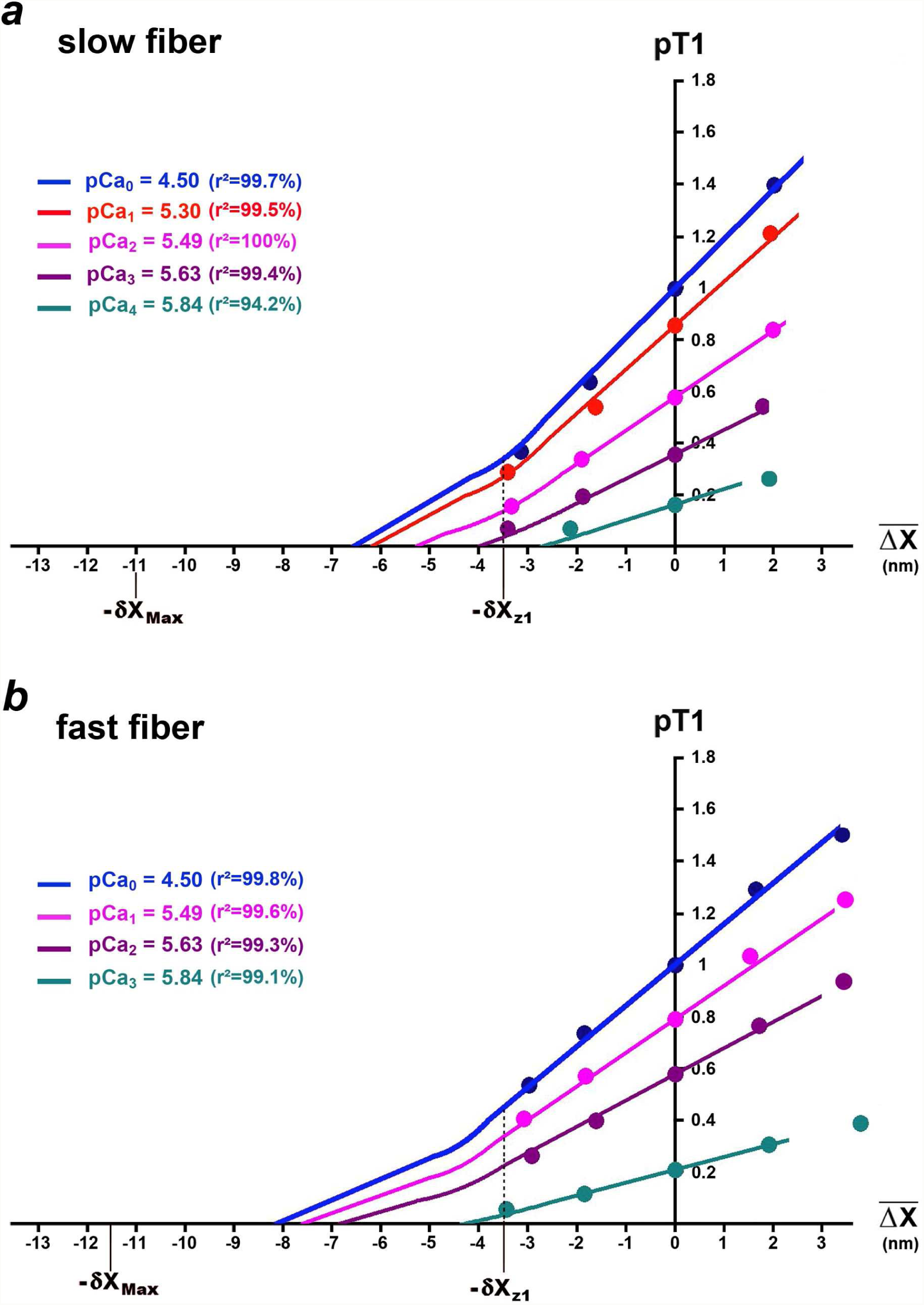
Relations of the intermediate tension at the end of phase 1 of a length step (pT1_i_) as a function of the average hs shortening 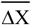 according to the intracellular calcium concentration (pCa_i_). (a) and (b) Experiments performed on a slow and fast human fiber, respectively. All points are from Figs 9C and 9D in [16].

2/ An intermediate experiment is performed on the same fiber where the experimental conditions are similar except for the concentration of intracellular calcium (pCa_i_) and consequently the intermediate tension of the isometric tetanus plateau (T0_i_); see Figs 9A and 9B in [16]. The viscous parameter (q_i_) and the associated multiplicative coefficient (K_i_) are calculated in Zone 1 Enlarged according to equations (27) and (26), respectively, for each intermediate experiment.

### Application

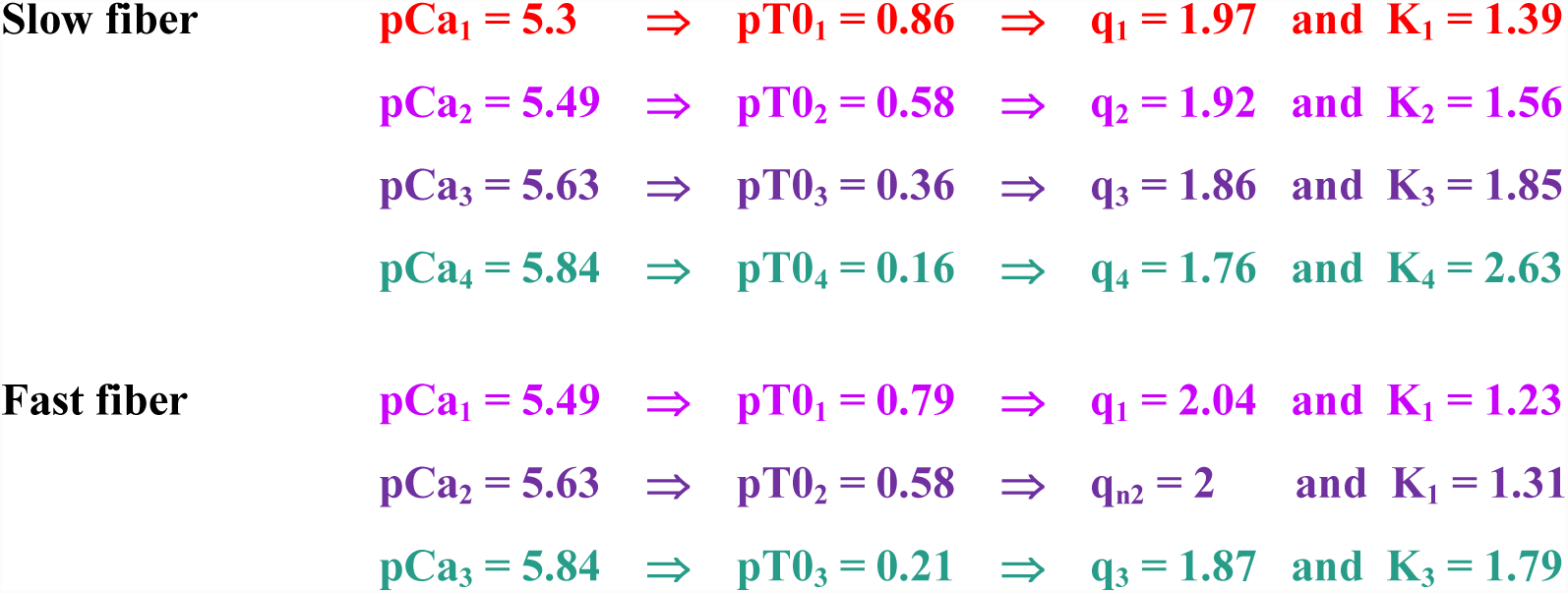

By following the method recommended in paragraph J.6, it is possible to determine all the parameters necessary for the construction of the pT1 curves as a function of 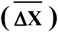 for each calcium concentration (Figs 4a and 4b). Between the experimental pT1 values collected in Figs 9C and 9D in [16] and the theoretical results, there is a good fit (r^2^>99%) with one exception.

With the expression (27), the mere knowledge of the pT0_i_ value corresponding to the intermediate concentration of calcium tested (pCa_i_) is sufficient to calculate the intermediate relative tension (pT1_i_) at the end of phase 1 of a length step.

### Study of the instant strain (Y_i_) during the tension rise to the isometric tetanus plateau followed by isometric relaxation

An isolated fiber at rest is isometrically tetanized for 300 ms, the tetanus plateau being reached in 100 ms. After 300 ms, the stimulation is stopped and the fiber maintained under isometric conditions is relaxed for 300 ms; see Fig 1 in [17]. The fiber is tested during the 600 ms (tetanization and relaxation) using sinusoidal oscillations in length (2 nm per hs peak to peak) at a frequency of 4 kHz. With the values displayed in the green column of Table 1 as reference data, the instant viscous parameter (q_i_) is expressed as a function of the instant tetanus tension (pT0_i_) according to (27):

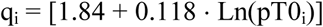

The theoretical and associated instant multiplier coefficient (K_i_) is determined with (26).

#### 1/ Rise to the isometric tetanus plateau

The instant linear range (δX_T,i_) is empirically modelled according to equation (J78a) presented in sub-paragraph J. 11.7 of Supplement S4.J, i.e. after affine transformation using (9):

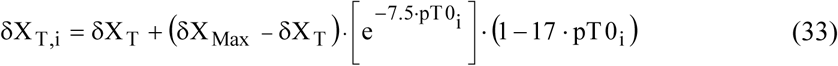

Equation (33) is represented by a light green line in Fig 5a. The light green points are calculated from equation (30b) where are introduced the experimental values of Y_i_ collected on Fig 6B in [17].

Knowing δX_T,i_ modelled by equation (33), the instant elastic origin slope (χ_i_) is determined with (25). The instant strain (Y_i_) is calculated from (28b) and its graphical representation appears as a light green line in Fig 5b. The light green dots are the reproduction of the solid circles in Fig 6B from [17]. There is a good agreement between theoretical and experimental values.

**Fig 5.**
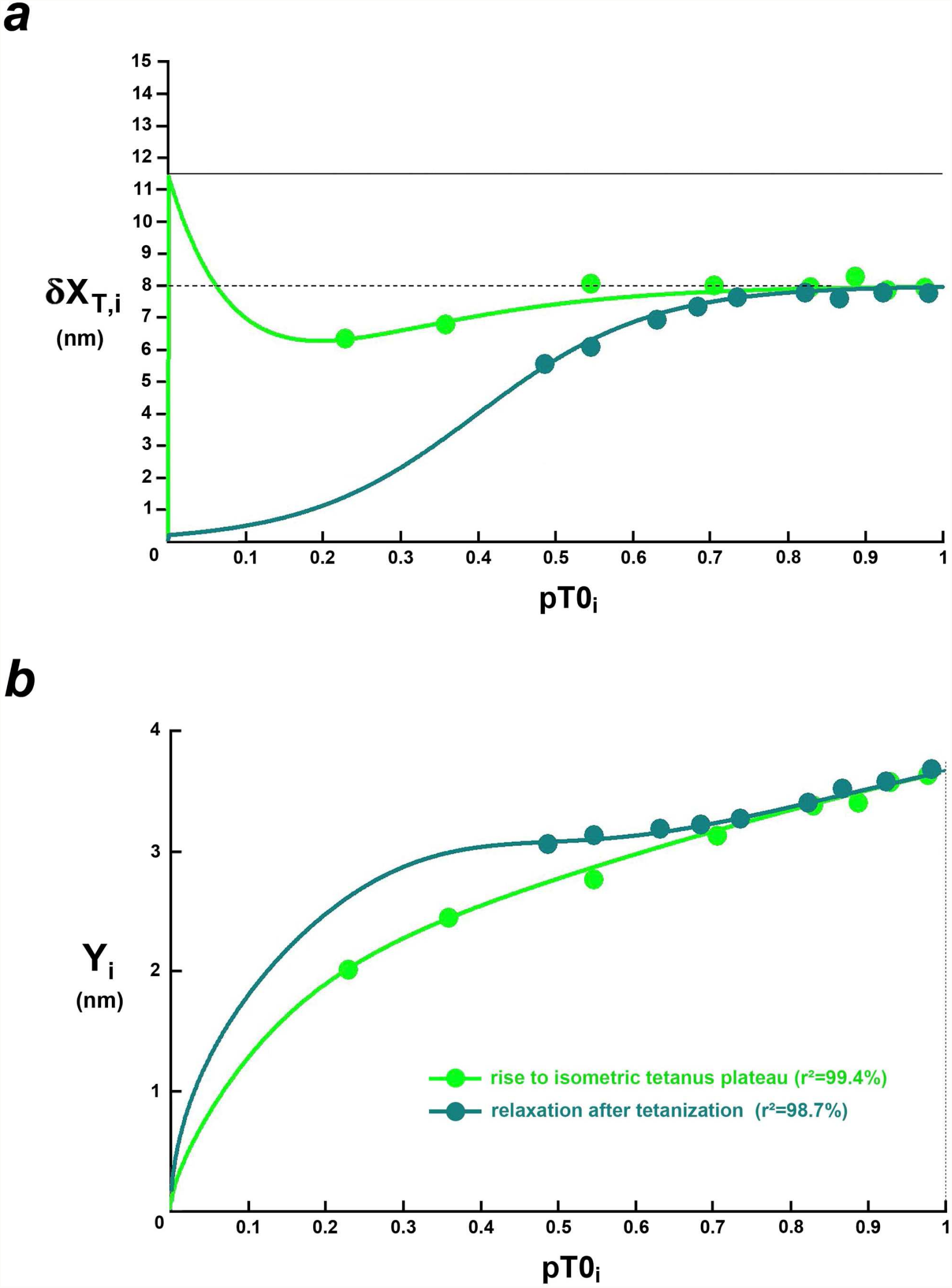
Relations of the instantaneous linear range (δX_T,i_) and the instantaneous theoretical strain (Y_i_) as a function of the instantaneous relative tension (pT0_i_) during the rise to the tetanus plateau and in the relaxation phase. (a) Curves of δX_T,ii_ according to (33) for the rise to the isometric tetanus plateau (light green line) and (34) for isometric relaxation (peacock blue line). The light green and peacock blue points are calculated using (30b) with the experimental values of Y_i_. (b) Curves of Y_i_ for the rise to the isometric tetanus plateau (light green line) and for isometric relaxation (peacock blue line). The light green and peacock blue dots are shown in Fig 6B from [17].

**Fig 6.**
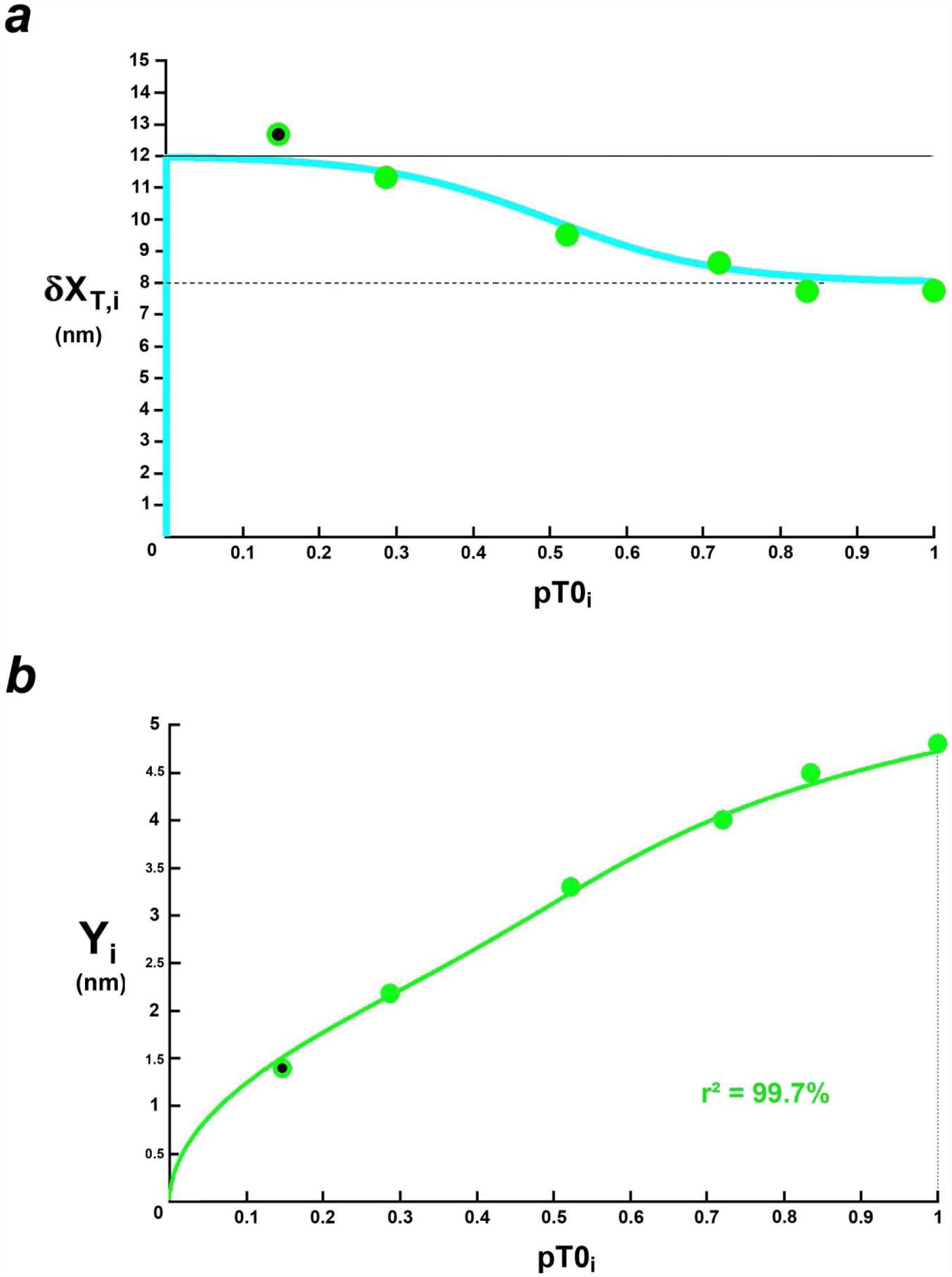
Relations of the intermediate linear range (δX_T,i_) and the intermediate theoretical strain (Y_i_) as a function of the intermediate relative tension (pT0_i_) during phase 4 of force steps. (a) Curve of δX_T,i_ (light blue line) according to (36); green points are calculated using (30b) in support of experimental values of Y_i_. (b) Y_i_ relation (light green line) with (28b); the green dots are recorded in Fig 3C from [13].

#### 2/ Isometric relaxation

The parameters q_i_ and K_i_ are determined in the same way.

The instant linear range (δX_T,i_) is modelled according to (J79), i.e. after affine transformation according to (9):

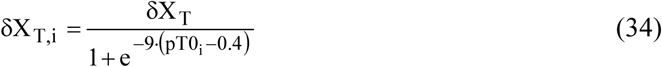

Starting from the tetanus plateau (pT0_i_ = 1), equation (34) is a sigmoid represented by a peacock blue line on Fig 5a, decreasing from δθ_T_ to 0. Peacock blue points are calculated from equation (30b) where the experimental values of Y_i_ are introduced.

Knowing δX_T,i_ modelled by (34), the instant slope χ_i_ is evaluated with (25). The theoretical instant strain (Y_i_) is calculated from (28b) and its curve appears as a peacock blue line in Fig 5b. The peacock blue dots are the reproduction of the empty circles in Fig 6B in [17]. There is an acceptable agreement between theoretical and experimental values.

### Analysis of the intermediate strain (Y_i_) during phase 4 of a series of force steps where tension and shortening velocity are steady

During phase 4 of a force step, the fiber is tested using sinusoidal oscillations in length (2 nm per hs peak to peak) at a frequency of 4 kHz. Such oscillations are approximated by a succession of length steps whose duration of phase 1 is equal to 90 µs; see equality (J46) in paragraph J.5 of Supplement S4.J. With the data presented in the purple column of Table 1, the value of q_i_ is evaluated according to the equality (J68) explained in sub-paragraph J. 11.3 of the Supplement S4.J, that is:

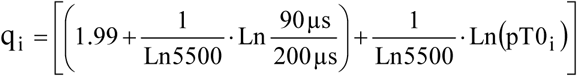

After computation:

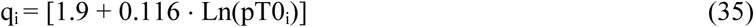

The corresponding theoretical factor K_i_ is determined according to (26). The intermediate angular range (δθ_T,i_) is empirically modelled according to (J80), i.e. after affine transformation using (9):

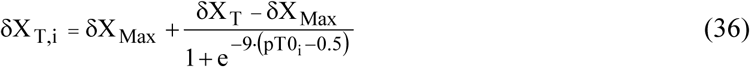

where δX_Max_ =12 nm and δX_T_ = 8 nm.

Equation (36) is a sigmoid represented by a light blue line in Fig 6a; from the tetanus plateau (pT0_c_=1) to almost total relaxation (pT0_i_ ≈ 0), the intermediate linear range (δX_T,i_) increases from δX_T_ to δX_Max_ The green points are calculated using equation (30b) where are introduced the experimental values of Y_i_ collected on Fig 3C in [13]. Knowing δX_T,i_, we determine in support to (25) the intermediate elastic origin slope (χ_i_). The theoretical intermediate strain (Y_i_) is calculated from (28b) and its curve appears as a green line in Fig 6b. The green dots come from Fig 3C in [13]. There is a good agreement between theoretical and experimental values.

The points in Figs 6a and 6b are numbered from the highest tension (i=0 for pT0_c_=1) to the lowest tension (i=5 for pT0_5_=0.15). On Fig 6a, the value of δX_T,5_ for the bicolor (green and black) point n° 5 is higher than δX_Max_. The explanation of this anomaly is provided in accompanying Paper 1 where the action of viscosity is important at high shortening velocities, i.e. superior at 1.5 nm·ms^−1^; see Fig 3 of Paper 1. Yet the hs shortening velocity of point n° 5 (u_5_) is 1.9 nm·ms^−1^ per hs as shown in Fig 3A from [13]; the viscous forces created by continuous shortening overlap with the viscous forces generated by oscillations with amplitude of 2 nm.hs^−1^ peak to peak.

In Fig 6b, point n° 5 has coordinates pT0_5_ = 0.146 and Y_5,mes_ = 1.4 nm. It is checked:

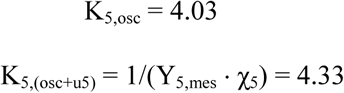

where K_5,osc_ is the multiplier coefficient induced by viscosity derived from the oscillations alone, theoretically calculated with (26), q_i_ being evaluated using (35); K_5,(osc+u5)_ is the multiplier coefficient resulting from viscous actions caused by both oscillations and high shortening velocity (u_5_), K_5,(osc+u5)_ is determined from (28b); χ_5_ is the elastic origin slope evaluated according to (25) where the linear range δX_T,5_ is determined with (36).

We deduce the multiplier coefficient elicited only from viscosity imposed by u_5_ (K_u5_):

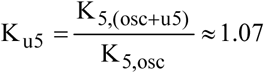

It is a modest figure compared to K_5,osc_; the result is comparable to the value found in sub-paragraph J.15.3 determined from the normalized stiffness. Paragraph J.15 is devoted to two other examples of fiber subjected to a series of force steps and tested by 4 kHz oscillations.

The sigmoid of Fig 6a essentially displays a three-phase look: constancy if 0.8 ≤ pT0_i_ ≤ 1, increasing linearity if 0.2 ≤ pT0_i_ ≤ 0.8 then again constancy if 0 ≤ pT0_i_ ≤ 0.2.

## Discussion

### Isometric tetanus plateau and determination of T0

#### Uniform density around an average angular position (θ_0_)

In each hs of an isometrically tetanized fiber, the angle θ which characterizes the orientation of the lever belonging to a WS head is distributed uniformly between θ_T_ and θ_up_ over the rangeδθ_T_ equal to about 50° (Fig 2). It is possible to interpret this uniform law as an angular dispersion equal to ± δθ_T_/2 around the average position θ_0_ defined in (4). For fibers isolated from *tibialis anterior* muscle of *Rana Esculenta* and *Rana Temporaria*, we find with the data from Table 1 of Paper 3:

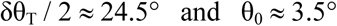

These values are in accordance with those of the literature: Fig 2 in [18]; Fig 3 and Table 1 in [19] ; Fig 2c in [20]; “*around 20°*” with Fig 4A in [21]; “*at least 17°*” in [15]; “*between 20° and 25°*” in [11]; Fig 4B in [22]; Fig 4B in [23].

#### Tension stability (T0) during the isometric tetanus plateau

Hypothesis 6 theorized with equation (3) which defines the motor-moment as an affine function of θ leads to the calculation of T0 formulated in (8). Composed of 5 constants, T0 is also a constant under identical experimental conditions. Once T0 is reached, the fiber is in stable equilibrium characterized by an isometric tetanus plateau [10]. If the equilibrium is disturbed by a length step, the tension becomes equal to T0 again after a few tens of milliseconds: all hs regain the characteristic uniform density between θ_T_ and θ_up_, because between θ_T_ and θ_down_ no head is found in WS, this absence being explained by the slow detachment of the myosin heads on δθ_E_ during the rise to the tetanus plateau (see accompanying Paper 1).

#### Analogy with the Buffon needle

In the 18th century, Georges-Louis Leclerc, still called Count of Buffon, cited as a naturalist and biologist, carried out various experiments where probability and geometry are interlocked. The most famous of his experiments is to calculate the universal constant π from random throws of a needle on a slatted parquet; read chapter 10 in [24]. The calculation of π result from the distribution of the angle of intersection of the needle with the groove in the parquet, angle uniformly spread over a semicircle after a considerable number of random throws. In a comparable way because of the abundant number of WS heads assigned to a hs of an isometrically tetanized fiber, the uniformity of θ between θ_up_ and θ_T_ leads to the calculation operated in equality (8) and to the constancy of T0.

#### Proportionality relationship between T0 and Λ_0_

The equality (8) indicates that T0 is proportional to Λ_0_, the number of WS heads per hs during the isometric tetanus plateau, result in accordance with observations [25,26,27,28]. This result is corroborated by the examples shown in Figs 6a and 6b as well as those shown in Figs J10 and J11 of Supplement S4.J.

#### Typology

With (8), T0 is also proportional to N_m_, the number of myofibrils. Thus, isometric tension must increase with the size of the muscle fibers and according to the typological classification of skeletal muscle fibers. We need to check:

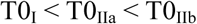

where T0_IIa_ and T0_IIb_ are the isometric tetanus tensions of type I, IIa and IIb fibers, respectively.

These inequalities are confirmed by various experimental measures [29,30,31,32].

### Minimum tension at the end of phase 1 of a length step (T1)

#### Presence of viscosity

For many physiologists, viscosity-induced forces are present and measured for elongations and shortenings on muscle fibers at rest, but when these same fibers are stimulated and shortened, the influence of viscosity disappears or is considered negligible [2,8,33,34]. On the contrary, the developments in Supplement S4.J reveal that viscosity forces contribute significantly to the drop in tension observed at the end of phase 1 of a length step. This difference in interpretation is explained: the hs of a muscle fiber is usually represented by a model with several elastic and viscous components distributed between the disks M and Z constituting the hs; see the Maxwell and Voigt rheological devices presented in Fig 35 in [2], Fig 5 in [35] or Fig 1 in [34]. The equations attached to these schemes are based on the relative velocities between the two disks; in the context of an isolated hs, the effects of viscosity are effectively negligible; see parameter ν defined in (15) whose order of magnitude is 10^−8^ nm^−1^. In our model, the viscous element is represented by a spring attached to the fixed end of the fiber (Figs J1 and J6 of Supplement S4.J), so the calculations are based on the absolute speeds of the disks M and Z. A hs cannot be isolated from the other hs. The myofibril must be studied in its entirety with all hs: on the one hand, the original elastic forces created by the WS heads are modelled by springs arranged in series and, on the other hand, the actions due to viscosity are modelled by springs arranged in parallel in a discretely progressive manner from hs to hs. From the calculations a multiplier coefficient (K) of the elastic slope (χ) emerges which, through equation (20), accords theory and observation. The values of K_z1_ and K_z2_ displayed in Table 1 for the examples studied are between 1.3 and 2, and are decisive for the evaluation of T1.

#### Adequacy of the model with the experimental data

The model equations correctly follow the experimental curves of pT1 as a function of 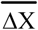; see figures in the article and supplement S4.J. The convex curvature of the slope present in the Mixed Zone (Fig 3A) is observable in many publications [1,2,5,7,33,36,37,38,39,40,41,42,43,44]. To our knowledge, our model is the first to provide an explanation. The imputation of viscosity forces in the equations implies that the shortenings of the hs of a myofibril are no longer equal. Shortening is increased from the proximal hs, i.e. close to the fixed end of the fiber, to the distal hs, i.e. adjacent to the mobile end of the fiber (Figs J1 and J4). With equations (20) and (21), the change between the two slopes (K_z1_·χ_z1_) and (K_z2_·χ_z2_) relating to the Zone 1 True and Zone True2, respectively, is done gradually in the Mixed Zone where the shortenings of the distal hs are in Zone 2 while those of the proximal hs are still in Zone 1.

Further evidence of the increasing variability of hs shortenings characterized at the end of phase 1 of a length step is given in Fig 5 in [8] where the stiffness of the segment that is measured with proximal hs near the transducer is greater than the stiffness of the fiber evaluated from hs located in the center of the myofibril..

#### Influence of the duration of phase 1 (τ_p1_)

Paragraph J.7 of Supplement S4.J is devoted to the role of τ_p1_ and the model indicates that viscosity forces decrease when τ_p1_ increases (and vice versa) in accordance with observations; see Fig 4B of [45]. The study in Fig J3 highlights the problem relating to the value of τ_p1_: either τ_p1_ is short, i.e. less than 0.2 ms as recommended, and viscosity forces are strongly present, or τ_p1_ is greater than 0.2 ms and the role of viscosity decreases but the rapid initiation of myosin heads specific to phase 2 begins during phase 1 and disrupts the interpretation.

#### Influence of the value of the intermediate or instantaneous tetanus tension (T0_i_)

Paragraph J.11 of the supplement S4.J is responsible for the theoretical developments of this topic, the main deductions from which are summarized in the Methods section. The experimental works brought to our attention in which T0_i_ is modified are explained and predicted by the model. The Results section on this theme and paragraphs J12 to J15 illustrate the robustness of the equations. Once again, the role of viscosity is essential. We note that the decrease in T0_i_ induces that of q_i_ via the expression (27), and increases exponentially K_i_ according to equality (26). This fact explains why the presence of viscosity is so pronounced when the fiber is at rest (T0_i_ ≈ 0).

#### Analytical “Nanoscope”

From equation (24) are deduced the instant or intermediate formulations of the normalized stiffness (e_i_/e _0_) and the strain (Y_i_) as a function of pT0_i_. These expressions lead to the calculation of the intermediate angular range (δθ_T,i_) in (31a) and (31b). This results in a “nanoscope” that allows analysis of the intermediate uniform density of the θ angle between the two equilibrium states, the total relaxation (T0_i_=0 ; δθ_T,i_=0) and the isometric tetanus plateau (T0_i_=T0; δθ_T,i_=δθ_T_). The evolution of δθ_T,i_ is formulated in four ways: (1) Exponential rise from 0 to δθ_T_ representing a variation in intracellular concentration, the presence of a cross-bridge disruptor or inhibitor (Inset within Fig J11b); (2) Exponential descent from δθ_Max_ to δθ_T_ characteristic of the tension rise from rest to the isometric tetanus plateau (Figs 5a, J12a, J12b, J12c and J13a); (3) Sigmoidal descent from δθ_T_ to 0 specific to the relaxation phase following the end of the stimulation.(Fig 5a) ; (4) Sigmoidal rise from δθ_T_ to δθ_Max_ emblematic of phase 4 of a series of force steps leading to the F/V relationship (Fig 6a, Insets within Figs J14a and J14b); in support of the description given in Fig 14 from [12], this sigmoidal rise corroborates the explanation given in the Discussion section of Paper 1 to “Why the maximum step appeared shorter for high tensions? “. This result will be confirmed again with the calculations carried out with accompanying Paper 6.

The 4 evolutions of δθ_T,i_ are empirically explained with the equations given from (J76) to (J78). These expressions, which are inspired by Pierre François Verhulst’s logistics laws, must be able to be demonstrated.

### Study of the slack-test as a conclusion

The significant variability in the hs shortenings observed at the end of phase 1 of a large length step 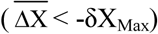 explains the slack-test results. The larger the step, the greater the velocity of shortening and the more influential the viscosity becomes with an exponential increase in the K coefficient associated with a decrease in the viscous parameter q (Fig J3). This particular case is illustrated in Fig J4 for q=1.5, a case in which the differences between shortening lengths are very marked depending on the position of the hs within the myofibril.

At the end of phase 1, i.e. t=0 according to Fig 1, the hs are divided into 3 categories.

1/ Distal hs: the shortening of these hs is greater in modulus than δX_Max_, and all the myosin heads that were in WS before the step are now detached while new heads are about to slowly initiate a WS (see Paper 5); the tension applied to the edges of each hs is zero and is at the origin of the slack of the fiber.

2/ Proximal hs: as the shortening of these hs is nil or almost nil, there is no change and all the myosin heads initially in WS remain operational at the end of phase 1; the tension at the terminals of each hs is always equal to T/N_m_ where T is the fiber tension before the step and where N_m_ is the number of myofibrils.

3/ Central hs: the shortening of these hs is either in Zone 1 or Zone 2; the tension at the endpoints of these hs varies gradually from T/N_m_ to 0.

After phase 1 of a slack-test step, the fiber returns to static equilibrium with rapid disappearance of viscosity (< 0.5 ms) and rebalancing of the length of shortenings around the average value 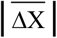.

We project a series of thought experiences.

1/The isometric tetanus plateau is reached with an average sarcomere length of 2.5 µm. Then the fiber is shortened with a constant velocity (u_Max_) which corresponds to the maximum speed observed during phase 4 of a force step. As an example, we choose the value presented in Fig 2 of Paper 1, that is in modulus:

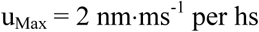

As a first approximation, the uniform density of θ for all hs of the fiber extends to δθ_Max_ with pT0_i_ close to zero as shown in Fig 6a.

When the isotonic ramp performed at u_Max_ reaches amplitude of 300 nm for an average sarcomere length of 2.2 µm, a large length step with 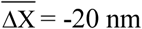 is applied.

After the end of phase 1, the distal and central hs are rebalanced in tension and length. For proximal hs that have not shortened, the uniform density of θ is at t=0 identical to that before the step, i.e. δθ_Max_. In a similar way to phase 4 of a force step corresponding to a very low tension near zero, the proximal hs shorten after the end of phase 1 to the u_Max_ velocity over the entire length 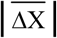 during the τ_slack_ duration such that:

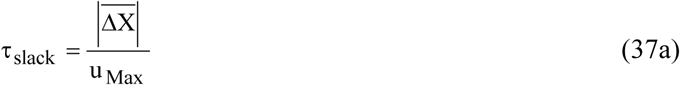

Once the proximal hs are at the right length at time t = τ_slack_, the rebalancing is completed and the fiber tension increases from 0, meaning the end of the slack.

This process is repeated for 3 other length steps with 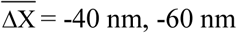 and −80 nm. The comments are similar and the relationship (37a) remains valid for the other 3 steps.

The relationship 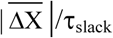 established in (37a) is represented in Fig 7 by a red line of u_Max_ slope passing through the origin and by the 4 red circles symbolizing the 4 length steps experienced by thought.

**Fig 7.**
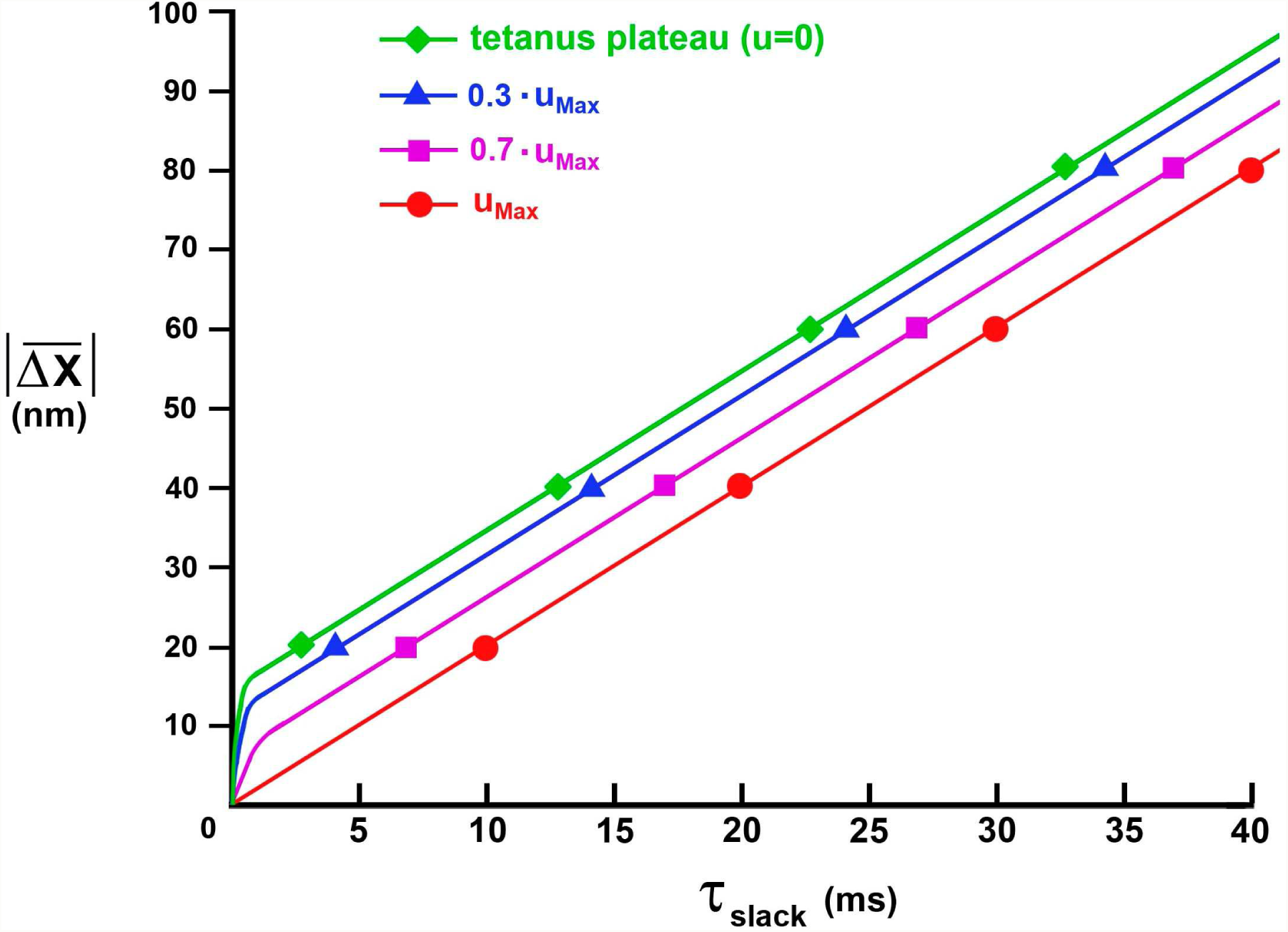
Theoretical relationship between the step length 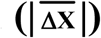 and the duration of the fiber slack (τ_slack_) according to the value of the shortening velocity of the isotonic ramp preceding the length step.

2/ The experiment is repeated by choosing a lower shortening velocity (0.7 u_Max_) for the isotonic ramp. After isometric tetanization where the average length of the sarcomeres is 2.5 µm, the fiber is shortened at a constant velocity equal to (0.7 u_Max_). As a first approximation, the uniform density of θ for all hs of the fiber extends over an angular range δθ_T,0.7_ smaller than δθ_Max_ according to Fig 6a.

Then once the amplitude of the isotonic ramp has reached 300 nm, a large length step of −20 nm is applied. For proximal hs that have not shortened, the uniform density of θ is unchanged and remains equal to δθ_T,0.7_. Since the tension exerted at the edges of these hs is greater than that of the previous case, the proximal hs are shortened to a speed (u0_.7_) greater than u_Max_ over a maximum distance equal to the stroke size of a myosin head (δX_Max_). Then the new heads slowly initiate a WS, the distribution of θ in the proximal hs extends again uniformly on δθ_Max_ and the shortening velocity becomes u_Max_ again over the remaining distance 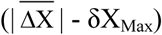.

To take into account this faster start on δX_Max_ compared to the previous case, a time delay (τ_0.7_) is introduced such that:

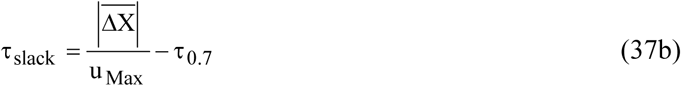

with

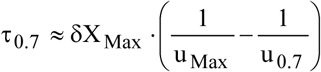

Once the proximal hs are at the correct length at time t = τ_slack_, the rebalancing is completed and the fiber tension increases from zero, meaning the end of the slack.

This process is repeated for 3 other length steps with 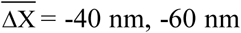 or −80 nm. The comments are similar: at the end of phase 1, the uniform density of θ is equal to δθ_T,0.7_ and the proximal hs shorten with the speed u_0.7_ on δX_Max_. Then the shortening speed becomes u_Max_ again on 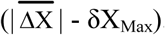. The expression (37b) remains valid with the same time lag τ_0.7_ for each of the other 3 steps.

The relationship relation 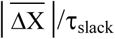 established in (37b) is represented in Fig 7 by a purple line of u_Max_ slope passing through 4 purple squares symbolizing the 4 length steps. For the same step value, the purple square is located to the left of the red point and the time difference between each purple square and each red point is equal to τ_0.7_.

3/ The experiment is repeated by choosing a slower velocity (0.3 u_Max_) for the isotonic ramp. Since the tension exerted at the endpoints of these hs is higher than in the previous case, the proximal hs are shortened to a velocity (u_0.3_) greater than u_0.7_ over the range δX_Max_. Then the new heads slowly initiate a WS and the shortening speed becomes u_Max_ again over the remaining distance to be covered 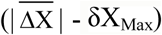. To take into account this faster start on δX_Max_, a time lag (τ_0.3_ > τ_0.7_) is introduced:

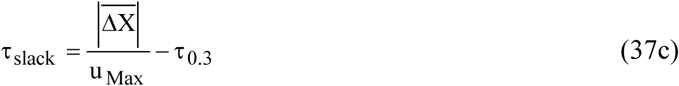

with

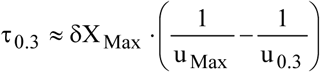

This process is repeated for 3 other length steps with 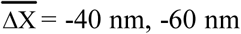 or −80 nm. The comments are similar and the expression (37c) remains valid with the same time delay τ_0.3_ for each of the other 3 steps.

The relationship 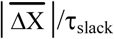 established in (37c) is represented in Fig 7 by a blue line of u_Max_ slope passing through 4 blue triangles symbolizing the 4 length steps. For the same step value, the blue triangle is located to the left of the purple square, the time difference between each blue triangle and each red dot is equal to τ_0.3_.

4/ The experiment is concluded by choosing a zero shortening velocity characteristic of the isometric tetanus plateau where the tension exerted at the edges of each hs is maximum (T0/N_m_). Just after the end of phase 1, the proximal hs shorten to a speed (u_0_) greater than u_0.3_ on δX_Max_ then the shortening speed becomes u_Max_ again over the remaining distance 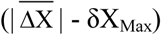.

To take into account this faster start on δX_Max_, a time delay (τ_0_ > τ_0.3_) is introduced:

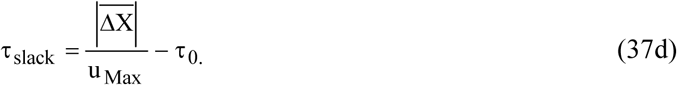

with

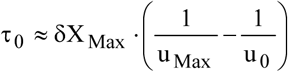

The relationship 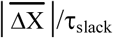 established in (37d) is represented in Fig 7 by a green line of u_Max_ slope passing through 4 green diamonds symbolizing the 4 steps of length. For the same step value, the green diamond is located to the left of the blue triangle and the time difference between each green diamond and each red dot is equal to τ_0_.

It so happens that this series of experiments was carried out on a frog fiber; see description associated with Fig 3 in [46]. The relationship 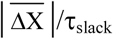 that is explored for 5 isotonic ramp velocities and 4 length steps is displayed in Fig 4A from [46] and is, with the exception of a few details, the carbon copy of Fig 4A. For information, with the values collected in Fig 4A from [46], the orders of magnitude relative to τ_0_ and u_0_ are respectively 5.5 ms and 20 nm·ms^−1^.

K. A. P. Edman was the first to perform the slack-test experiment [47] and to indicate that the slope of the straight line connecting the points of the relationship 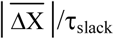 was equal to u_Max_.

The explanation given to the experimental results of the slack-test provides new demonstration of the preeminent role of viscosity in rapid shortening. The same is true for fast stretching.

## Supporting information

Supplementary Chapter. Calculations for T0 and T1Elas

Data used for computer programs used for Paper 4

Computer programs for Paper 4

Supplementary Chapter. Calculation of T1 with its two elastic and viscous components

## Acknowledgements

For this article,

I thank Professor V. Lombardi for allowing me to reproduce the points presented in Fig 4.

I thank Professor M. Irving for allowing me to reproduce the points presented in Figs 5 and 6.

For the Supplement S4.J,

I thank Professor M. Linari for allowing me to reproduce the points presented in Fig J11.

I thank Professor G. Cecchi for allowing me to reproduce the points presented in Fig J12.

I thank Professor G. Piazzesi for allowing me to reproduce the points presented in Fig J13.

I thank Professor K.A.P. Edman for allowing me to reproduce the points presented in Fig J14.

## Supporting information

### S4.I Supplementary Chapter. Calculations for T0 and T1Elas

I.1 Preamble: relationship of homogeneity between discrete and continuous uniform laws

I.2 Cases where the only actions involved are the forces and moments of inter-segmentary links

I.3 Isometric cases

I.4 Linear relationship between the displacement of the right half-sarcomere (hsR) and the rotation of the lever belonging to a WS head

I.5 Calculation of pT1_Elas_, the relative tension applied at the end of phase 1 of a length step in the absence of viscosity

I.6 Redefining of Zones 1 and 2

References of Supplement S4.I

### S4.J Supplementary Chapter. Calculation of T1 with its two elastic and viscous components

J.1 Viscosity forces: general

J.2 Viscosity forces during shortening (ΔL<0) or lengthening (ΔL>0) of a muscle fiber

J.3 Calculation of the tension at the end of phase 1 in the presence of viscosity forces when all hs shortenings belong to Zone 1

J.4 Calculation of the tension at the end of phase 1 in the presence of viscosity forces when all hs shortenings belong to Zone 2

J.5 Calculation of the tension at the end of phase 1 in the presence of viscosity when all elongations and shortenings of hs belong to Zone O

J.6 Application with study of an example

J.7 Explicit relationship of the tension at the end of phase 1 after a length step in Zone 1 Enlarged

J. 8 Forces induced by the presence of viscosity

J. 9 Influence of the duration of phase 1 (τ_p1_)

J. 10 Influence of tetanus plateau tension under isometric conditions (T0)

J.11 Influence of instantaneous or intermediate tetanus tension (T0_i_)

J.12 Study of instant strain (Y_i_) and instant angular range (δθ_T,i_) with an example

J.13 Study of the instant linear range (δX_T,i_) and instant normalized stiffness (e_i_/e0) during the rise to the tetanus plateau according to 3 models

J.14 Study of the instant linear range (δX_T,i_), the instant stiffness referred to T0_c_ (e_i_/T0_c_) and the instant strain (Y_i_) during the tension rise to the isometric tetanus plateau

J.15 Study of the intermediate normalized stiffness (e_i_/e0) during phase 4 of a series of force steps where the tension and the velocity of shortening are constant

J.16 Factors affecting viscosity forces

References of Supplement S4.J

**C4 Supplementary Material. Computer programs for calculating and plotting the relationships 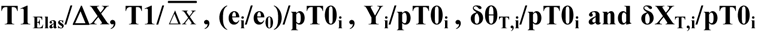**. Algorithms are written in Visual Basic 6.

**D4 Supplementary Material. Data used for computer programs used for Paper 4.** Access Tables are transferred to Excel sheets.

