## Supplementary Chapter. Calculations for T0 and T1Elas for "Mechanical model of muscle contraction. 4. Theoretical calculations of the tension during the isometric tetanus plateau and the tension exerted at the end of phase 1 of a length step"

### S4.I Supplementary Chapter of Paper 4: Calculations of T0 and $T1_{Elas}$

#### I 1 Preamble: homogeneity relation between discrete and continuous Uniform laws

Hypothesis 4 in Paper 2 indicates that the lever (S1b) of a myosin head in working stroke (WS) moves in a fixed plane. The orientation of S1b in this fixed plane is given by the angle θ having as bounds $\theta_{up}$ and $\theta_{down}$ relative to the *up* and *down* positions; the values of $\theta_{up}$ and $\theta_{down}$ are determined in (12a) and (12b) in Paper 2.

In a half-sarcomere (hs) located on the right, we consider the two angles $\theta_1$ and $\theta_2$ as follows:

$$\theta_{down} \le \theta_2 < \theta_1 \le \theta_{up}$$

The random variable Θ is associated with the angle θ; see definition of a random variable in paragraph A.1 of Supplement S1.A to accompanying Paper 1.

The discrete Uniform law of the discrete random variable Θ consists of Λ real values ($\theta_k$) spaced by the same distance on the interval [$\theta_2$ ; $\theta_1$]. The probability ($a_k$) associated with each of the Λ values $\theta_k$ verifies:

$$a_k = \frac{1}{\Lambda} \tag{I1}$$

The continuous Uniform law ($\mathcal{U}$) of the continuous random variable Θ on the interval [$\theta_2$ ; $\theta_1$] is formulated:

$$\mathcal{U}(\theta) = \frac{1}{\delta\theta} \cdot \mathbf{1}_{[\theta_2;\theta_1]}(\theta) \tag{I2}$$

where $\delta\theta = |\theta_2 - \theta_1|$ ; $\mathbf{1}$ is the indicator function defined in (A2b) in Supplement S1.A of Paper 1.

We study two uniform distributions of Θ under identical conditions in a hs on the right.

$\Lambda_A$ is an integer number of heads in WS. The discrete random variable Θ follows a discrete Uniform law according to (I1) and the $\Lambda_A$ values of θ are spaced the same distance ($\delta\theta_k$) over the interval [$\theta_{A2}$;$\theta_{A1}$] such that:

$$\delta\theta_A = (\theta_{A2} - \theta_{A1})$$

with $\theta_{down} \le \theta_{A2} \le \theta \le \theta_{A1} \le \theta_{up}$

$\Lambda_B$ is an integer number of heads in WS. The discrete random variable Θ follows a discrete Uniform law according to (I1) and the $\Lambda_B$ values of θ are spaced the same distance ($\delta\theta_k$) over the interval [$\theta_{B2}$;$\theta_{B1}$] such that:

$$\delta\theta_B = (\theta_{B2} - \theta_{B1})$$

with $\theta_{down} \le \theta_{B2} \le \theta \le \theta_{B1} \le \theta_{up}$

We assume that $\Lambda_A$ and $\Lambda_B$ are large enough for the transition from discrete to continuous to be possible and that the random variable $\Theta$ is distributed uniformly on a continuous basis on $\delta\theta_A$ and $\delta\theta_B$, respectively. Thus, depending on whether one considers $\Theta$ as a discrete or continuous random variable, by intrinsic homogeneity of the discrete and continuous Uniform laws, one checks according to (I1) and (I2):

$$\frac{\Lambda_A}{\Lambda_B} = \frac{\delta\theta_A}{\delta\theta_B} \tag{I3}$$

By symmetry, the equality (I3) is valid in a hs located on the left.

#### I.2 Case where the only actions involved are the linking forces and moments

An instantaneous tension (T) is applied to an isolated fiber in isometry or shortening at steady velocity (V). The case where the only actions involved are the linking forces and moments is studied in paragraph F.3 of Supplement S2.F to accompanying Paper 2. It is demonstrated that at any given time, each hs of the fiber has an equal number of WS heads, an identical distribution of the angle $\theta$, and an equality of tensions ($T_{hs}$) acting on both ends of the hs alternately delimited by a Z-disk and a M-disk (Figure F1a). Tension $T_{hs}$ is calculated in modulus from equation (F14) of Supplement S2.F, reproduced below:

$$T_{hs} = T_m = \frac{T}{N_m} = \frac{1}{L_{S1b} \cdot S_{WS}} \cdot \sum_{b=1}^{\Lambda_L} \mathcal{M}^{(b)} \tag{I4}$$

where $T_m$ is the tension in modulus applied to the 2 ends of each of the $N_m$ myofibrils (Fig F1a of Supplement S2.F); $N_m$ is the number of myofibrils of the fiber; $L_{S1b}$ is the length of the lever (S1b); $S_{WS}$ is a constant characteristic of the myosin head defined in the equations (13) and (14) of Paper 2; $\Lambda_L$ is the common instantaneous number of WS myosin heads per hs; b is the index of a WS head; $\mathcal{M}^{(b)}$ is the module of the instantaneous motor-moment applied to the lever of WS head n° b.

$\overline{\mathcal{M}_L}$ is the module of the instantaneous average moment of the $\Lambda_L$ motor-moments, equal by definition to:

$$\overline{\mathcal{M}_L} = \frac{\sum_{b=1}^{\Lambda_L} \mathcal{M}^{(b)}}{\Lambda_L} \tag{I5}$$

The combination of (I4) and (I5) provides:

$$T_{hs} = \frac{\Lambda_L}{L_{S1b} \cdot S_{WS}} \cdot \overline{\mathcal{M}_L} \tag{I6}$$

**I.3 Isometric case**

In each hs of a fiber stimulated under isometric conditions, hypothesis 5 enacted in accompanying Paper 3 states that the angle θ of the levers belonging to the WS heads is distributed over the interval $[\theta_2 ; \theta_1]$ according to the same Uniform law ($\mathcal{U}_L$ ):

$$\mathcal{U}_L(\theta) = \frac{1}{\delta\theta_L} \cdot \mathbf{1}_{[\theta_2;\theta_1]}(\theta) \tag{I7}$$

where $\delta\theta_L$ is the angular range equal to:

$$\delta\theta_L = |\theta_2 - \theta_1| \tag{I8}$$

and where $\theta_1$ and $\theta_2$ are two angles that in a hs on the right check the following inequalities:

$$\theta_{down} \leq \theta_2 \leq \theta \leq \theta_1 \leq \theta_{up} \tag{I9}$$

The equality (16) of Paper 3 introduces the angle $\theta_T$ (Fig I1) and the angular range $\delta\theta_T$ equal to about 50°, observed during the isometric tetanus plateau. The spatial density of θ has been postulated as uniform on $\delta\theta_T$ by various authors [1,2,3,4,5]. The geometric study developed in Paper 3 leads to a similar resolution formulated in (19) and reproduced here:

$$\mathcal{U}_T(\theta) = \frac{1}{\delta\theta_T} . \mathbf{1}_{[\theta_T;\theta_{up}]}(\theta) \tag{I10}$$

The density $\mathcal{U}_T$ is represented in Fig I2a by a green rectangle of width $\delta\theta_T$ and height $(1/\delta\theta_T)$.

During the isometric tetanus plateau, the common number of WS myosin heads per hs is $\Lambda_0$ and the $\Lambda_0$ values of θ are spaced the same distance ($\delta\theta_0$) over the $\delta\theta_T$ range.

The experimental conditions being identical to those leading to the isometric tetanus plateau, we consider the common number of WS myosin heads per hs ($\Lambda_L$) such that the $\Lambda_L$ values of θ are spaced the same distance ($\delta\theta_0$) over the $\delta\theta_L$ range defined with (I8) and (I9). We assume that $\Lambda_L$ is important enough to consider the transition from discrete to continuous, i.e. the random variable Θ associated with the angle θ is interpreted indifferently as a discrete or continuous random variable. With relations (I7), (I8) and (I10), the formula (I3) becomes:

$$\Lambda_L = \Lambda_0 \cdot \frac{\delta\theta_L}{\delta\theta_T} \tag{I11}$$

Hypothesis 6 postulates that the motor-moment ($\mathcal{M}$) is an affine function of θ given in equation (3) of Paper 4 and reproduced below:

$$\mathcal{M}(\theta) = \mathcal{M}_{up} \cdot \frac{(\theta - \theta_{down})}{\delta\theta_{Max}} . \mathbf{1}_{[\theta_{down};\theta_{up}]}(\theta) \tag{I12}$$

where $\mathcal{M}_{up}$ is the maximum moment relating to *up* position; $\delta\theta_{Max} = |\theta_{up} - \theta_{down}|$.

By definition of the average of a variable ($\mathcal{M}$) that is a bijective function of another variable ($\theta$) associated with a continuous random variable ($\Theta$) distributed according to (I7), the calculation of the average moment ($\overline{\mathcal{M}_L}$) is written with (I9):

$$\overline{\mathcal{M}_L} = \int_{\theta_2}^{\theta_1} \mathcal{M}(\theta) \cdot \mathcal{V}_L(\theta) \cdot d\theta \tag{I13}$$

The integration of (I13) with (I12) gives:

$$\overline{\mathcal{M}_L} = \frac{\mathcal{M}_{up} \cdot \left[\frac{\theta_1 + \theta_2}{2} - \theta_{down}\right]}{\delta\theta_{Max}} \tag{I14}$$

When the angle $\theta$ is distributed uniformly between the 2 angles $\theta_1$ and $\theta_2$, discretely or continuously in each hs of an isometrically stimulated fiber, the calculation of $T_{hs}$ is formulated after rewriting the relationship (I6) by juxtaposing the expressions (I5) and (I14):

$$T_{hs} = \left(\frac{\Lambda_L \cdot \mathcal{M}_{up}}{L_{S1b} \cdot S_{WS} \cdot \delta\theta_{Max}}\right) \cdot \left[\frac{\theta_1 + \theta_2}{2} - \theta_{down}\right] \tag{I15}$$

The calculation of $T_{hs}$ is performed with angular positions taken in a hs on the right. By symmetry we obtain an identical result in a hs on the left, hence the generic index "hs".

##### *I.3.1 Application to the isometric tetanus plateau*

During the isometric tetanus plateau, the terminals $\theta_1$ and $\theta_2$ are $\theta_{up}$ and $\theta_T$ and we check:

$$\delta\theta_L = \delta\theta_T \tag{I16a}$$

$$\Lambda_L = \Lambda_0 \tag{I16b}$$

The angle $\theta_0$ is located at the middle of $\delta\theta_T$ (Fig I1) such that:

$$\theta_0 = \frac{\theta_{up} + \theta_T}{2} \tag{I17}$$

$T0_{hs}$ is the tension at the ends of any hs during the isometric tetanus plateau. The calculation of $T0_{hs}$ is done with equation (I15) combined with equations (I16a), (I16b) and (I17):

$$T0_{hs} = \left(\frac{\Lambda_0 \cdot \mathcal{M}_{up}}{L_{S1b} \cdot S_{WS} \cdot \delta\theta_{Max}}\right) \cdot [\theta_0 - \theta_{down}] \tag{I18}$$

##### *I.3.2 Relative tension of a fiber stimulated in isometric conditions*

The relative value of $T_{hs}$ to $T0_{hs}$ ($pT_{hs}$) is deduced from the equations (I15) and (I18) associated with the relationship (I11):

$$pT_{hs} = \frac{T_{hs}}{T0_{hs}} = \left(\frac{\delta\theta_L}{\delta\theta_T}\right) \cdot \frac{\left[\frac{\theta_1 + \theta_2}{2} - \theta_{down}\right]}{(\theta_0 - \theta_{down})} \qquad \text{(I19)}$$

T0 is the tension at the extremities of the fiber during the isometric tetanus plateau. According to the equality (I4), the relative tension of the fiber with respect to T0 (pT) is:

$$pT = \frac{T}{T0} = \frac{N_m \cdot T_m}{N_m \cdot T0_m} = \frac{T_{hs}}{T0_{hs}} = pT_{hs} \qquad \text{(I20)}$$

Tensions at the endpoints of the muscle fiber, myofibrils and all hs are equal when expressed in relation to the tensions of the isometric tetanus plateau referring to each of these elements.

#### I.4 Linear relationship between the displacement of the half-sarcomere and the rotation of the lever belonging to a WS head

To characterize the hs displacement along the longitudinal axis of the myofibril, it is resorted to the abscissa X of point A representing the site of attachment of the WS head to the actin filament relatively to point D representing the ball joint between the myosin filament and the rod (Fig D2b of Supplement S2.D to the accompanying Paper 2). By arbitrarily matching the zero of the abscissa X with the angle $\theta_0$ defined in (I17) and after integrating equation (15) of Paper 2, the relationship between X and θ is formulated:

$$X = L_{S1b} \cdot R_{WS} \cdot (\theta - \theta0) \,.\, \mathbf{1}_{[\theta_{down}\,;\theta_{up}]}(\theta) \qquad \text{(I21a)}$$

where $S_{WS}$ is a constant characteristic of the myosin head defined in the equations (13) and (14) of Paper 2; $S_{WS}$ is equal to $R_{WS}$.

Conversely θ is expressed as a function of X according to:

$$\theta = \left(\theta_0 + \frac{X}{L_{S1b} \cdot R_{WS}}\right) .\, \mathbf{1}_{[\theta_{down}\,;\theta_{up}]}(\theta) \qquad \text{(I21b)}$$

The relationships (I21a) and (I21b) imply affine scale correspondence between X and θ in the linear domain framed by $\theta_{down}$ and $\theta_{up}$ (Fig I1).

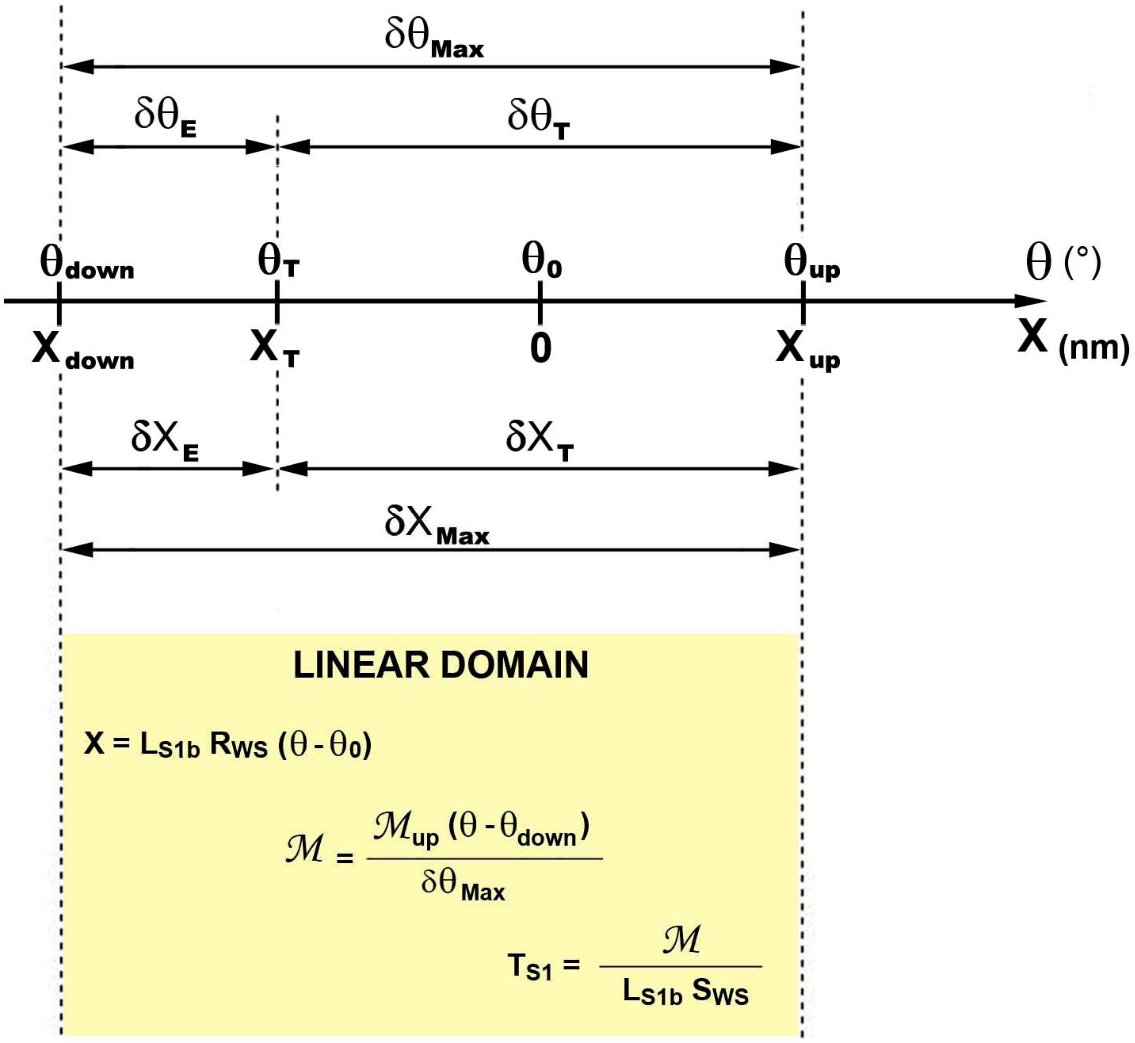


**Fig I1. Affine relation in the linear domain defined by $\theta_{up}$ and $\theta_{down}$ between the θ angle of lever S1b and the relative position X of the binding-site on the actin molecule with respect to the myosin filament along the longitudinal axis of a half-sarcomere on the right.**

From the above, the following expressions are deducted:

$$\delta X_T = (X_{up} - X_T) = L_{S1b} \cdot R_{WS} \cdot \delta\theta_T \qquad \text{(I22)}$$

$$\delta X_E = (X_T - X_{down}) = L_{S1b} \cdot R_{WS} \cdot \delta\theta_E \qquad \text{(I23)}$$

$$\delta X_{Max} = (X_{up} - X_{down}) = (\delta X_T + \delta X_E) = L_{S1b} \cdot R_{WS} \cdot \delta\theta_{Max} \qquad \text{(I24)}$$

$$X_{up} = -X_T = \frac{\delta X_T}{2} \qquad \text{(I25)}$$

$$X_{down} = (X_{up} - \delta X_{Max}) = \left(\frac{\delta X_T}{2} - \delta X_{Max}\right) = -\left(\frac{\delta X_{Max} + \delta X_E}{2}\right) \qquad \text{(I26)}$$

$$\frac{1}{|X_{down}|} = \frac{1}{\left(\delta X_{Max} - \frac{\delta X_T}{2}\right)} = \frac{1}{L_{S1b} \cdot R_{WS} \cdot \left(\delta\theta_{Max} - \frac{\delta\theta_T}{2}\right)} \qquad \text{(I27)}$$

The importance of the relationship (I27) will thereafter be noted.

##### *I.4.1 Application to the isometric tetanus plateau*

The tension $T0_{hs}$ is exerted by the $\Lambda_0$ WS heads on both sides of a hs during the isometric tetanus plateau. The expression (I18) is rewritten with the transformation of variables between θ and X provided by (I21b):

$$T0_{hs} = \left(\frac{\Lambda_0 \cdot \mathcal{M}_{up}}{L_{S1b} \cdot S_{WS}}\right) \cdot \frac{|X_{down}|}{\delta X_{Max}} \qquad \text{(I28a)}$$

With (I27), $T0_{hs}$ is also formulated according to $\delta X_{Max}$ and $\delta X_T$:

$$T0_{hs} = \left(\frac{\Lambda_0 \cdot \mathcal{M}_{up}}{L_{S1b} \cdot S_{WS}}\right) \cdot \left(1 - \frac{\delta X_T}{2 \cdot \delta X_{Max}}\right) \qquad \text{(I28b)}$$

With (I4), the tension of the isometric tetanus plateau is:

$$T0 = N_m \cdot \left(\frac{\Lambda_0 \cdot \mathcal{M}_{up}}{L_{S1b} \cdot S_{WS}}\right) \cdot \left(1 - \frac{\delta X_T}{2 \cdot \delta X_{Max}}\right) \qquad \text{(I29)}$$

##### *I.4.2 Application to a fiber stimulated in isometric conditions*

The 2 abscissa $X_1$ and $X_2$ correspond to the 2 angles $\theta_1$ and $\theta_2$ defined in (I9). With (I21a), they are equal to:

$$X_1 = L_{S1b} \cdot R_{WS} \cdot (\theta_0 - \theta_1)$$

$$X_2 = L_{S1b} \cdot R_{WS} \cdot (\theta_0 - \theta_2)$$

The linear range ($\delta X_L$) between $X_1$ and $X_2$ is calculated using (I8):

$$\delta X_L = |X_1 - X_2| = L_{S1b} \cdot R_{WS} \cdot \delta\theta_L \qquad \text{(I30)}$$

When $\theta$ is uniformly distributed between $\theta_1$ and $\theta_2$, the tension ($T_{hs}$) exerted by the $\Lambda$ myosin heads in WS on both sides of a hs belonging to an isometrically stimulated fiber is rewritten according to (I15) and (I21b):

$$T_{hs} = \left( \frac{\Lambda_L \cdot \mathcal{M}_{up}}{L_{S1b} \cdot S_{WS} \cdot \delta X_{Max}} \right) \cdot \left[ \frac{X_1 + X_2}{2} + |X_{down}| \right] \qquad \text{(I31a)}$$

The equalities (I19) and (I20) become:

$$pT = pT_{hs} = \left( \frac{\delta X_L}{\delta X_T} \right) \cdot \left[ 1 + \frac{(X_1 + X_2)}{2 \cdot |X_{down}|} \right] \qquad \text{(I31b)}$$

#### I. 5 Calculation of the tension at the end of phase 1 of a length step in the absence of viscosity ($pT1_{Elas}$)

During the isometric tetanus plateau, the angle $\theta$ is associated with the random variable $\Theta$ which follows the Uniform law $\mathcal{U}_T$, defined in (I10) and represented by a green rectangle in Fig I2a. The relative tension (pT0) associated with this uniform density is equal to 1 (Fig I2a'). The fiber is then quickly shortened by a length step ($\Delta L$) at constant velocity (V) during $\tau_{p1}$, the time of phase 1. Any hs is shortened by a length step ($\Delta X$) at constant speed (u) during $\tau_{p1}$. The $\Lambda_0$ levers S1b belonging to the $\Lambda_0$ WS heads all undergo the same angular variation ($\Delta\theta$) according to the linear relationship (I21b) as:

$$\Delta\theta = \frac{\Delta X}{L_{S1b} \cdot R_{WS}} \qquad \text{(I32)}$$

At the end of phase 1, the relative tension exerted on both edges of the hs is called as $pT1_{Elas}$.

We assume, as many authors do [3,5,6], that no new head has time to initiate a WS during phase 1 since the average rapid initiation duration is about 1 ms [3,7,8,9,10,11,12], a time much greater than 0.2 ms, the recommended duration for $\tau_{p1}$ [13].

**There are two zones for the calculation of $pT1_{Elas}$**

***I.5.1 Calculation of $pT1_{Elas}$ in Zone 1 ≡ [ $-\delta X_E$ ; 0 ]***

The range $\delta X_E$ is shown in Fig I1. The hs shortening ($\Delta X$) checks for inequalities:

$$-\delta X_E \leq \Delta X < 0 \qquad \text{(I33a)}$$

With (I21a) and (I23), the corresponding inequalities for the rotation $\Delta\theta$ of the $\Lambda_0$ S1b are:

$$-\delta\theta_E \leq \Delta\theta < 0 \qquad \text{(I33b)}$$

In Zone 1, the $\Lambda_0$ WS heads at the beginning of phase 1 are still in WS at the end of phase 1 since the $\theta$ angles of the $\Lambda_0$ levers are all between $\theta_{down}$ and $\theta_{up}$. Switching from discrete to continuous, the angle $\theta$ is associated with $\Theta$, the continuous random variable uniformly distributed over the interval [$\theta_T$+$\Delta\theta$ ; $\theta_{up}$+$\Delta\theta$] represented by a green rectangle in Fig I2b. The Uniform law ($\mathcal{U}_{z1}$) is stated:

$$\mathcal{U}_{z1}(\theta) = \frac{1}{\delta\theta_T} \cdot \mathbf{1}_{[\theta_T + \Delta\theta ; \theta_{up} + \Delta\theta]}(\theta) \qquad \text{(I34)}$$

The two abscissa $X_1$ and $X_2$ relative to the two angles ($\theta_{up}$+$\Delta\theta$) and ($\theta_T$+$\Delta\theta$) are calculated with (I21a) and (I32):

$$X_1 = X_{up} + \Delta X \qquad \text{(I35a)}$$

$$X_2 = X_T + \Delta X \qquad \text{(I35b)}$$

In support of (I30) and (I22), we deduce:

$$\delta X_L = \delta X_T \qquad \text{(I36)}$$

Thanks to (I25), we obtain:

$$X_1 + X_2 = 2 \cdot \Delta X \qquad \text{(I37)}$$

At the end of phase 1 of the length step, the velocity is cancelled and the fiber is again in isometry. At this precise time, the conditions are met to apply (I31b), and the relative tension in Zone 1 ($pT1_{Elas,z1}$) is formulated using (I36) and (I37):

$$pT1_{Elas,z1} = 1 + \frac{\Delta X}{|X_{down}|} \qquad \text{(I38)}$$

In isometric conditions, the tensions exerted on each side of the hs of the fiber are all equal in modulus. From this assertion we deduce with (I38) that the shortening $\Delta X$ is a constant and it is concluded that all hs are shortened by the same variation in length belonging to Zone 1 and equal to:

$$\Delta X = \frac{\Delta L}{N_{hs}} \qquad \text{(I39)}$$

where $N_{hs}$ is the number of hs per myofibril.

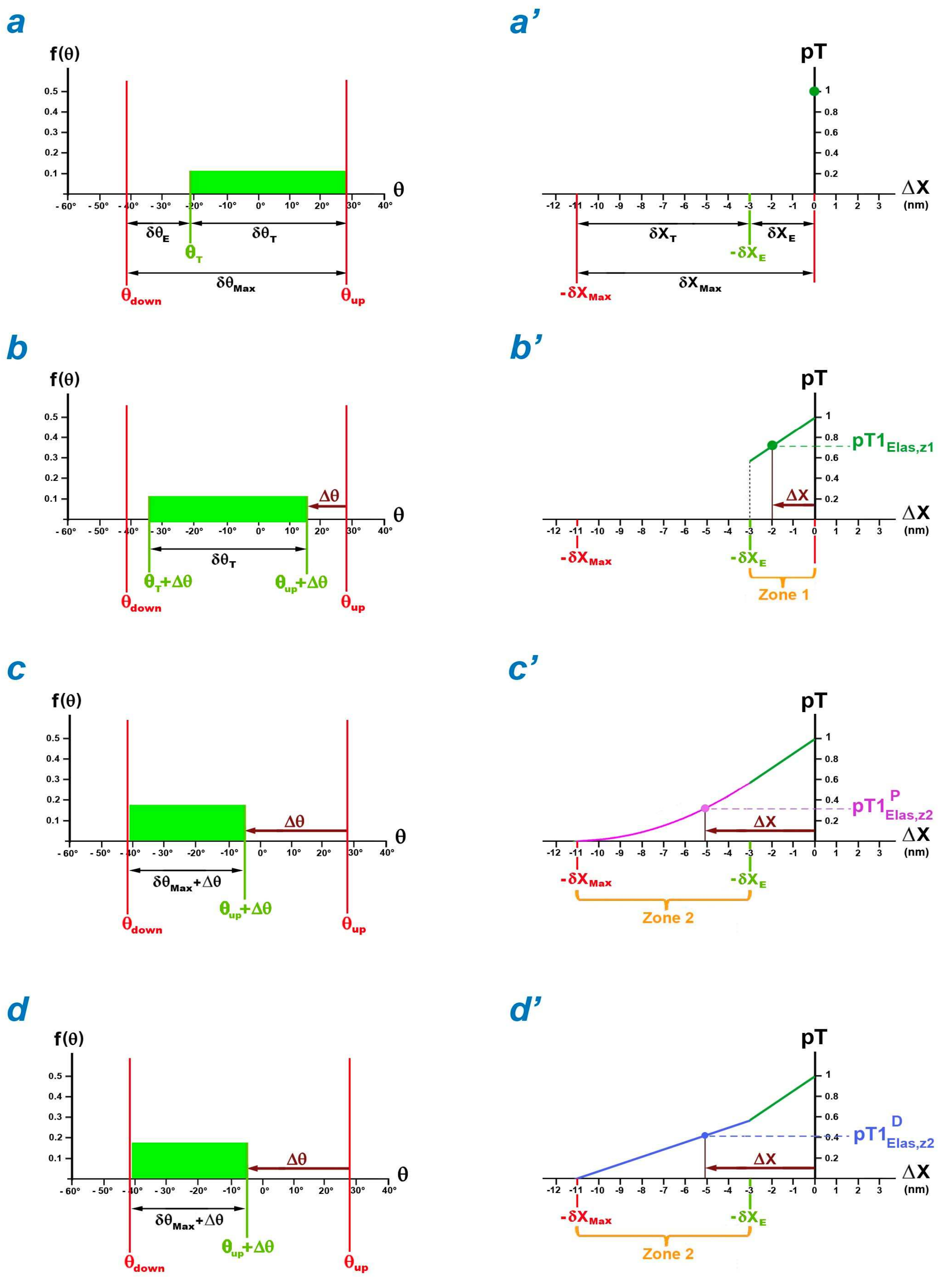


**Fig I2. Uniform distributions of θ in a hs on the right at the origin of the calculations of $pT1_{Elas}$ as a function of the hs length step (ΔX).**

(a) Uniform density of θ between $\theta_T$ and $\theta_{up}$ for $\Lambda_0$ WS heads, characteristic of the isometric tetanus plateau. (b) Uniform density of θ between $(\theta_T + \Delta\theta)$ and $(\theta_{up}+\Delta\theta)$ after the rotation Δθ of the $\Lambda_0$ levers. (c) and (d) Uniform densities of θ between $\theta_{down}$ and $(\theta_{up}+\Delta\theta)$ after the rotation Δθ of the $\Lambda_{WS}$ and $\Lambda_0$ levers, respectively. (a') Unit relative tension. (b') Line segment representing the relative tension $(pT1_{Elas,z1})$ as a function of ΔX between $-\delta X_E$ and 0. (c') and (d') Parabolic arc and straight line segment representing, respectively, the relative tensions $pT1_{Elas,z2}{}^P$ and $pT1_{Elas,z2}{}^S$ as a function of ΔX between $-\delta X_{Max}$ and $-\delta X_E$.

In Zone 1, the affine function (I38) is represented by a dark green straight line segment in Fig I2b'. The slope ($\chi_{z1}$) is based on (I27):

$$\chi_{z1} = \frac{1}{|X_{down}|} = \frac{1}{L_{S1b} \cdot R_{WS} \cdot \left(\delta\theta_{Max} - \frac{\delta\theta_T}{2}\right)} \qquad \text{(I40)}$$

The slope $\chi_{z1}$ depends on several geometric parameters (see Table 1 of Paper 2) characteristic of the myosin II head, i.e. $L_{S1b}$, $\delta\theta_{Max}$, $\delta\theta_T$, $R_{WS}$. The last term $R_{WS}$ itself depends on the geometric parameters $L_{S2}$, $d_{AMfil}$, $X_{AB}$ and $Y_{AB}$.

##### *I.5.2 First calculation of $pT1_{Elas}$ in Zone 2 ≡ [ $-\delta X_{Max}$ ; $-\delta X_E$ [*

The hs shortening ($\Delta X$) checks for inequalities:

$$-\delta X_{Max} \leq \Delta X < -\delta X_E < 0 \qquad \text{(I41a)}$$

From (I21a) and (I24) result the corresponding inequalities for the rotation $\Delta\theta$ of the $\Lambda_0$ levers:

$$-\delta\theta_{Max} \leq \Delta\theta < -\delta\theta_E < 0 \qquad \text{(I41b)}$$

$\Lambda_{WS}$ is the number of heads still in WS at the end of phase 1 among the initial $\Lambda_0$ heads, i.e. the $\Lambda_{WS}$ heads whose $\theta$ angle of their lever verifies the condition:

$$\theta_{up} \leq (\theta + \Delta\theta) \qquad \text{(I42)}$$

The ($\Lambda_0$-$\Lambda_{WS}$) heads whose $\theta$ angle of their lever does not achieve the condition (I42) have a zero contribution to the tension in accordance with (I12).

The $\theta$ angle of the $\Lambda_{WS}$ levers S1b is associated with the random variable $\Theta$ which follows a discrete uniform law on the interval [$\theta_{down}$ ; $\theta_{up}$+$\Delta\theta$] according to hypothesis 5 of uniformity. By passing to continuous, the angle $\theta$ is associated with $\Theta$, a continuous random variable following the Uniform law ($\mathcal{U}_{z2}$) which is formulated:

$$\mathcal{U}_{z2}(\theta) = \frac{1}{(\delta\theta_{Max} + \Delta\theta)} \cdot \mathbb{1}_{[\theta_{down}\,;\,\theta_{up}+\Delta\theta]}(\theta) \qquad \text{(I43)}$$

The $\mathcal{U}_{z2}$ law is represented in Fig I2c by a green rectangle of width ($\delta\theta_{Max}$+$\Delta\theta$) and height 1/($\delta\theta_{Max}$+$\Delta\theta$).

The two abscissa $X_1$ and $X_2$ relative to the two angles ($\theta_{up}$+$\Delta\theta$) and $\theta_{Tdown}$ are calculated with (I21a) and (I32):

$$X_1 = X_{up} + \Delta X \qquad \text{(I44a)}$$

$$X_2 = X_{down} \qquad \text{(I44b)}$$

The sum gives:

$$X_1 + X_2 = X_{down} + X_{up} + \Delta X \quad \text{(I45)}$$

It is deduced with (I24) and (I30):

$$\delta X_L = |X1 - X2| = \delta X_{Max} + \Delta X \quad \text{(I46)}$$

At the end of phase 1 of the length step, the shortening velocity is cancelled and the fiber is again in isometry. At this precise time, the conditions are met to apply (I31b) and the relative tension in Zone 2 ( $pT1^{P}_{Elas,z2}$ ; P for Parabola) is formulated by introducing (I45) and (I46):

$$pT1^{P}_{Elas,z2} = \frac{(\delta X_{Max} + \Delta X)^2}{2 \cdot \delta X_T \cdot |X_{down}|} \quad \text{(I47)}$$

The expression (I47) is represented by a parabolic arch (Fig I2c'; purple line).

In isometric conditions, the tensions exerted on each side of the hs of the fiber are all equal in modulus. From this assertion we deduce with (I47) that the hs shortening ($\Delta X$) is a constant and it is concluded that all hs are shortened by the same variation in length belonging to Zone 2 according to equality (I39).

##### *I.5.3 Second calculation of $pT1_{Elas}$ in Zone 2 ≡ [ $-\delta X_{Max}$ ; $-\delta X_E$ [*

When the lever is in the *down* position, a landmark event at the end of the working stroke, the next step in the chemical-mechanical cycle of the cross-bridge [14,15] is the detachment of a myosin head that takes effect over a period of a few milliseconds. This event named as {FastDE} is presented in paragraph B.5 and discussed in paragraph B.6 of Supplement S1.B to accompanying Paper 1. The occurrence delay of {FastDE} is 1 ms and the time constant of {FastDE} is between 3 and 5 ms (Table B1), which is much longer than $\tau_{p1}$. Subsequently, at the end of phase 1, the $\Lambda_0$ WS heads are still strongly bound to the actin filament, including the ($\Lambda_0$-$\Lambda_{WS}$) heads whose lever is assumed to have reached and exceeded the *down* position. The ($\Lambda_0$-$\Lambda_{WS}$) heads initially in WS are likely to contribute to the tension. We conjecture an extreme situation where all $\Lambda_0$ heads would be in WS at the end of phase 1 and where the angle $\theta$ of the $\Lambda_0$ levers is associated with $\Theta$, a random variable distributed uniformly over the interval [$\theta_{down}$ ; $\theta_{up}$+$\Delta\theta$] according to $\mathcal{U}_{z2}$ defined in (I43); see Fig I2d identical to Fig I2c. The expressions (I44a), (I44b), (I45) and (I46) remain valid but the homogeneity relationship stated in (I11) is no longer verified. It is necessary to use the formulation (I31a) which brings with (I16b), (I24) and (I45) a new expression of the tension en Zone 2 ( $T1^{S}_{hs,z2}$ ; S for Straight line):

$$T1^{S}_{hs,z2} = \left( \frac{\Lambda_0 \cdot \mathcal{M}_{up}}{L_{S1b} \cdot S_{WS}} \right) \cdot \left[ \frac{\delta X_{Max} + \Delta X}{2 \cdot \delta X_{Max}} \right] \quad \text{(I48)}$$

The relative tension in Zone 2 ($pT1^{S}_{Elas,z2}$) is calculated by dividing $T1^{S}_{hs,z2}$ with $T0_{hs}$ whose calculation is provided in (I28a) as follows:

$$pT1^{S}_{Elas,z2} = \frac{1}{2 \cdot |X_{down}|} \cdot (\delta X_{Max} + \Delta X) \quad \text{(I49)}$$

In Zone 2, the affine function (I49) is a straight line segment (Fig I2d'; blue line). The slope ($\chi_{z2}$) is:

$$\chi_{z2} = \frac{1}{2 \cdot |X_{down}|} = \frac{\chi_{z1}}{2} \quad \text{(I50)}$$

That is half of the one found for Zone 1.

#### I.6 Redefining of Zones 1 and 2

Depending on the conditions of realization of phase 1, in particular the duration $\tau_{p1}$, and the internal geometry of the myofibril, in particular the inter-filament space, the value of the real tension in Zone 2 lies between the 2 extreme cases, $pT1^{P}_{Elas,z2}$ and $pT1^{S}_{Elas,z2}$, determined in (I47) and (I49), respectively.

We propose a compromise modeling between these 2 borderline cases (Fig I3).
The linear abscissa $-\delta X_{z1}$ is a parameter depending on the experimental conditions, defined as:

$$-2 \cdot \delta X_E \leq -\delta X_{z1} \leq -\delta X_E \quad \text{(I51)}$$

**Zones 1 and 2 are redefined in relation to the abscissa $-\delta X_{Z1}$.**
**Zone 1 ≡ [ -δXz1 ; 0 [**

The relationship between pT1 and ΔX determined in (I38) remains valid:

$$pT1_{Elas,z1} = 1 + \chi_{z1} \cdot \Delta X \quad \text{(I52a)}$$

Equality (I52a) is a straight line segment extended to $-\delta X_{z1}$ and represented by a dark green straight line in Fig I3. The slope ($\chi_{z1}$) is formulated in (I40) and reproduced below:

$$\chi_{z1} = \frac{1}{|X_{down}|} \quad \text{(I52b)}$$

2/ **Zone 2 ≡ [ $-\delta X_{Max}$ ; $-\delta X_{z1}$ [**

The relation between pT1 and ΔX is represented by a straight line segment that connects the coordinate point ($\Delta X = -\delta X_{z1}$ ; $pT1 = 1 - \chi_{z1} \cdot \delta X_{z1}$) to the coordinate point ($\Delta X = -\delta X_{Max}$ ; pT1=0):

$$pT1_{Elas,z2} = \chi_{z2} \cdot (\delta X_{Max} + \Delta X) \quad \text{(I53a)}$$

The slope ($\chi_{z2}$) is equal to:

$$\chi_{z2} = \frac{(1 - \chi_{z1} \cdot \delta X_{z1})}{\delta X_{Max} - \delta X_{z1}} \quad \text{(I53b)}$$

The straight line segment appears as light green in Fig I3.
In the particular case where $\delta X_{z1} = \delta X_E$, the relationship (I50) is found using (I26) and (I27).

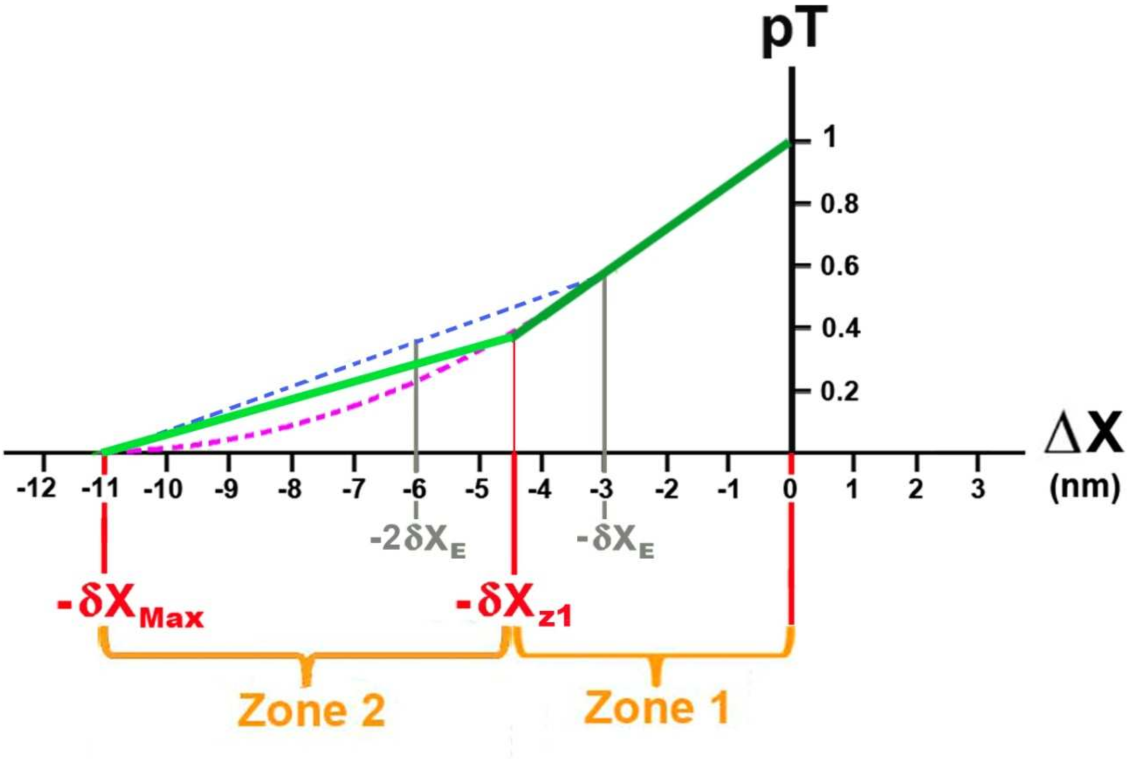


**Fig I3. Redefinition of Zones 1 and 2 with the abscissa -δ$X_{z1}$ between -2·δ$X_E$ and -δ$X_E$.**

The relative tension (pT1) calculated according to ΔX in (I52a) and (I52b) is represented by a straight line segment colored dark green for Zone 1 and light green for Zone 2. The two extreme cases $pT1^{P}_{Elas,z2}$ and $pT1^{S}_{Elas,z2}$, determined in (I47) and (I49), are represented in pink and blue dotted lines, respectively.
