## Supplementary material for "Mechanical model of muscle contraction. 4. Theoretical calculations of the tension during the isometric tetanus plateau and the tension exerted at the end of phase 1 of a length step": Computer programs for Paper 4

### CP4 Computer programs relative to Paper 4

#### CP4.1 Calculation and curve of T1 as a function of ΔX

The curve on the computer screen is done by calling the "SUB AAPH_ARTNEG_trace()" routine.

Once the data have been recovered (See values displayed in the Tables of Paper 4 and in the Excel sheets of Supplement DA4), the experimental pT1 tensions from published articles and the pT1 theoetrical curves from model equations are plotted as a function of the hs shortening (Fig CP4.1).

A regression is performed between experimental and theoretical tensions (see Methods section).

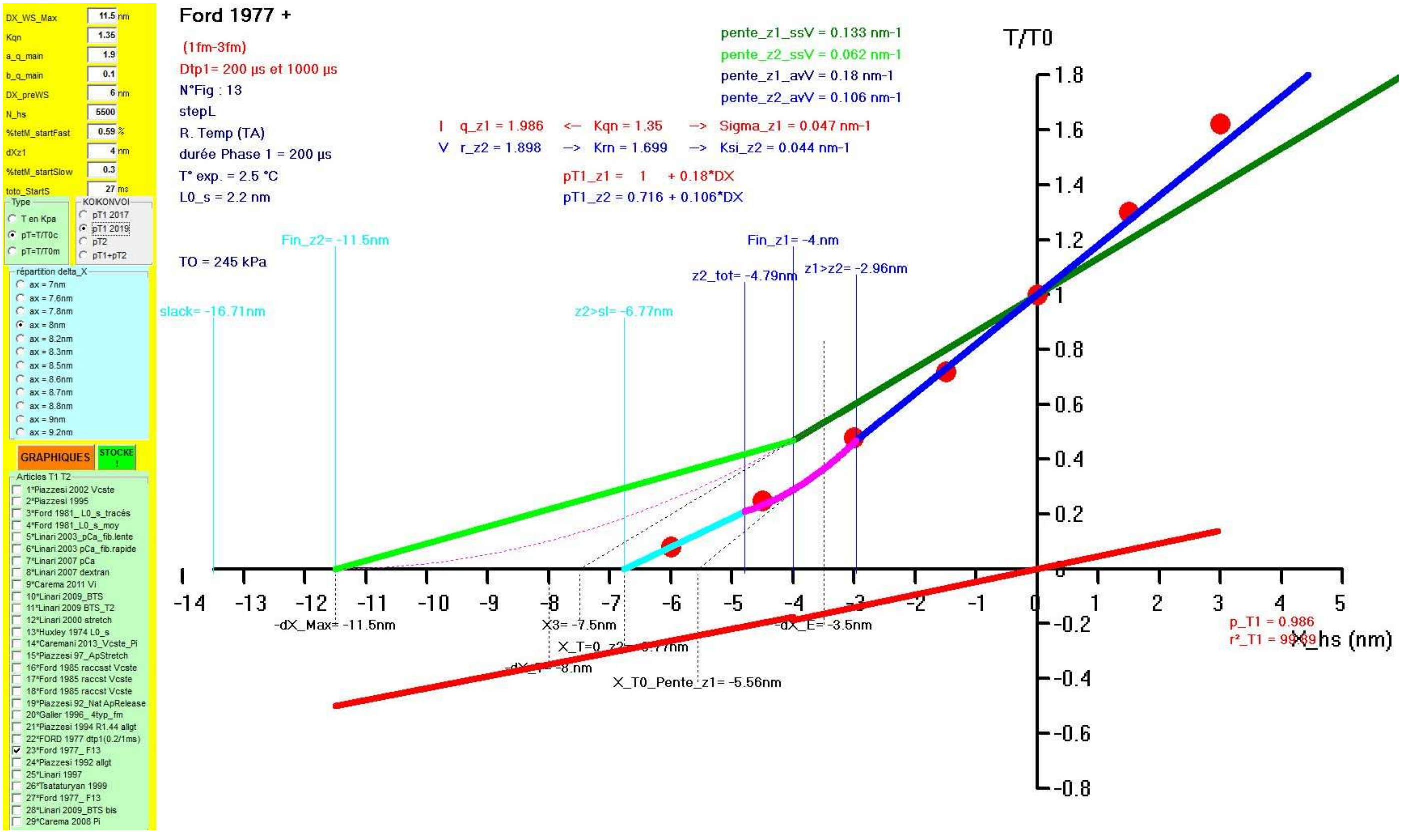


**Fig CP4.1 Screenshot after starting the the "Sub AAPH_ARTNEG_trace()" routine.**

### Sub AAPH_ARTNEG_trace()

```
Dim choix_coul As Integer, droites_VIsco As Integer, t_arti As Integer

droites_VIsco = 1 'on voit les droites viscosité
'droites_VIsco = 0 'par défaut

If Left(ressort.Tag, 4) <> "ph12" Then Stop
If mmm = 1 Then
  O.AutoRedraw = True: O.Cls 'verifier que fonction autoredrax=tue
  FrT2n.Visible = 0

'prepa pour prsentation tracés article
  With Frpres 'pres= presentaion 0="TOUT" 1="ART +DR_Visc +DR_Elast" 2="ART -DR_Visc +DR_Elast" 3="ART +DR_Visc -DR_Elast" 4="ART -DR_Visc -DR_Elast")
      .Top = Height * 0.35: .Left = Width * 0.9
      .Visible = -1
   End With
  For t = 0 To 4 '6
     With Opres(t)
       .Visible = -1
       .Caption = Choose(t + 1, "TOUT", "ART +DR_Visc +DR_Elast", "ART -DR_Visc +DR_Elast", "ART +DR_Visc -DR_Elast", "ART -DR_Visc -DR_Elast")
      End With
   Next

   FrTemp.Visible = 0
End If

Np4 = 100
e_pt = 25 * IIf(mmm = 1, 1, 3)
e_tt = 4 * IIf(mmm = 1, 1, 3)
L_WS = Val(TAM(0).Text) '    dX_Max
aX = OaX(FraX.Tag).Tag * 2 ' dX_T      T pour Tétanos T
Xmin = aX - L_WS '           -dX_E <0  E pour End

X1 = aX / 2 '               Xup = dX_T/2 >0
X2 = -X1                 '    X_T = -Xup   <0
X3 = X1 - L_WS '            Xdown = - dX_T/2-dX_E = -(dW_Max+dX_E)/2

chi_p1 = 1 / Abs(X3) 'pente ss Vis Zone 1   = 2/(DXmin + DXmax)
KqnZ1 = Val(TAM(1).Text) '0.1 '* 2 '0.25 - chi_p1 ' 0.033 * 3 'viscosité_MetZ_disjoints

a_q = Val(TAM(2).Text) 'pour Varaiation T / temp /rigor
b_q = Val(TAM(3).Text)

L_preWS = Val(TAM(4).Text)
N_hs = Val(TAM(5).Text) '5000
P_startF = Val(TAM(6).Text) 'coeff remontee max
dXz1 = Val(TAM(7).Text) 'pT Xa1 pour linéarisation de la parabolle zonze2 amorti
P_startS = Val(TAM(8).Text) 'coeff remontee max
to1_startS = Val(TAM(9).Text)

If dXz1 > 2 * Abs(Xmin) Then
  vd = MsgBox("Revoir valeur Xz1")
  TAM(7).Text = Format(Abs(Xmin))
  Exit Sub
End If
If Fresc.Tag = 0 Then '2016
   'chi_p2 = chi_p1 / 2 'pente ss Vis Zone 2    = 1/(DXmin + DXmax)
   X_Z1 = Xmin
ElseIf Fresc.Tag = 1 Then '2019
    X_Z1 = -dXz1
```

```
   'chi_p2 = (1 - chi_p1 * dXz1) / (L_WS - dXz1)
End If

chi_p2 = (1 + X_Z1 * chi_p1) / (L_WS + X_Z1)

'-------qZ1------------KqnZ1 = N_hs ^ (1 - qZ1 / 2) * coth[N_hs ^ (1 - qZ1 / 2)]
calcul_qoK (KqnZ1)
qZ1 = qOK
xmi = N_hs ^ (1 - qZ1 / 2)
cH_n = (Exp(xmi) + Exp(-xmi)) / 2
sH_n = (Exp(xmi) - Exp(-xmi)) / 2
z1_min = X_Z1 / KqnZ1
z1_max = z1_min * cH_n
Sigma_Vi_z1 = (KqnZ1 - 1) * chi_p1

'-------rZ2------------
rZ2 = qZ1 - Log(chi_p1 / chi_p2) / Log(N_hs)
xmi = N_hs ^ (1 - rZ2 / 2)
cH_n = (Exp(xmi) + Exp(-xmi)) / 2
sH_n = (Exp(xmi) - Exp(-xmi)) / 2
KrnZ2 = calcul_Kqn(rZ2)
z2_min = -L_WS / KrnZ2
z2_max = z2_min * cH_n
Ksi_Vi_z2 = (KrnZ2 - 1) * chi_p2
X_T0 = z2_min

vx = "X_hs (nm)"
vx = "X_hs (nm)"
xn = -L_WS - IIf(Fresc.Tag < 2, 2, 4)
xx = 5
xt = 1

'T0c=250'Kpa
If Frtyp.Tag = 1 Then
  impscale X0, Y0 + 130, X0 + larg, Y0 - 10 '14
  vy = "T(kPa)": yn = 0: yx = 500: yt = 50
Else
   impscale X0, Y0 + 130, X0 + larg, Y0 + 10 '14
   T0c = 1
   vy = "T/T0"
   yn = IIf(droites_VIsco = 1, -0.8, -0.4) 'pour voir viscosité = -0.4 sino par défaut
   yx = 1.8: yt = 0.2
End If

O.DrawWidth = e_tt
impenvi_Xneg

    '                fresc.tag=0        1          2      2
    '.Caption = Choose(t + 1, "pT1 ", "pT1 tout", "pT2", "pT1+pT2")

 '--  trace ds points enregistrés ds table_T1T2
   O.DrawMode = 13
   N_fm = 0 'compteur de tracées

   For t_arti = 0 To N_articles - 1
       If ChT1T2(t_arti) = 1 Then 'c'est ok
       'en 1 pour temp etudie ensemble a partir du T0 de 0℃
        z = ChT1T2(t_arti).Tag
        N_fm = N_fm + 1: zfm = z
        'les complments
        T1T2met.Seek "=", z
        If T1T2met.NoMatch = 0 Then
```

```
     AAPH_ARTNEG_Recalcul 'on part de T1c et de T2 puis on recacule tout ds pT1c et pT2
    If Frtyp.Tag = 0 Then T0c = T0(0)
    If IsNull(T1T2met("methode")) = 0 Then
      Select Case T1T2met("methode")
      Case "variation_T0":  AAPH_ARTNEG_variation_T0 ("T0") 'qi = a + b *log(pT0i)
       Case "variation_tp1" 'on passe
      Case "rigor_vite_D":  AAPH_ARTNEG_variation_T0 ("Rigor") 'AAPH_ARTNEG_RIGOR_D 'D pour Droites avec stepL
release et stretch rapides (< 0.2 ms)
      Case "rigor_stretch_lent_P":  AAPH_ARTNEG_RIGOR_P 'P pour Parabole avec stretching lent (> 1 sec ie > plusiers
millers de ms)"
      Case "variation_temp": AAPH_ARTNEG_variation_T0 ("temp") 'qi = a + b * pT0i ou exp ?
      Case Else: Stop
     End Select
    End If
   End If
 'titre

  T1T2.Seek "=", z, -99: If T1T2.NoMatch Then Stop
  'titre = titre + Trim(T1T2("fm")) + " + "

 'tracés des pts (surtout tout lasser ici) 'avec recalculs faits ds pT1c et pT2
  O.DrawWidth = e_tt
  O.DrawWidth = e_pt

  'couleur
  choix_coul = 1 'par defaut

  'on personnlaise
  Select Case zfm
   Case 22 'ford 77
     choix_coul = 2
      bcol(10) = &H80FF&   'orange
      bcol(1) = &HFF00FF  'mauve
   Case 1 'piazzesi 2002 +1995
    choix_coul = 2
    bcol(1) = QBColor(11) 'bleu ciel
    bcol(12) = &HFF00FF  'mauve
  End Select

  ' bcol(10) = &H80FF&   'orange
  'bcol(1) = QBColor(0) 'noir
   'bcol(0) = qbcolor(1) 'bleu noir
  ' bcol(1) = QBColor(11) 'bleu ciel
   'bcol(0) = &H808080    'gris
  ' bcol(2) = &HFF&    'rouge
  ' bcol(10) = &H80FF&   'orange
  ' bcol(12) = &HFF00FF  'mauve
  ' bcol(5) = &H8080&   'jaune foncé
  ' bcol(6) = &HFF&    'rouge

  T1T2.MoveNext
  Do While T1T2("cas") = z '10000 'z
   If Frpres.Tag <> 2 Then
    'pT1c ****
     If Fresc.Tag <> 2 And IsNull(T1T2("X1")) = 0 And IsNull(T1T2("pT1c")) = 0 Then
      xav = T1T2("X1"): yav = T1T2("pT1c") '/ 100
      xav = (xav - xn) * 100 / (xx - xn): yav = (yav - yn) * 100 / (yx - yn)
      If Fresc.Tag = 3 Then 'pT1+pT2
        O.PSet (xav, yav), QBColor(9)
      ElseIf choix_coul <= 2 Then 'classqi
        O.PSet (xav, yav), QBColor(T1T2("coul_pt"))
      Else
```

```
              coul = bcol(T1T2("coul_pt"))
              O.PSet (xav, yav), coul
             End If
           End If

           'pT2 ***
           If Fresc.Tag > 1 And IsNull(T1T2("X2")) = 0 And IsNull(T1T2("pT2")) = 0 Then
             'If z = 1 Or z = 3 Then coul = QBColor(T1T2("cod_t2"))
             xav = T1T2("X2"): yav = T1T2("pT2") ' / 100
             xav = (xav - xn) * 100 / (xx - xn): yav = (yav - yn) * 100 / (yx - yn)
             If choix_coul = 2 Then
               coul = bcol(T1T2("coul_pt2"))
               O.PSet (xav, yav), coul
             Else
               O.PSet (xav, yav), QBColor(T1T2("coul_pt")) 'special pour T2 QBColor(T1T2("coul_pt2")) por avoir les pts 1995 et 1992
             End If
           End If
         '*********************
         End If
         T1T2.MoveNext: If T1T2.EOF Then Exit Do
      Loop
    End If
  Next 'article


With O

   'Fresc.tag =  0= unisqt ss VISCosité//1= uniqt av VISCosité // 2= 2 ss et av VISCosité
   .DrawWidth = IIf(mmm = 1, 1, 3)
   .FontSize = 12


If N_fm = 1 And (Frpres.Tag = 0 Or Frpres.Tag = 3) Then 'frpres.tag donne le type de representation de la figure que l'on veur
(legende pas legende etc)

   T0c = IIf(Frtyp.Tag = 1, T0(0), 1)
   'legende valeurs en X
   xap = 5
   yap = 11


   For i = 0 To IIf(Fresc.Tag < 2, 11, 3)
       If i >= 6 And i <= 8 Then
         .ForeColor = QBColor(9)
      ElseIf i > 8 Then
         .ForeColor = QBColor(11)
      Else
        .ForeColor = QBColor(0)
      End If

      Select Case i
        Case 0: xav = X3: tx1 = "X3": z = 7 - xap 'trait L_WS
        Case 1: xav = Xmin: tx1 = "-dX_E": z = 7 - xap 'trait Xmin
        Case 2: xav = -L_WS: tx1 = "-dX_Max": z = 7 - xap 'trait L_WS
        Case 3: xav = -aX: tx1 = "-dX_T": z = 13 - xap 'trait ax
        Case 4: xav = X_T0: tx1 = "X_T=0_z2": z = 10 - xap 'trait ax
        Case 5: xav = -1 / (chi_p1 * KqnZ1): tx1 = "X_T0_Pente_z1": z = 15 - xap 'trait ax

       'limite vraies zone 1 et vraie zone 2
        Case 6: xav = X_Z1: tx1 = "Fin_z1": z = -40 - yap 'trait ax
        Case 7: xav = z1_min: tx1 = "z1>z2": z = -36 - yap 'trait ax
        Case 8: xav = z1_max: tx1 = "z2_tot": z = -35 - yap 'trait ax
```

```
     Case 9: xav = -L_WS: tx1 = "Fin_z2": z = -40 - yap 'trait ax
     Case 10: xav = z2_min: tx1 = "z2>sl": z = -30 - yap 'trait ax
     Case 11: xav = z2_max: tx1 = "slack": z = -30 - yap 'trait ax
   End Select
   tx1 = tx1 + "= " + Format(xav, "##.##") + "nm"
   If xav < xn Then xav = xn
   xav = (xav - xn) * 100 / (xx - xn)
   ymi = (-0.15 - yn) * 100 / (yx - yn)
   .DrawStyle = IIf(i < 6, 2, 0)
   O.Line (xav, ymi - z)-(xav, ymi + IIf(i = 1, 38, 5))
   .CurrentX = xav - .TextWidth(tx1) * 0.5
   .CurrentY = ymi - IIf(i < 6, z - 1, z - 2)
   O.Print tx1
 Next 'i

 'legende valeur en Y ss viscosité
   xav = Xmin: xav = (xav - xn) * 100 / (xx - xn)
   yav = T0c * (1 + xav * chi_p1)
   yap = (yav - yn) * 100 / (yx - yn)
   xap = (1 - xn) * 100 / (xx - xn)
   O.Line (xav, yap)-(xap, yap)
   .CurrentX = xap + 0.2
   .CurrentY = yap + 1
   O.Print "pT = " + Format(yav, "0.###")

 ' If Fresc.Tag < 2 Then 'legende valeur en Y avec viscosite
 '   xav = Xmin: yav = T0c * (1 + xav * (chi_p1 * KqnZ1))
 '   xav = (xav - xn) * 100 / (xx - xn): yap = (yav - yn) * 100 / (yx - yn)
 '   O.Line (xav, yap)-(xap, yap)
 '   .CurrentX = xap + 0.2
 '   .CurrentY = yap + 1
 '   O.Print "pT = " + Format(yav, "0.###")
 ' End If

'legende en haut à gauche

  For i = 1 To IIf(Fresc.Tag < 2, IIf(IsNull(T1T2met("methode")), 15, 12), 4)

   Select Case i
     Case 1: xav = chi_p1: tx1 = "pente_z1_ssV": k = 2           '1 color 2
     Case 2: xav = chi_p2: tx1 = "pente_z2_ssV": k = 10          '2 color 10
     Case 3: xav = chi_p1 * KqnZ1: tx1 = "pente_z1_avV": k = 1  '3 color 1
     Case 4: xav = chi_p2 * KrnZ2: tx1 = "pente_z2_avV": k = 9 '4 color 9
     Case 5: xav = qZ1: tx1 = "|    q_z1": k = 12               '5 color 12
     Case 6: xav = KqnZ1: tx1 = "<--   Kqn": k = 12             '6 color 12
     Case 7: xav = Sigma_Vi_z1: tx1 = "-->   Sigma_z1": k = 12   '7 color 12
     Case 8: xav = rZ2: tx1 = "V   r_z2": k = 9                '8  color 9
     Case 9: xav = KrnZ2: tx1 = "-->   Krn": k = 9             '9  color 9
     Case 10: xav = Ksi_Vi_z2: tx1 = "-->   Ksi_z2": k = 9      '10 color 9
     Case 11: xav = chi_p1 * N_hs ^ -qZ1 * 10000000: tx1 = "Nu_z1": k = 1 '11 color 1
     Case 12: xav = chi_p2 * N_hs ^ -rZ2 * 10000000: tx1 = "Nu_z2": k = 9 '12 color 9
     Case 13: xav = KqnZ1 * chi_p1: tx1 = "pT1_z1 =    1     + ": k = 12 '13 color 12
     Case 14: xav = chi_p2 * L_WS: tx1 = "pT1_z2": k = 9 '14 color 9
     Case 15: xav = KrnZ2 * chi_p2: tx1 = " + ": k = 9   '15 color 9
  End Select

   If i = 5 Or i = 8 Then 'q_z1 et r_z2
      O.CurrentX = 20
   ElseIf i = 6 Or i = 9 Or i = 13 Or i = 14 Then  'Kqn_z1 et Krn_z2 et Ksn_e3
      O.CurrentX = 31
   ElseIf i = 7 Or i = 10 Then 'Sigma_z1 et Ksi_z2 et Ksi_z3
```

```
      O.CurrentX = 42.2
    ElseIf i = 11 Or i = 12 Then 'nu
      O.CurrentX = 27
    ElseIf i = 15 Then  '
      O.CurrentX = 40
    Else
       O.CurrentX = 45
    End If

    If i >= 5 And i <= 7 Then 'Visc z1
      O.CurrentY = 112 - 6 * 3
    ElseIf i >= 8 And i <= 10 Then 'Vis_z2
      O.CurrentY = 112 - 7 * 3
   ElseIf i = 11 Or i = 12 Then  'nu_z1 et nu_z2
      O.CurrentY = 112 - (i - 13) * 3
    ElseIf i = 13 Then
      O.CurrentY = 87
    ElseIf i > 13 Then
      O.CurrentY = 84 '5
    Else
      O.CurrentY = 110 - i * 3
    End If
    O.ForeColor = QBColor(k)
    If i < 5 Or i = 7 Or i = 10 Then
     tx1 = tx1 + " = " + Format(xav, "0.###") + " nm-1"
    ElseIf i = 11 Or i = 12 Then
      tx1 = tx1 + " = " + Format(xav, "0.###") + " 10-7 nm-1"
    ElseIf i = 13 Or i = 15 Then
      tx1 = tx1 + Format(xav, "0.###") + "*DX"
    Else
     tx1 = tx1 + " = " + Format(xav, "0.###")
    End If
    O.Print tx1
   Next 'i

 End If
```

**AAPH_ART_legende**

```
'tracés droites de viscosité

  '0="TOUT" 1="ART +DR_Visc +DR_Elast" 2="ART -DR_Visc +DR_Elast" 3="ART +DR_Visc -DR_Elast" 4="ART -DR_Visc -
DR_Elast")



    '                fresc.tag=0        1          2      2
    '.Caption = Choose(t + 1, "pT1 ", "pT1 tout", "pT2", "pT1+pT2")
   drap = 0

    'tracés droites vetes elasticité
    '0="TOUT" 1="ART +DR_Visc +DR_Elast" 2="ART -DR_Visc +DR_Elast" 3="ART +DR_Visc -DR_Elast" 4="ART -DR_Visc -
DR_Elast")

    ''' AVEC droites vertes (ss Visc)
     For i = IIf(Frpres.Tag <= 2, 1, 4) To IIf(droites_VIsco = 1, 8, 6) '6

      Select Case i 'TT =trat plein  P=pointillé

       'SS viscosité
       Case 1 'TT zone 1 ss viscosité // pT=(1+0.15X)
          yav = 1.8 * T0c: xav = (1.8 - 1) / (chi_p1)
```

```
        xap = X_Z1: yap = T0c * (1 + xap * chi_p1) ' chi_p1 = Pente_ssV_Z1 = 1/abs(X3) = 0.15
        coul = QBColor(2) 'trait epais vert foncé

    Case 2 'P zone 1 ss viscosité// pT=(1+0.15X) : suite en pointille
         xav = X_Z1: yav = T0c * (1 + xav * chi_p1) ' il faut la terminer en pointille
         xap = X3: yap = 0
         coul = QBColor(0) 'pointillés noirs

    Case 3 'TT zone 2 ss viscosité // pT= 0.075(10+X)
        xav = X_Z1: yav = T0c * (1 + xav * chi_p1) '  chi_p2 = chi_p1 /2 = 1/ (2*abs(X3))
        xap = -L_WS: yap = 0
        coul = QBColor(10) 'trait epais vert clair

'AV viscosité
    Case 4 'TT zone 1 av viscosité // pT=(1+0.25X)
        xav = (1.8 - 1) / (chi_p1 * KqnZ1) '-z1_min ?
        yav = 1.8 * T0c
        xap = z1_min
        yap = T0c * (1 + xap * chi_p1 * KqnZ1) ' Pente_ssV_Z1 = chi_p1 + sigma_visc = 0.15 + 0.1 = 0.25

        If yap < 0 Then xap = -1 / (chi_p1 * KqnZ1): yap = 0: drap = 1
        If T1T2met("methode") = "variation_tp1" Then
           coul = &H80FF&    'orange QBColor(9) 'bleu
        ElseIf zfm = 1 Then
           coul = QBColor(13) 'bleu
        Else
           coul = QBColor(9) 'bleu
        End If

    Case 5 'P zone 1 av viscosité 'SUITE) // pT=(1+0.25X)
        'xav = Xmin 'avant
        xav = z1_min '2018
        yav = T0c * (1 + xav * chi_p1 * KqnZ1) ' il faut la terminer en pointille
        xap = -1 / (chi_p1 * KqnZ1): yap = 0
         coul = QBColor(0) 'pointillés noirs

    Case 6 'TT zone 2 av viscosité // pT= 0.175(4.3+X)
        'X_T0 = Xmin - (1 + (chi_p1 + sigma_visc) * Xmin) / (chi_p2 + sigma_visc) '-4.3nm
        'xav = Xmin 'avant
        xav = z1_max
        yav = T0c * (Abs(X_T0) + xav) * chi_p2 * KrnZ2
        xap = X_T0: yap = 0
        If T1T2met("methode") = "variation_T0" Then
           coul = QBColor(9) 'bleu
        ElseIf T1T2met("methode") = "variation_tp1" Then
           coul = &H80FF&    'orange QBColor(9) 'bleu
        Else
           coul = QBColor(11) 'bleu clair
        End If


     Case 7 'zone O que viscosité
         xav = -z1_min
         yav = T0c * chi_p1 * (KqnZ1 - 1) * xav

        'xap = Xmin 'avanr
        xap = X_Z1
        yap = T0c * chi_p1 * (KqnZ1 - 1) * xap
        coul = QBColor(12)

     Case 8 'zone 2 que viscosité
```

```
        xav = X_Z1
        yav = T0c * chi_p2 * (KrnZ2 - 1) * xav

        xap = -L_WS
        yap = T0c * chi_p2 * (KrnZ2 - 1) * xap
        coul = QBColor(12)


     End Select

     If xap < xav Then
         xav = (xav - xn) * 100 / (xx - xn): yav = (yav - yn) * 100 / (yx - yn)
         xap = (xap - xn) * 100 / (xx - xn): yap = (yap - yn) * 100 / (yx - yn)
        .DrawStyle = IIf(i = 2 Or i = 5, 2, 0)
        .DrawWidth = IIf(mmm = 1, 1, 3) * IIf(i = 2 Or i = 5, 1, 8)

         'coul = QBColor(Choose(i, 2, 0, 10, 9, 0, 11, 0))
         If Frpres.Tag = 0 Then
            O.Line (xav, yav)-(xap, yap), coul
         ElseIf Frpres.Tag = 1 And i = 5 Then 'il faut la terminer en pointille de la droite ZONE 1
         'ElseIf i = 5 Then 'si on veut le trait tout le temsp
             O.Line (xav, yav)-(xap, yap), coul
         ElseIf .DrawStyle = 0 And Fresc.Tag <> 2 Then
             O.Line (xav, yav)-(xap, yap), coul
         End If
     End If

     If drap = 1 Then Exit For
  Next

'**********LISSAGE entre ZONE 1 et ZONE2 2
 If Fresc.Tag <> 2 Then
     .DrawWidth = 8
      xav = z1_min
       yav = T0c * (1 + xav * chi_p1 * KqnZ1)
       O.CurrentX = (xav - xn) * 100 / (xx - xn)
       O.CurrentY = (yav - yn) * 100 / (yx - yn)
       If T1T2met("methode") = "variation_T0" Then
          coul = QBColor(9) 'on reste en bleu clair
      ElseIf T1T2met("methode") = "variation_tp1" Then
           coul = &H80FF&    'orange QBColor(9) 'bleu
       ElseIf Fresc.Tag = 3 Then 'pT1+pT2
         coul = QBColor(9)
       Else
         coul = QBColor(13) 'ou 5 ou QBColor(IIf(xmi < X_Z1, 11, 9))
       End If
       For t = 1 To Np4
           xmi = z1_min + t * (z1_max - z1_min) / Np4
           If xmi < X_T0 Then
              Exit For
           End If
           yav = T0c * (1 + xmi * chi_p1 * KqnZ1)
           yap = T0c * chi_p2 * (L_WS + KrnZ2 * xmi)
           poids = t / Np4
           ymi = (1 - poids) * yav + poids * yap
           If ymi < 0 Then Exit For
           xap = (xmi - xn) * 100 / (xx - xn)
           yap = (ymi - yn) * 100 / (yx - yn)
           O.Line -(xap, yap), coul
       Next
   End If
```

```
   '"'**ZONE parabole du mode Amorti interpolée linéairement
  '              fresc.tag=0         1          2      3
   '.Caption = Choose(t + 1, "pT12017 ", "pT12018", "pT2", "pT1+pT2")
   If Fresc.Tag = 1 And Frpres.Tag = 0 Then 'Or Fresc.Tag = 3
      .DrawStyle = 2
      .DrawWidth = 1
     'P tracé parabole en pointille: pT1=(chi_p2/ax0*(L_WS+DX)^2
      'coul = QBColor(IIf(i = 1, 10, 2)) 'vert clair // vert foncé
      xav = Xmin
      yav = (L_WS + xav) ^ 2 / (2 * Abs(X3) * aX) 'pT=(L_WS+X)^2/(2*ax*abs(X3))
      yav = T0c * yav
      .CurrentX = (xav - xn) * 100 / (xx - xn)
      .CurrentY = (yav - yn) * 100 / (yx - yn)
      For t = 1 To Np4
         xap = Xmin - t * (Xmin + L_WS) / Np4
         yap = (L_WS + xap) ^ 2 / (2 * Abs(X3) * aX)
         yap = T0c * yap
         If yap <= 0 Then Exit For
         xap = (xap - xn) * 100 / (xx - xn): yap = (yap - yn) * 100 / (yx - yn)
         O.Line -(xap, yap), QBColor(13) 'jaunefoncé
       Next
   End If


r_HelA = 0 'sert à laire la moyenne des r2
If N_fm = 1 And Fresc.Tag <= 1 Then 'T1 -------------------------------------------- reg T1

      Debug.Print "Controle", Format(T0(nSM)) + " kPa", "Bz1_min = "; z1_min, "Bz1_max = "; z1_max, "Bz2_min = "; z2_min,
"Bz2_max = "; z2_max

      Pds = z1_max 'normal
      If z2_min > z1_max Then Pds = z2_min

      T1T2.Seek "=", zfm, 1: If T1T2.NoMatch Then Stop
      T0_CC = T0c
      sN = 0: sX = 0: sXX = 0: sY = 0: sYY = 0: sXY = 0
      nSM = T1T2("cod_T1"): coul = QBColor(T1T2("coul_pt"))
      'pente instannée
      AAPH_ARTNEG_variation_T0_chi_i nSM

      Do While T1T2("cas") = zfm
         If T1T2("cod_T1") > nSM Then 'on chnde droite
           ' If T1T2met("methode") <> "variation_T0" Then Stop 'petit verif
            'on ecrit la régression
             AAPH_ARTNEG_trace_R2 "T1"
            'il faut tout recalculer
            qOK = a_q + b_q * Log(T0(nSM) / T0(0))
            KqnZ1 = calcul_Kqn(qOK)
            xmi = N_hs ^ (1 - qOK / 2)
            cH_n = (Exp(xmi) + Exp(-xmi)) / 2
            sH_n = (Exp(xmi) - Exp(-xmi)) / 2
            z1_min = X_Z1 / KqnZ1
            z1_max = z1_min * cH_n

            'pente instannée
            AAPH_ARTNEG_variation_T0_chi_i nSM
            r = qOK + Log(chi_p2_i / chi_p1_i) / Log(N_hs)
             KrnZ2 = calcul_Kqn(r)
            xmi = N_hs ^ (1 - r / 2)
            cH_n = (Exp(xmi) + Exp(-xmi)) / 2
            sH_n = (Exp(xmi) - Exp(-xmi)) / 2
            z2_min = -L_WS / KrnZ2
```

```
        z2_max = z2_min * cH_n
        Pds = z1_max 'normal
        If z2_min > z1_max Then Pds = z2_min

       'on repart de zéro
        T0_CC = IIf(Frtyp.Tag = 1, T0(nSM), T0(nSM) / T0(0))
        sN = 0: sX = 0: sXX = 0: sY = 0: sYY = 0: sXY = 0
        nSM = T1T2("cod_T1"): coul = QBColor(T1T2("coul_pt"))
      End If

      If IsNull(T1T2("X1")) = 0 And IsNull(T1T2("pT1c")) = 0 Then
        xav = T1T2("X1"): yav = T1T2("pT1c")
        drap = 1
        If T1T2("cas") = 5 And T1T2("pt") = 22 And T1T2("cod_T1") = 5 Then z1_min = 2
        'If T1T2("cas") = 7 And T1T2("pt") = 12 And T1T2("cod_T1") = 3 Then z1_min = 2
        'If T1T2("cas") = 8 And T1T2("pt") = 19 And T1T2("cod_T1") = 4 Then z1_min = 2
        If T1T2("cas") = 9 And T1T2("pt") = 12 And T1T2("cod_T1") = 3 Then z1_min = 2
        If T1T2("cas") = 28 And T1T2("pt") = 27 And T1T2("cod_T1") = 3 Then z1_min = 2
        If xav > Abs(z1_min) Or yav < 0 Then 'hors zone
          drap = 0
        ElseIf xav <= Abs(z1_min) And xav > z1_min Then 'zone 1 VRAIE
            yap = T0_CC * (1 + xav * chi_p1_i * KqnZ1)
        ElseIf xav > Pds Then 'z1_max Then 'zone mixte
            xmi = T0_CC * (1 + xav * chi_p1_i * KqnZ1)
            ymi = T0_CC * chi_p2_i * (L_WS + KrnZ2 * xav)
            poids = (xav - z1_min) / (z1_max - z1_min) ' t / Np4 ' xmi = z1_min + t * (z1_max - z1_min) / Np4
            yap = (1 - poids) * xmi + poids * ymi
        ElseIf Pds = z1_max And xav > z2_min Then 'zone 2 vraie
            'petite verif
            'If xav < X_T0 Then Stop ' a verifier
            yap = T0_CC * (Abs(X_T0) + xav) * chi_p2_i * KrnZ2
        Else
            yap = 0
            'drap = 0
        End If
        If drap = 1 Then 'OK
          'If T1T2("cod_T1") = 5 Then Debug.Print xav, yav, yap
          T1T2.Edit: T1T2("pT1t") = yap: T1T2.Update
          sN = sN + 1
          sX = sX + yav: sXX = sXX + yav ^ 2
          sY = sY + yap: sYY = sYY + yap ^ 2
          sXY = sXY + yav * yap
        End If
      End If
      T1T2.MoveNext: If T1T2.EOF Then Exit Do
    Loop
      AAPH_ARTNEG_trace_R2 "T1"

End If


End With
End Sub
```

#### Sub AAPH_ARTNEG_Recalcul() 'recalcul en fonction du typ chosi

```
Dim z_coD As Integer, z_X1 As Single
'**** Informations recueillies ds T1T2met
   'T1T2met.Seek "=", z: If T1T2met.NoMatch Then Stop
    N_droites = T1T2met("N_facteur")
    For i = 0 To N_droites - 1
          vd = "T0" + IIf(i = 0, "c", "m" + Format(i))
          If IsNull(T1T2met(vd)) Then Stop
          If T1T2met(vd) = 0 And z <> 703 Then Stop
          T0(i) = T1T2met(vd)
    Next

'**** On travaille T1T2
'index=PK =cas + pt
T1T2.Seek "=", z, -99: If T1T2.NoMatch Then Stop

If IsNull(T1T2("cod_T1")) Then
   T1_pT = 0
Else
   T1_pT = T1T2("cod_T1")
End If

If IsNull(T1T2("cod_T2")) Then
   T2_pT = 0
Else
   T2_pT = T1T2("cod_T2")
End If
T1T2.Seek "=", z, 1: If T1T2.NoMatch Then Stop
z_coD = T1T2("cod_T1"): If z_coD <> 1 And zfm <> 19 And zfm <> 11 Then Stop 'verif
sN = 0: sX = 0: sXX = 0: sY = 0: sYY = 0: sXY = 0

Do While T1T2("cas") = z

  T1T2.Edit
  If T1_pT > 0 Then
    If IsNull(T1T2("X1")) = 0 And IsNull(T1T2("T1c")) = 0 And T1T2("cod_T1") > 0 Then
      Select Case Frtyp.Tag
          Case 1: T1T2("pT1c") = T1T2("T1c")  ' T1 en kPa
          Case 2: T1T2("pT1c") = T1T2("T1c") / T0(0) ' pT1 calculé sur TOc
          Case 3: T1T2("pT1c") = T1T2("T1c") / T0(T1T2("cod_T1") - 1) ' pT1 calculé sur TOm
       End Select
       z_X1 = T1T2("X1")
      If IsNull(T1T2("cod_T1")) = 0 And T1T2("cod_T1") > z_coD Then 'on change de Droite donc on EnRegistre
         If sN < 2 Then Stop
         'r
           r_regL = (sXY - sX * sY / sN) / Sqr((sXX - sX * sX / sN) * (sYY - sY * sY / sN))
           If r_regL < 0.8 Or r_regL > 1.00001 Then Stop
         'pente
           p_regL = (sXY - sX * sY / sN) / (sXX - sX * sX / sN)
           If p_regL < 0 Then Stop
         'ordonné à l'origine
           Y0_regL = sY / sN - p_regL * sX / sN
         'pente visoc
          p_regL = p_regL / T0(z_coD - 1) ' /T0(0) car car calculé à partir de ymi = T1T2("T1c") (voir plus bas)
          Y0_regL = Y0_regL / T0(z_coD - 1)
          If (p_regL - chi_p1) <= 0 Then
             qOK = 2.5
          Else
             Kqn = p_regL / chi_p1  'T1i/T0i=1+(Chi_p1+Sigma_i)*DX
             calcul_qoK (Kqn)
         End If
```

```
            'on enregistre
             T1T2met.Edit
                 tx1 = "_" + IIf(z_coD = 1, "c", "m" + Format(z_coD - 1))
                 T1T2met("p" + tx1) = p_regL 'T1i/T0i=1+(Chi_p1+Sigma_i)*DX
                 T1T2met("Y0" + tx1) = Y0_regL
                 T1T2met("r2" + tx1) = r_regL ^ 2
                 T1T2met("q" + tx1) = qOK
                 T1T2met("nu" + tx1) = chi_p1 * N_hs ^ -qOK * 10000000
             T1T2met.Update
             'et on repart
             z_coD = T1T2("cod_T1") 'en kpa
             sN = 0: sX = 0: sXX = 0: sY = 0: sYY = 0: sXY = 0
         End If
         If z_X1 >= Xmin And z_X1 <= 0 Then
            ymi = T1T2("T1c")
            sN = sN + 1
            sX = sX + z_X1: sXX = sXX + z_X1 ^ 2
            sY = sY + ymi: sYY = sYY + ymi ^ 2
            sXY = sXY + z_X1 * ymi
         End If
      End If
   End If
   If T2_pT > 0 Then
      If IsNull(T1T2("T2")) = 0 And T1T2("cod_T2") > 0 Then
        Select Case Frtyp.Tag
            Case 1: T1T2("pT2") = T1T2("T2")  ' T2 en kPa
            Case 2: T1T2("pT2") = T1T2("T2") / T0(0) ' pT2 calculé sur TOc
            Case 3: T1T2("pT2") = T1T2("T2") / T0(T1T2("cod_T2") - 1) ' pT2 calculé sur TOm
        End Select
      End If
   End If
   T1T2.Update
   T1T2.MoveNext
Loop
'***********
  If sN > 1 Then
    'r
     r_regL = (sXY - sX * sY / sN) / Sqr((sXX - sX * sX / sN) * (sYY - sY * sY / sN))
    'pente
     p_regL = (sXY - sX * sY / sN) / (sXX - sX * sX / sN)
    'ordonné à l'origine
     Y0_regL = sY / sN - p_regL * sX / sN
     p_regL = p_regL / T0(z_coD - 1)
     Y0_regL = Y0_regL / T0(z_coD - 1)
      If (p_regL - chi_p1) <= 0 Then 'pas de viscosité
          qOK = 2.5
      Else
        Kqn = p_regL / chi_p1
        calcul_qoK (Kqn) '1 / 8.01) * Log(2800000 / Kqn) '1.9
      End If
          'on enregistre
          T1T2met.Edit
                 tx1 = "_" + IIf(z_coD = 1, "c", "m" + Format(z_coD - 1))
                 T1T2met("p" + tx1) = p_regL
                 T1T2met("Y0" + tx1) = Y0_regL
                 T1T2met("r2" + tx1) = r_regL ^ 2
                 T1T2met("q" + tx1) = qOK
                 T1T2met("nu" + tx1) = chi_p1 * N_hs ^ -qOK * 10000000
         T1T2met.Update
End If
End Sub
```

#### Sub AAPH_ARTNEG_variation_T0(iX As String)

```
Dim ZONE As Integer, leg As Integer, ztx As String, cH_i As Single, sH_i As Single, Bz1_min As Single, Bz1_max As Single,
Bz2_min As Single
Dim Bpoids As Single, Super_theoriq As Integer
If zfm = 11 Or zfm = 19 Or zfm = 22 Then Exit Sub
If Fresc.Tag = 2 And Frpres.Tag = 4 Then Exit Sub 'uniqt T2
If N_droites = 1 Then Exit Sub
If N_droites < 1 Then Stop
If zfm = 3 Or zfm = 4 Then
    Select Case zfm
    Case 3 'Ford 81 F6
       T0(1) = T0(0) * (2.607 - 0.7143 * 2.6)
       T0(2) = T0(0) * (2.607 - 0.7143 * 3.1)
    Case 4 'Ford 81 F11
       T0(1) = T0(0) * (2.607 - 0.7143 * 3.1)
    End Select
Else
 Super_theoriq = 0 'normal on part sur les T0 mesurées
End If

If Super_theoriq = 1000 Then
   Select Case zfm
    Case 3 'Ford 81 F6
       T0(1) = T0(0) * (2.607 - 0.7143 * 2.6)
       T0(2) = T0(0) * (2.607 - 0.7143 * 3.1)
    Case 4 'Ford 81 F11
       T0(1) = T0(0) * (2.607 - 0.7143 * 3.1)
    Case 5 'Linai 2003 pCa slow fibre
       For i = 1 To 4
         xmi = Choose(i, 5.3, 5.49, 5.63, 5.84)
         ' q = Choose(i + 1, 4.5, 5.3, 5.49, 5.63, 5.84)
         Debug.Print "i= "; i
         Debug.Print "Avant : T0i = "; T0(i)
         T0(i) = T0(0) * (1 - 1 / (1 + Exp(30.5 - 5.5 * xmi))) ' 'a = 30.5:  b = 5.5 'OK1
         'T0(i) = T0(0) * (1 - 1 / (1 + Exp(13.685 - 2.4452 * xmi)))
         Debug.Print "Aprés : T0i = "; T0(i)
      Next
    Case 6 'Linai 2003 pCa
    End Select
End If

' 1 : on cherche la relation q_i =a_q + b_q * log(pT_i)
 sN = 0: sX = 0: sXX = 0: sY = 0: sYY = 0: sXY = 0
 For i = 0 To N_droites - 1 'la 1ere droites C=0
     tx1 = IIf(i = 0, "c", "m" + Format(i))
     Select Case iX
      Case "T0": xmi = Log(T0(i) / T0(0)): ztx = " * Log(pT0_i)"
      Case "temp2": xmi = Exp(T0(i) / T0(0)): ztx = " * exp(pT0_i)"
      Case "temp", "Rigor": xmi = T0(i) / T0(0): ztx = " * pT0_i"
     Case Else: Stop 'a faire
     End Select
     ymi = T1T2met("q_" + tx1)
     sN = sN + 1
     sX = sX + xmi: sXX = sXX + xmi ^ 2
     sY = sY + ymi: sYY = sYY + ymi ^ 2
     sXY = sXY + xmi * ymi
 Next 'i

     'regression q = a_q + b_q * log(T0i/T0c)
      If sN = 2 Then
        r_regL = 1: p_regL = 1
```

```
      Else
       'r
         r_regL = (sXY - sX * sY / sN) / Sqr((sXX - sX * sX / sN) * (sYY - sY * sY / sN))
       'pente
         p_regL = (sXY - sX * sY / sN) / (sXX - sX * sX / sN)
      End If
      'ordonné à l'origine
        Y0_regL = sY / sN - p_regL * sX / sN

'2: on stocke
            T1T2met.Edit
              T1T2met("r2_q_REG") = r_regL ^ 2
              T1T2met("a_q_REG") = Y0_regL
              T1T2met("b_q_REG") = p_regL
              If Trim(TAM(3).Text) = "0.1" And Trim(TAM(2).Text) = "1.9" Then
                Stop 'à verifier ce truc 2019
                 T1T2met("a_q_main") = Y0_regL: a_q = Y0_regL: TAM(2).Text = Format(a_q, "#.###")
                 T1T2met("b_q_main") = p_regL: b_q = p_regL: TAM(3).Text = Format(b_q, "0.###")
              End If
            T1T2met.Update

'3 on trace
With O
.FontBold = -1
'relatopn q en fonction de log(pTi)1
    If N_hs > 0 Then
        O.FontSize = 12: O.CurrentX = 20: O.CurrentY = 88
       O.Print "THEOR .:   q_i = " + Format(Y0_regL, "#.###") + " + " + Format(1 / Log(N_hs), "0.###") + ztx
       O.FontSize = 8: O.CurrentX = 46: O.CurrentY = 88
       O.Print "b_q = 1/log(N_hs)"
    End If

     O.FontSize = 12: O.CurrentX = 20: O.CurrentY = 85
    O.Print "REGR.   :   q_i = " + Format(Y0_regL, "#.###") + " + " + Format(p_regL, "0.###") + ztx
      O.FontSize = 8: O.CurrentX = 46: O.CurrentY = 85
    O.Print "PAS SE FOCALISER car regression calculèe"
    O.CurrentX = 46: O.CurrentY = 84
    O.Print "à partir des 2/3 pts entre 0 et -DXend"
    O.FontSize = 12: O.CurrentX = 20: O.CurrentY = 82
    O.Print "STOCKé :   q_i = " + Format(a_q, "#.###") + " + " + Format(b_q, "0.###") + ztx

 For i = 0 To N_droites - 1 'la 1ere droites C=0 deja tracée
    .DrawWidth = IIf(i = 0, 1, 6)
    .DrawStyle = IIf(i = 0, 2, 0)

    ZONE = 1

    'point haut' A ce stade a_q=a_q_main   b_q=b_q_main
       Select Case iX
         Case "T0": qOK = a_q + b_q * Log(T0(i) / T0(0))
         Case "temp2": qOK = a_q + b_q * Exp(T0(i) / T0(0))
         Case "temp", "Rigor": qOK = a_q + b_q * T0(i) / T0(0)
         Case Else: Stop 'a faire
       End Select

       Kqn = calcul_Kqn(qOK)
       xmi = N_hs ^ (1 - qOK / 2)
       cH_i = (Exp(xmi) + Exp(-xmi)) / 2
       sH_i = (Exp(xmi) - Exp(-xmi)) / 2
       Bz1_min = X_Z1 / Kqn
       Bz1_max = Bz1_min * cH_i
```

```
'Pente instantanée***************************************************************************
```

**AAPH_ARTNEG_variation_T0_chi_i i**

```
  D = chi_p1_i * Kqn

  Select Case Frtyp.Tag
    Case 1: xav = 1 / D ' T1m= T0m*[1+D* X]
    Case 2: xav = (2 * T0(0) / T0(i) - 1) / D ' pT1m'= Tm/T0c= T0m/T0c *[1+D* X]
    Case 3: xav = 1 / D ' pT1m=Tm/T0m= [1+D* X] '=(2 - 1) / D
   End Select
   yav = 2 * IIf(Frtyp.Tag = 1, T0(i), 1)  '* T0c '1.5 '=PT1c

'point bas
 xap = -1 / D
 yap = 0

 xap = Bz1_min
 ymi = 1 + chi_p1_i * Kqn * xap
 Select Case Frtyp.Tag
   Case 1: yap = T0(i) * ymi ' T1m= T0m*[1+D* X]
   Case 2: yap = ymi * T0(i) / T0(0) ' pT1m'= Tm/T0c= T0m/T0c *[1+D* X]
   Case 3: yap = ymi ' pT1m=Tm/T0m= [1+D* X]
  End Select
```

**AAPH_ARTNEG_var_choix_couleur**

```
xav = (xav - xn) * 100 / (xx - xn)
yav = (yav - yn) * 100 / (yx - yn)
xap = (xap - xn) * 100 / (xx - xn)
yap = (yap - yn) * 100 / (yx - yn)
O.Line (xav, yav)-(xap, yap), coul

'*************légende ZONE 1
If Frpres.Tag = 0 Then
  'legende en haut à gauche
  .FontSize = 10
  For leg = 0 To 4 'IIf(Ix = "T0", 6, 4) 'IIf(Fresc.Tag = 0, 10, 4)
     xmi = 9
     Select Case leg
       Case 0: tx1 = "droite " + IIf(i = 0, "REF", Format(i)): O.CurrentX = xmi
       Case 1: xav = T0(i) / T0(0): tx1 = "pT0_i": O.CurrentX = xmi + 6
       Case 2: xav = qOK: tx1 = "q_i": O.CurrentX = xmi + 14
       Case 3: xav = Kqn: tx1 = "Kqn_i":    O.CurrentX = xmi + 20
       Case 4: xav = Kqn * chi_p1_i: tx1 = "pT1_z1=1+": O.CurrentX = xmi + 27.5
     End Select

     O.CurrentY = 78 - i * 3

     O.ForeColor = coul 'QBColor(Choose(i, 2, 10, 1, 9, 12, 12, 12, 9, 9, 9, 6, 6, 13, 0, 0))
     If leg = 0 Then
        O.Print tx1
     Else
        If leg = 4 Then
           tx1 = tx1 + Format(xav, "0.###") + "*DX"
        Else
           tx1 = tx1 + " = " + Format(xav, "0.###")
        End If
        O.Print tx1
    End If
  Next 'leg
End If
```

```
'***********************
 r = qOK + Log(chi_p2_i / chi_p1_i) / Log(N_hs)
 Krn = calcul_Kqn(r)
 Bz2_min = -L_WS / Krn ': If Bz2_min > X_Z1 Then Stop 'pas normal

 If Bz2_min < Bz1_max Then 'zone 2 Vraie

    '**legende zone 2
     If Frpres.Tag = 0 Then
      For leg = 0 To 3 'IIf(Fresc.Tag = 0, 10, 4)
         xmi = 47
         Select Case leg
          Case 0: xav = r: tx1 = "r_i": O.CurrentX = xmi
          Case 1: xav = Krn: tx1 = "Krn_i": O.CurrentX = xmi + 5.5
          Case 2: xav = chi_p2_i * L_WS: tx1 = "pT1_z2": O.CurrentX = xmi + 12.5
          Case 3: xav = Krn * chi_p2_i: tx1 = "+": O.CurrentX = xmi + 20
         End Select

         O.CurrentY = 78 - i * 3
           O.ForeColor = coul 'QBColor(Choose(i, 2, 10, 1, 9, 12, 12, 12, 9, 9, 9, 6, 6, 13, 0, 0))
         If leg = 3 Then
            tx1 = tx1 + Format(xav, "0.###") + "*DX"
         Else
            tx1 = tx1 + " = " + Format(xav, "0.###")
         End If
         O.Print tx1
      Next 'leg
    End If

   'vraie zone 2 avec  X_T0 = -L_WS / KrnZ2 ' < 0
       xav = Bz1_max
       yav = chi_p2_i * L_WS + xav * chi_p2_i * Krn

       Select Case Frtyp.Tag
        Case 1: yav = T0(i) * yav ' T1m= T0m*[1+D* X]
        Case 2: yav = yav * T0(i) / T0(0) ' pT1m'= Tm/T0c= T0m/T0c *[1+D* X]
        Case 3: 'OK ' pT1m=Tm/T0m= [1+D* X]
       End Select

       xap = Bz2_min: yap = 0
        xav = (xav - xn) * 100 / (xx - xn)
       yav = (yav - yn) * 100 / (yx - yn)
       xap = (xap - xn) * 100 / (xx - xn)
       yap = (yap - yn) * 100 / (yx - yn)
       O.Line (xav, yav)-(xap, yap), coul
 End If


 '*** lissage entre zone 1 et zone 2
 xav = Bz1_min 'Fin zone 1vraie
 yav = (1 + xav * chi_p1_i * Kqn)
 Select Case Frtyp.Tag
   Case 1: yav = T0(i) * yav ' T1m= T0m*[1+D* X]
   Case 2: yav = yav * T0(i) / T0(0) ' pT1m'= Tm/T0c= T0m/T0c *[1+D* X]
   Case 3: 'OK ' pT1m=Tm/T0m= [1+D* X]
 End Select
 O.CurrentX = (xav - xn) * 100 / (xx - xn)
 O.CurrentY = (yav - yn) * 100 / (yx - yn)

  For t = 1 To Np4
        xmi = Bz1_min + t * (Bz1_max - Bz1_min) / Np4
        'If xmi < -L_WS / Krn Then 'X_T=0
```

```
        If xmi < Bz2_min Then 'X_T=0
           Exit For
        End If
        yav = (1 + xmi * chi_p1_i * Kqn)
        'yap = (L_WS / Krn + xmi) * chi_p2 * Krn
        yap = (Abs(Bz2_min) + xmi) * chi_p2_i * Krn
        Bpoids = t / Np4
        ymi = (1 - Bpoids) * yav + Bpoids * yap

        Select Case Frtyp.Tag
         Case 1: yap = T0(i) * ymi ' T1m= T0m*[1+D* X]
         Case 2: yap = ymi * T0(i) / T0(0) ' pT1m'= Tm/T0c= T0m/T0c *[1+D* X]
         Case 3: yap = ymi ' pT1m=Tm/T0m= [1+D* X]
        End Select

        xap = (xmi - xn) * 100 / (xx - xn)
        yap = (yap - yn) * 100 / (yx - yn)
        O.Line -(xap, yap), coul ', QBColor(IIf(xmi < X_z1, 11, 9))
     Next
 Next

End With
'T0c = 1
End Sub
```

#### Sub calcul_qoK(Kx As Single)

```
Dim Res As Single, siGne As Integer, siGne2 As Integer, qA As Single, qB As Single, qC As Single, qD As Single, t_q As Integer
If Kx < 1 Then Stop
If Kx = 1 Then qOK = 3: Exit Sub 'pas de viscosité
Res = 0.0002 'résoluton/erreur min
qOK = (1 / 8.01) * Log(2700000 / (Kx - 1))
xmi = N_hs ^ (1 - qOK / 2)
Kqn = xmi * (Exp(2 * xmi) + 1) / (Exp(2 * xmi) - 1) + (1 / N_hs) ^ (qOK - 1)
drap = 0
'etape 1
If Abs(Kqn - Kx) > Res Then 'on cherche
  siGne = IIf(Kqn > Kx, 1, -1)
  For t_q = 1 To 100
     yap = qOK + siGne * t_q / 10
     Kqn = calcul_Kqn(yap)
     If Abs(Kqn - Kx) <= Res Then qOK = yap: drap = 10: Exit For 'on cherche
     siGne2 = IIf(Kqn > Kx, 1, -1)
     If siGne2 <> siGne Then drap = 1: qA = qOK + siGne * (t_q - 1) / 10: Exit For
  Next
  If drap = 0 Then Stop
  If drap = 1 Then
      For t_q = 0 To 100
         yap = qA + siGne * t_q / 100
           Kqn = calcul_Kqn(yap)
         If Abs(Kqn - Kx) <= Res Then qOK = yap: drap = 10: Exit For 'on cherche
         siGne2 = IIf(Kqn > Kx, 1, -1)
         If siGne2 <> siGne Then drap = 2: qB = qA + siGne * (t_q - 1) / 100: Exit For
      Next
      If drap < 2 Then Stop
      If drap = 2 Then
         For t_q = 0 To 100
            yap = qB + siGne * t_q / 1000
            Kqn = calcul_Kqn(yap)
            If Abs(Kqn - Kx) <= Res Then qOK = yap: drap = 10: Exit For  'on cherche
            siGne2 = IIf(Kqn > Kx, 1, -1)
            If siGne2 <> siGne Then drap = 3: qC = qB + siGne * (t_q - 1) / 1000:: Exit For
         Next
         If drap < 3 Then Stop

         If drap = 3 Then
           For t_q = 0 To 100
               yap = qC + siGne * t_q / 10000
               Kqn = calcul_Kqn(yap)
               If Abs(Kqn - Kx) <= Res Then qOK = yap: drap = 10: Exit For  'on cherche
               siGne2 = IIf(Kqn > Kx, 1, -1)
               If siGne2 <> siGne Then drap = 4: qD = qC + siGne * (t_q - 1) / 1000:: Exit For
            Next
         End If

     End If
  End If
End If

'derniere verif
xmi = N_hs ^ (1 - qOK / 2)
Kqn = xmi * (Exp(2 * xmi) + 1) / (Exp(2 * xmi) - 1) + (1 / N_hs) ^ (qOK - 1) '- 1
If Abs(Kqn - Kx) > Res Then Stop

End Sub
```

#### Function calcul_Kqn(zq As Single) As Single

```
 If N_hs < 500 Then Stop 'bizarre_ surement un oubli
 If zq < 0.7 Then
   Stop
     calcul_Kqn = 10
 Else
     ymi = N_hs ^ (1 - zq / 2)
     calcul_Kqn = ymi * (Exp(2 * ymi) + 1) / (Exp(2 * ymi) - 1) + N_hs ^ (1 - zq)
 End If
End Function
```

#### Sub AAPH_ARTNEG_variation_T0_chi_i(iX As Integer)

```
    chi_p1_i = chi_p1 'par défaut
    chi_p2_i = chi_p2 'par défaut

    If Fresc.Tag = 1 And (zfm = 16 Or zfm = 17 Or zfm = 18) Then '2019
        bo1_pT = 9: bo2_pT = 0.5 '0.6
        dX_Ti = L_WS + (aX - L_WS) / (1 + Exp(-bo1_pT * (T0(iX) / T0(0) - bo2_pT)))
        chi_p1_i = 1 / (L_WS - dX_Ti / 2)
        chi_p2_i = (1 + X_Z1 * chi_p1_i) / (L_WS + X_Z1)
    End If
End Sub
```

### Sub AAPH_ARTNEG_var_choix_couleur()

```
 'Jaune vif &H0000FFFF&
   'Jaune foncé : &H0000C0C0&
   'Orange clair : &H0080C0FF&
   'Orange : &H000080FF&
   'Rose : &H00FF80FF&
   'Rose 2: &H00C0C0FF&
   'Rose 3: &H008080FF&
   'Bordeaux : &H000000C0&
   'Vert clair : &H00C0FFC0&
   'Vert : &H0000FF00&
   'Vert foncé : &H0000C000&
   'Bleu clair : &H00FFFFC0&
   'Bleu clair 2 : &H00FFFF80&Gris: &H00C0C0C0&
   'Bleu cile : &H00FFFF00&
   'Bleu gris : &H00C0C000&
   'Bleu roi : &H00C00000&
   'Violet pale : &H00FFC0C0&
   'Mauve intense
   'Gris foncé : &H00808080&


   'coul = QBColor(Choose(i, 11, 3, 13, 4, 10, 6))
   Select Case zfm
    Case 3 'Foe 81 F3 L0s'
    'Ford 81 F6 et F11
       coul = QBColor(Choose(i + 1, 0, 12, 13)) 'Ford 81 fig 11
    Case 4: coul = QBColor(Choose(i + 1, 0, 11)) 'Ford 81 fig 11
  'Linari 32003 pCa Fig 9C et
    Case 5: coul = QBColor(Choose(i + 1, 0, 12, 13, 5, 3, 10, 6)) 'mauve clair + mauve foncé + bleu+ bleu canard
    Case 6: coul = QBColor(Choose(i + 1, 0, 13, 5, 3, 10, 6)) 'mauve clair + mauve foncé + bleu canard

    Case 8: coul = QBColor(Choose(i + 1, 0, 12, 13, 10))
    Case 9, 12: coul = QBColor(Choose(i + 1, 0, 12, 10))
    Case 15, 21, 24: coul = QBColor(Choose(i + 1, 0, 12, 10, 4)) 'R ESCUL piazzesi

  'FORD 85
    Case 16, 17, 18: coul = QBColor(Choose(i + 1, 0, 11, 13, 4, 12)) 'bleu ciel /mauve clair/marron/rouge

   'tasaturyan 99
    Case 26: coul = QBColor(Choose(i + 1, 0, 6, 10, 6))


    Case 703: coul = QBColor(Choose(i + 1, 0, 10, 12, 10, 4))
    Case Else: coul = QBColor(Choose(i + 1, 1, 12, 13, 11, 4, 13, 10))
   End Select
End Sub
```

### Sub AAPH_ART_legende()

```
Dim titre As String
'"ph12_NEG_pT", "ph12_STRAIN", "ph12_RIGOR_pT", "ph12_TEMP_pT"
Select Case ressort.Tag
 Case "ph12_NEG_pT": k = 0
 Case "ph12_STRAIN"
    Select Case TE
     Case "Divers": k = 500
     Case "Edman": k = 1100
     Case "Lapin": k = 1200
     Case Else: Stop
   End Select
 Case "ph12_RIGOR_pT": k = 0
 Case "ph12_TEMP_pT": k = 0
 Case Else: Stop
End Select

T1T2met.Index = "PK"
 z = 0
 titre = ""
 For i = 0 To 30
     If ChT1T2(i) = 1 Then
        t = i: z = z + 1 'c'est ok
        T1T2met.Seek "=", ChT1T2(i).Tag + k
        If T1T2.NoMatch Then Stop
        titre = titre + Trim(T1T2met("fm")) + " + "
     End If
 Next
  'titre
   O.FontSize = 18
      O.CurrentX = -3
      O.CurrentY = 110
      O.ForeColor = QBColor(0)
   O.Print titre
If z <> 1 Then Exit Sub
N_fm = 1

    O.FontBold = -1
    O.FontSize = 12
     If N_fm = 1 Then '1 seule fibre donc on donne cses caracteritiques
        For i = 1 To 11
            tx1 = ""
            Select Case i
              Case 1
                  If IsNull(T1T2met("methode")) = 0 Then
                    tx1 = T1T2met("methode")
                 ElseIf IsNull(T1T2met("facteur")) = 0 Then
                   tx1 = tx1 + " (" + T1T2met("facteur") + ")"
                 End If
              Case 2: If IsNull(T1T2met("comm")) = 0 Then tx1 = T1T2met("comm")
              Case 3: tx1 = "N℉ig : " + T1T2met("n o_Fig")
              Case 4: tx1 = T1T2met("typ_step")
              Case 5: tx1 = T1T2met("l_Espece") + " (" + T1T2met("l_Musc") + ")"
              Case 6: If IsNull(T1T2met("du_p1")) = 0 Then tx1 = "durée Phase 1 = " + T1T2met("du_p1") + " µs"
              Case 7: tx1 = "T°exp. = " + T1T2met( "Texp") + " ℃"
              Case 8:  If IsNull(T1T2met("L0_s")) = 0 Then tx1 = "L0_s = " + T1T2met("L0_s") + " nm"
              Case 9:  If IsNull(T1T2met("L_fm")) = 0 Then tx1 = "L_fm = " + T1T2met("L_fm") + " mm"
              Case 10: If IsNull(T1T2met("S_fm")) = 0 Then tx1 = "S_fm = " + T1T2met("S_fm") + " 10-3 µm²"
              Case 11: If IsNull(T1T2met("T0c")) = 0 Then tx1 = "TO = " + Format(T1T2met("T0c")) + " kPa"

            End Select
            If Len(tx1) > 0 Then
```

```
            O.ForeColor = QBColor(IIf(i < 3, 12, 1))
            If ressort.Tag = "ph12_STRAIN" Then
              O.CurrentX = 5
            Else
              O.CurrentX = IIf(Right(ressort.Tag, 2) = "pT", -3, 15)
           End If
           O.CurrentY = 108 - i * 3
            O.Print tx1
        End If
      Next 'i
  End If
End Sub
```

#### CP4.2 Calculations and curves of Y, e, $e/e_0$ as a function of pT

The curve on the computer screen is done by calling the "SUB AAPH_STRAIN_trace()" routine.

Once the data have been recovered (See values displayed in the Tables of Paper 4 and in the Excel sheets of Supplement DA4), the experimental points from published articles and the theoretical curves from model equations are plotted as a function of the relative tension (Fig CP4.2).

A regression is performed between experimental and theoretical parameter values.

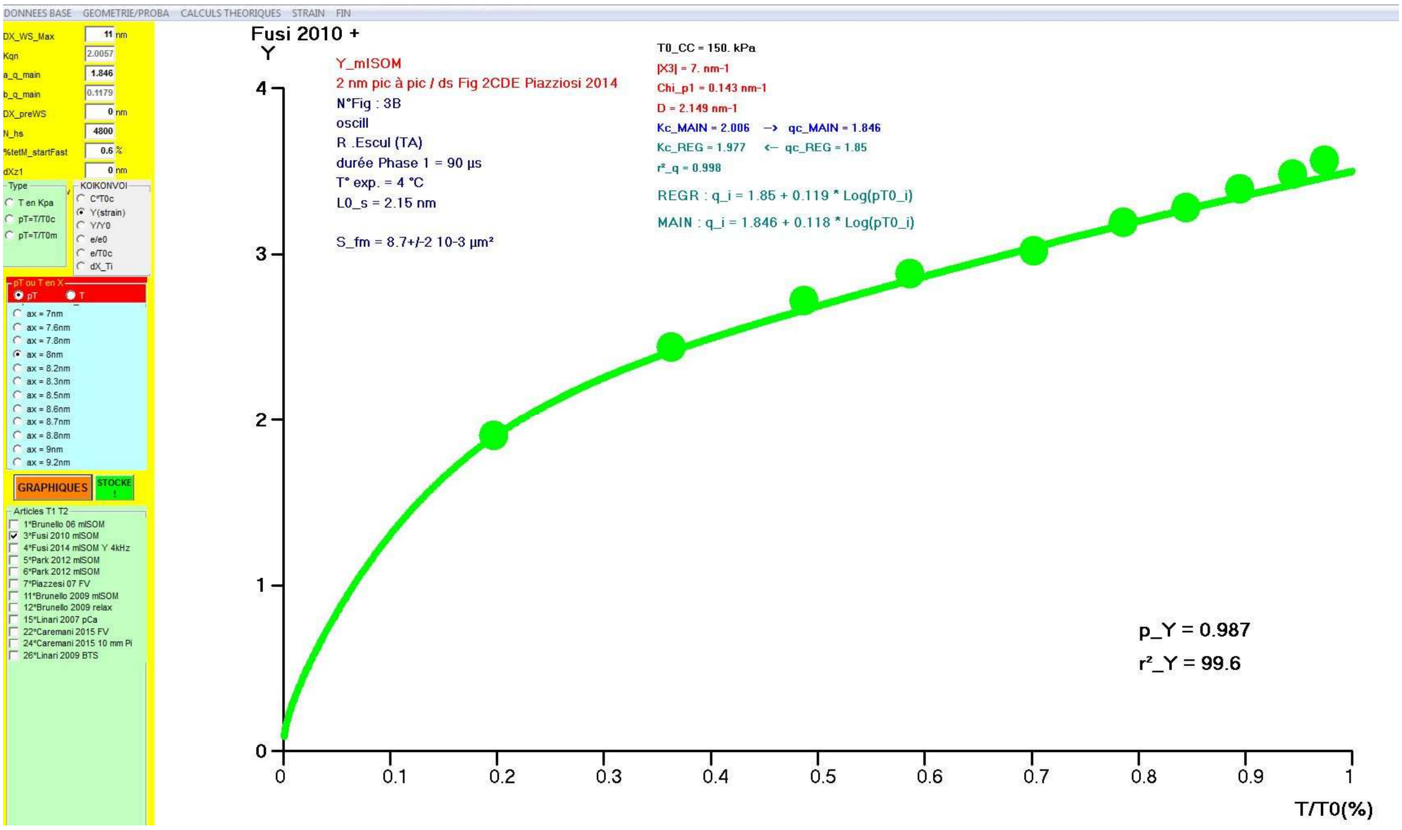


**Fig CP4.2 Screenshot after starting the the "Sub AAPH_STRAIN_trace()" routine.**

### Sub AAPH_STRAIN_trace()

```
Dim e_tt2 As Integer, arti As Integer, N_coché As Integer, a_qOK As Single, b_qOK As Single, Y0_dX_Ti As Single, p_dX_Ti As Single, C2_dX_Ti As Single ', COMPliance As Integer
Dim sXm As Single, sXXm As Single, sYt As Single, sYYt As Single, sXYt As Single, r_deux As Single, chi_i As Single, k_deb As Integer, rayon As Single, no_F As Integer ', pT0_borne As Single
If ressort.Tag <> "ph12_STRAIN" Then Stop
iLeg = 1 '  0=pas de comentaires 1=commentaires- pour graphiques de l'article
'COMPliance = 0 '1 compliance a la place de Y (strain)
If mmm = 1 Then O.AutoRedraw = True: O.Cls 'verifier que fonction autoredrax=tue
  O.Cls
Select Case TE
 Case "Divers": k_deb = 500
 Case "Edman": k_deb = 1100
 Case "Lapin": k_deb = 1200
End Select
Np4 = 1000
e_pt = 30 '50
e_tt = 6
e_tt2 = 2
'hpage =30 haut =140 large =110
impscale -12, Y0 + 130, X0 + larg, Y0
O.DrawMode = 13
'TAM(1).Enabled = 0 mis à la fin de AA_Neg_affi car Kqn = Kz1 calculé à partit de a_q main donc on le modifie pas
'STRAIN
L_WS = Val(TAM(0).Text) '10
aX = OaX(FraX.Tag).Tag * 2
Xmin = aX - L_WS '3.4 nm
X1 = aX / 2 '3.3 nm'3.33
X2 = -X1
X3 = X1 - L_WS '= (DXmin + DXmax) / 2 '-6.7 '-2*x1
chi_p1 = 1 / Abs(X3) 'pente ss Vis Zone 1   = 2/(DXmin + DXmax)
N_hs = Val(TAM(5).Text) '5000
If N_hs < 10000 Then
'If InStr(T1T2met("methode"), "mISOM") > 0 Then 'les champs b_q et Knz1=K0 sont bloqués
   TAM(1).Enabled = 0
   TAM(3).Enabled = 0
   b_q = 1 / Log(N_hs)
   TAM(3).Text = b_q '  coeff viscosité_tetM_liés independante T0
   a_q = Val(TAM(2).Text) 'a_q=q_T0c
   qZ1 = a_q
   KqnZ1 = calcul_Kqn(qZ1)
   TAM(1).Text = KqnZ1
Else 'les champs b_q et Knz1=K0 ne sont plus sont bloqués
    TAM(1).Enabled = -1
    TAM(3).Enabled = -1
    KqnZ1 = Val(TAM(1).Text) '0.1 '* 2 '0.25 - chi_p1 ' 0.033 * 3 'viscosité_MetZ_disjoints
    a_q = Val(TAM(2).Text) 'a_q=q_T0c
    b_q = Val(TAM(3).Text) '  coeff viscosité_tetM_liés independante T0
    N_hs = Val(TAM(5).Text) '5000
    calcul_qoK (KqnZ1) '1 / 8.01) * Log(2800000 / Kqn) '1.9
    qZ1 = qOK
End If
'Oesc(Ix).Tag = Choose(Ix, "Y", "Y/Y0", "e/e0", "e/T0c")
Select Case Fresc.Tag
     Case 0: vy = "C": yn = 0: yt = 5: yx = 30 'compliance
     Case 1 'Y
           vy = "Y": yn = 0: yt = 1
           Select Case zfm
             Case 503, 504, 511, 512, 1118, 1218: yx = 4
             Case 501, 507: yx = 5
             Case 515, 522, 524: yx = 8
             Case 1108, 1208: yx = 10
```

```
            Case Is > 1100: yx = 8
            Case Else: yx = 6
          End Select

    Case 2: vy = "Y/Y0": yn = 0: yx = 1: yt = 0.1
    Case 3: vy = "e/e0": yn = 0: yx = 1:   yt = 0.1
    Case 4: vy = "e/T0": yn = 0: yx = 0.3: yt = 0.05
    Case 5: vy = "dX_T0i": yn = 0: yx = IIf(zfm = 1109 Or zfm = 1110 Or zfm = 1111, 18, 15): yt = 1
    Case Else: Stop
End Select

'T0c = 1 '300 kPa?'on travaille sur pT pour l(instant
  T0c = 1:  vx = "T/T0(%)": xn = 0: xx = 1: xt = 0.1

impenvi
N_fm = 0 'compteur de tracées
tx2 = IIf(FrpTouT.Tag = 0, "pT", "T")

'pour savoir si 1 plusiers coché
For arti = 0 To N_articles - 1
     If ChT1T2(arti) = 1 Then N_coché = N_coché + 1
Next

zYb = "_dX_Ti"
For arti = 0 To N_articles - 1
  If ChT1T2(arti) = 1 Then 'c'est ok
        'rappel 1 : on prend exclivement d'apres choix de oesc(fr.esc.tag).tag
        'rapple 2 : N°fm (500_1100 (Edman) _1200(L apin) +?) concernée stockée ChT1T2(t).Tag = sTRain("cas")

       'en 1 pour temp etudie ensemble a partir du T0 de 0°C
       z = ChT1T2(arti).Tag
       'zfm = z + 500
       zfm = k_deb + z
       N_fm = N_fm + 1

       'les complments
       T1T2met.Seek "=", zfm: If T1T2met.NoMatch Then Stop
       sTRain.Seek "=", z, -99: If sTRain.NoMatch Then Stop
       If FrpTouT.Tag = 1 Then T0_CC = sTRain("T0")

       'Pvaluers consigne pour modélisation de dX_Ti
       no_F = 0 'par défaut
       If IsNull(sTRain("no_F")) = 0 Then
          no_F = sTRain("no_F")
          If IsNull(sTRain("Yhyp")) = 0 Then
             Yhyp = sTRain("Yhyp")
          Else
             Yhyp = 0
          End If
          bo1_pT = sTRain("bo1_pT")
          If IsNull(sTRain("bo2_pT")) = 0 Then
             bo2_pT = sTRain("bo2_pT")
          Else
             bo2_pT = 0
          End If
       End If

      'tracés des pts surtout tout laisser ici
      'O.DrawWidth = e_tt
       If Fresc.Tag = 5 Then 'dX_Ti

          'If zfm = 505 Or zfm = 506 Or zfm = 507 Or zfm = 511 Or zfm = 512 Or zfm = 516 Or zfm = 517 Then
```

```
      '   O.DrawWidth = e_pt 'courbe
      'ElseIf zfm = 1101 Or zfm = 1102 Then 'Parkn / Piazzesi FV2007
      '   O.DrawWidth = e_pt 'courbe
      'Else
      '  O.DrawWidth = e_pt * 1.5 'insert/encart
      'End If
    Else
       O.DrawWidth = e_pt
    End If
    '*******************************
    'If N_coché = 1 Then 'REGresion si un seul article coché
    sN = 0: sX = 0: sXX = 0: sY = 0: sYY = 0: sXY = 0
    sXm = 0: sXXm = 0: sYt = 0: sYYt = 0: sXYt = 0
    sTRain.MoveNext
    Do While sTRain("cas") = z
      'X_pT ou X_T
      If IsNull(sTRain("X_" + tx2)) = 0 And IsNull(sTRain("Y")) = 0 Then
       '1 recup Ymesuréees
          xav = sTRain("X_" + tx2) 'pT0i=T0i/T0c
          If Fresc.Tag = 5 Then
            yav = sTRain("dX_Ti")
          Else
            yav = sTRain("Y") 'Yi ou Yi/Y0 ou ei/e0 ou e1/T0
            If yav < 0 Then Stop
          End If

       If Fresc.Tag < 5 Then 'pas dX_Ti
         If FrCHI.Tag = 2 And no_F > 0 Then 'chi variable
            AAPH_STRAIN_CONSIGNE_calcul no_F
            If dX_Ti < 0 Then Stop
            chi_i = 1 / (L_WS - dX_Ti / 2)
         Else 'chi fixe
           chi_i = chi_p1
         End If
         Select Case Fresc.Tag
           Case 0: Kqn = 1 / (chi_i * yav * xav) ' Yi = 1/(chi_i* Ki*pT0i) --> Ki= 1/(chi_i* Yi*pT0i)
           Case 1: Kqn = 1 / (chi_i * yav) ' Yi = 1/(chi_i* Ki) --> Ki= 1/(chi_i* Yi)
           Case 2: Kqn = (chi_p1 * KqnZ1) / (chi_i * yav) ' Yi/Y0 = (chi_p1 * K0) / (chi_i* Ki) --> Ki= [[chi_p1 * Kz1)/chi_i]*(1/(Y0/Yi)
           Case 3: Kqn = yav * chi_p1 * KqnZ1 / (chi_i * xav) ' ei/e0 = [(chi_p1_i* Ki)/(chi_p1*Kz1)]*pT0i --> Ki= (e1/e0) * (Kz1*
chi_p1*Kz1)/(chi_i*pT0i)
           Case 4: Kqn = yav / (xav * chi_i) 'ei/T0c = pT0i * Ki * chi_i--> Ki= (e1/T0c)/(Chi_i*pT0i)
          End Select
          If zfm = 1216 And sTRain("pt") = 1 Then
          'Kqn=750
            Stop 'juste pour revoir
            qOK = 0.3
          Else
            If (Kqn < 1 Or Kqn > 16) Then Stop
             calcul_qoK (Kqn)
         End If

   '2/ on stocke
          sTRain.Edit
            sTRain("Ki_mes") = Kqn 'Ki calculé à partir de Yi ,yi/Y0, ei/e0 ou ei/T0
            sTRain("qi_mes") = qOK 'qi calculé à partir de Ki_mes par interpolatinc
          sTRain.Update

        '3/ on régresse
          If T1T2met("facteur") = "temperature" Then 'Davis temperature
            xmi = sTRain("X_pT")
          Else
            xmi = Log(sTRain("X_pT"))
```

```
    End If

   sN = sN + 1                         'qi= qc + p x Log(pT0i)   Temp: qi= qc + p x (pT0i-1)
   sX = sX + xmi: sXX = sXX + xmi ^ 2 'X=Log(pT0i)              Temp: X=(pT0i-1)
   sY = sY + qOK: sYY = sYY + qOK ^ 2 'Y=qi
   sXY = sXY + xmi * qOK
 End If 'if fresc.tag=5

'4/ on trace
   If Fresc.Tag < 5 Then
      If zfm = 501 Or zfm = 502 Or zfm = 1109 Or zfm = 1110 Or zfm = 1111 Then 'Brunello_2006
          O.DrawWidth = e_pt * 1.5
      ElseIf zfm = 503 Or zfm = 1115 Then
          O.DrawWidth = e_pt * 1.2
      ElseIf zfm = 507 Or zfm = 511 Or zfm = 512 Or zfm = 1117 Or zfm = 1118 Or zfm = 1112 Or zfm = 1120 Then
          O.DrawWidth = e_pt * 1.1
      Else
          O.DrawWidth = e_pt '* IIf(zfm = 1115, 1.2, 1)
      End If

   Else 'Fresc.Tag = 5 dX_Ti
     If zfm = 503 Or zfm = 505 Or zfm = 506 Or zfm = 516 Or zfm = 517 Then
         O.DrawWidth = e_pt
     ElseIf zfm = 511 Or zfm = 512 Then
          O.DrawWidth = e_pt * 1.1
     ElseIf zfm = 507 Or zfm = 1115 Or zfm = 1117 Or zfm = 1118 Then
       O.DrawWidth = e_pt * 1.2
     ElseIf zfm = 1205 Or zfm = 1206 Then
       O.DrawWidth = e_pt
     Else
       O.DrawWidth = e_pt * 1.5 'pour les inserts
     End If
   End If
    xap = (xav - xn) * 100 / (xx - xn)
    yap = (yav - yn) * 100 / (yx - yn)
   'croix
   If (zfm = 1102 And sTRain("pt") > 4) Or (zfm = 1119 And sTRain("pt") > 6) Then
      O.DrawWidth = IIf(Fresc.Tag < 5, 5, 8)
      xmi = 2 'IIf(Fresc.Tag < 5, 2, 3)
      ymi = 3 'IIf(Fresc.Tag < 5, 2.67, 4)
      O.Line (xap - xmi, yap)-(xap + xmi, yap), QBColor(sTRain("coul_pt"))
      O.Line (xap, yap - ymi)-(xap, yap + ymi), QBColor(sTRain("coul_pt"))

   Else 'point
    O.PSet (xap, yap), QBColor(sTRain("coul_pt"))
   End If

   If Fresc.Tag < 5 Then 'pas dX_Ti

    If k_deb >= 1100 Then
      If IsNull(sTRain("Ect_" + tx2)) = 0 And IsNull(sTRain("Ect_Y")) = 0 Then 'tx2 = IIf(FrpTouT.Tag = 0, "pT", "T")
       If sTRain("Ect_" + tx2) > 0 And sTRain("Ect_Y") > 0 Then
          O.DrawWidth = 1
          xmi = (sTRain("Ect_" + tx2) - xn) * 100 / (xx - xn)
          ymi = (sTRain("Ect_Y") - yn) * 100 / (yx - yn)
          rayon = IIf(xmi >= ymi, xmi, ymi) / 2
          O.Circle (xap, yap), rayon, QBColor(sTRain("coul_pt")), , , ymi / xmi
          'O.DrawWidth = e_pt
       End If
      End If
    End If
```

```
        '5/calcul Y theoriques
          If T1T2met("facteur") = "temperature" Then 'Davis temperature
            '   qOK = T1T2met("a_q_main") + xav / Log(N_hs)
           'ou qOK = T1T2met("a_q_main") + (xav-1) / Log(N_hs)
           'ou qOK = T1T2met("a_q_main")- 1/Log(N_hs) + xav / Log(N_hs)
            Stop 'a retester    avec excel
          Else
            qOK = T1T2met("a_q_main") + Log(xav) / Log(N_hs) 'qi_theorical
          End If
          Kqn = calcul_Kqn(qOK) 'Ki_theorical
          If FrCHI.Tag = 2 And no_F > 0 Then 'Chi_i variable et modélisée selon 4 types
            AAPH_STRAIN_CONSIGNE_calcul no_F
            chi_i = 1 / (L_WS - dX_Ti / 2)
          Else 'Chi_i fixe
            chi_i = chi_p1
          End If
         Select Case Fresc.Tag 'Yi_theorical
          Case 0: ymi = 1 / (chi_i * Kqn * xav) ' Yi = T0/(Chi_p1 * Yi*T0i)=1/(Chi_p1 * Yi*pT0i)
          Case 1: ymi = 1 / (chi_i * Kqn) ' Yi = 1/(Chi_p1 * Yi)
          Case 2: ymi = (chi_p1 * KqnZ1) / (chi_i * Kqn) ' Yi/Y0 = (Kz1*chi_p1)/(Ki/chi_p1_i)
          Case 3: ymi = xav * (chi_i * Kqn) / (chi_p1 * KqnZ1) 'ei/e0 =pT0i* Ki/chi_p1_i) / (Kz1*chi_p1)
          Case 4: ymi = Kqn * chi_i * xav ' ei/T0 = Ki* chi_p1*pT0i
         End Select
        '6/ on re-stocke
         sTRain.Edit
          sTRain("qi_th") = qOK 'qi_theorical eq_27_Papier4
          sTRain("Ki_th") = Kqn 'Ki_theorical eq_26_Papier4
          sTRain("Yt") = ymi    'Yi_theorical eq_28a_28b_Papier4
           Select Case Fresc.Tag '0=C 1=Y 2=Y/Y0 3=e/e0 4=e/T0
            Case 0:  xmi = yav * Kqn * xav ' Ci = 1/(chi_i* Ki*pT0i)  --> 1/(chi_i= Yi * Ki*pT0i
            Case 1:  xmi = yav * Kqn ' Yi = 1/(chi_i* Ki)  --> 1/chi_i= Yi * Ki
            Case 2: xmi = yav * Kqn / (KqnZ1 * chi_p1) ' Yi/Y0 = (chi_z1 * Kz1)/(chi_i* Ki)--> 1/chi_i= (Yi/Y0)*Ki/(Kz1*chi_p1)
            Case 3: xmi = (1 / yav) * Kqn * xav / (KqnZ1 * chi_p1) ' ei/e0 = pT0i * (chi_i* Ki)/(Chi_p1*Kz1)  --> 1/Chi_i) =
[1/(ei/e0)]*(Ki*pT0i)/(chi_p1*Kz1)
            Case 4: xmi = Kqn * xav / yav ' ei/T0 = T0i/T0 * chi_i* Ki  --> 1/Chi_i) = Ki*pT0i/(chi_p1*Kz1)
           End Select
          'Y sTRain("dX_Ti") = 2 * (L_WS - sTRain("Y") / (chi_p1 * ymi))
           sTRain("dX_Ti") = 2 * (L_WS - xmi) 'dX_Ti_mes eq_30a_30b_Papier4 avec xav=PT0i mesure et yav=Yi mesure
           'If sTRain("dX_Ti") > L_WS Then Stop 'verif
         sTRain.Update
        '7/ on re-régresse
          'sN fait avant
          sXm = sXm + yav: sXXm = sXXm + yav ^ 2 'Xm = Ymesuré
          sYt = sYt + ymi: sYYt = sYYt + ymi ^ 2 'Yy= Ytheoriq=qi
          sXYt = sXYt + yav * ymi
        End If
      End If
      sTRain.MoveNext: If sTRain.EOF Then Exit Do
    Loop
If Fresc.Tag < 5 Then 'on enregistre la regression ds T1T2met : q en fonction pT0i
    If sN < 2 Then Stop '?
    'r
    r_regL = (sXY - sX * sY / sN) / Sqr((sXX - sX * sX / sN) * (sYY - sY * sY / sN))
    'pente
    p_regL = (sXY - sX * sY / sN) / (sXX - sX * sX / sN)
    'ordonné à l'origine
    Y0_
    T1T2met.Edit
        T1T2met("r2_q_REG") = r_regL ^ 2
        T1T2met("a_q_REG") = Y0_regL
        T1T2met("b_q_REG") = p_regL
    T1T2met.Update
```

```
  '***************************************** regeression entre Ymesurées et Y theoriques du modele
   O.ForeColor = coul
     'pente
     'droite passant par l'origine maois piege car pb d'homthétie
       p_regL = sXYt / sXXm
       r_deux = 1 - (sYYt - 2 * p_regL * sXYt + sXXm * (p_regL ^ 2)) / (sYYt - sYt * sYt / sN)
    'donc on revient à la normale
      'p_regL = (sXYt - sXm * sYt / sN) / (sXXm - sXm * sXm / sN)
      'r_regL = (sXYt - sXm * sYt / sN) / Sqr((sXXm - sXm * sXm / sN) * (sYYt - sYt * sYt / sN))
     If r_deux > 1 Then Stop 'bizarre
     txV = Choose(Fresc.Tag + 1, "C*T0c", "Y", "Y/Y0", "e/e0", "e/T0c")
     O.ForeColor = QBColor(0)
     O.CurrentX = 80: O.CurrentY = 20: O.Print "p_" + txV + " = " + Format(p_regL, "0.###")
     O.CurrentX = 80: O.CurrentY = 15: O.Print "r²_" + txV + " = " + Format(r_deux * 100, "###.##")
End If 'fresc.tag<5
 '***************************************** regeression entre Ymesurées et Y theoriques du modele
    'trace courbe adhoc
     O.DrawStyle = 0
     O.DrawWidth = e_tt
    'pt départ
     xav = 1 / Np4 '* IIf(FrpTouT.Tag = 0, 1, T0_CC)
     If FrCHI.Tag = 2 And no_F > 0 Then 'Chi variable
          AAPH_STRAIN_CONSIGNE_calcul no_F
        chi_i = 1 / (L_WS - dX_Ti / 2)
     Else
        chi_i = chi_p1
     End If
     qOK = T1T2met("a_q_main") + T1T2met("b_q_main") * Log(1 / Np4) 'qoK theorique
      Kqn = calcul_Kqn(qOK) 'Ki théorique
     Select Case Fresc.Tag
      Case 0: yav = 1 / (chi_i * Kqn * xav)
      Case 1: yav = 1 / (chi_i * Kqn) ' Yi = 1/(Chi_p1 * Yi)
      Case 2: yav = (chi_p1 * KqnZ1) / (chi_i * Kqn) ' Yi/Y0 = (Kz1*chi_p1)/(Ki/chi_p1_i)
      Case 3: yav = xav * (chi_i * Kqn) / (chi_p1 * KqnZ1) 'ei/e0 =pT0i* Ki/chi_p1_i) / (Kz1*chi_p1)
      Case 4: yav = Kqn * chi_i * xav ' ei/T0 = Ki* chi_p1*pT0i
      Case 5: xav = 0: yav = 0
     End Select
    O.CurrentX = (xav - xn) * 100 / (xx - xn)
    O.CurrentY = (yav - yn) * 100 / (yx - yn)
    Select Case zfm
     Case 505, 506 'Park dogfish
       drap = IIf(zfm = 505, 7, 12) 'stepL=gris osc=rouge
       If Fresc.Tag = 5 Then drap = 11: O.DrawWidth = 12
    'Case 507 'piazzesi FV dX_Ti
    '  If Fresc.Tag = 5 Then drap = 0: O.DrawWidth = 4
    Case 516, 1212 'EDman 97 FV
      drap = 4:   If Fresc.Tag = 5 Then O.DrawWidth = 4
    Case 517, 1113, 1213 'EDman 97 mISOM
     drap = 12:   If Fresc.Tag = 5 Then O.DrawWidth = 4
    Case Else
       If FrCHI.Tag = 2 And no_F > 0 Then
         If zfm = 503 Or zfm = 508 Or zfm = 509 Or zfm = 510 Or zfm = 1117 Or zfm = 1118 Then
            O.DrawWidth = 9 'IIf(zfm = 1115, 9, 12) 'cecchi 86 specail
         ElseIf zfm = 511 Or zfm = 512 Then
            O.DrawWidth = 7
         ElseIf Fresc.Tag < 5 And (zfm = 507 Or zfm = 1112 Or zfm = 1120) Then
             O.DrawWidth = 7
         Else
            O.DrawWidth = 12
         End If
          drap = Choose(Fresc.Tag + 1, 1, 10, 2, 12, 13, 11, 0) 'C=11 bleu ciel Y=vert clair Y/Y0=vert foncé e/e0=rouge e/T0 mauve
       Else
```

```
          O.DrawWidth = 4
          drap = Choose(Fresc.Tag + 1, 1, 2, 2, 5, 5, 8, 0) 'C=11 bleu ciel Y=vert clair Y/Y0=vert foncé e/e0=rouge e/T0 mauve
        End If
      End Select
      If zfm = 511 Then drap = 10
      If zfm = 512 Then drap = 3 'petit rectificatif
      coul = QBColor(drap) 'bleu
      If FrpTouT.Tag = 1 Then Stop 'je fais pas
      For i = IIf(Fresc.Tag = 5, 1, 2) To Np4 ' tx2 = IIf(FrpTouT.Tag = 0, "pT", "T")
       'T0i ou pT0i
        xav = i / Np4 ' * IIf(FrpTouT.Tag = 0, 1, T0_CC)
       'chi_p1_i
        If FrCHI.Tag = 2 And no_F > 0 Then 'Chi variable
           AAPH_STRAIN_CONSIGNE_calcul no_F
          chi_i = 1 / (L_WS - dX_Ti / 2)
        Else
           chi_i = chi_p1
        End If
         'Ki
         a_qOK = T1T2met("a_q_main")
         b_qOK = T1T2met("b_q_main")
         If T1T2met("facteur") = "temperature" Then
            qOK = a_qOK + b_qOK * i / Np4
         Else
            qOK = a_qOK + b_qOK * Log(i / Np4)
         End If
        'qOK = T1T2met("a_q_REG") + T1T2met("b_q_REG") * Log(i / Np4) 'toujours log (pT)
         Kqn = calcul_Kqn(qOK)
         Select Case Fresc.Tag
          Case 0: yav = 1 / (chi_i * Kqn * xav)  ' Compliance Yi = 1/(Chi_p1 * Yi)
          Case 1: yav = 1 / (chi_i * Kqn) ' Yi = 1/(Chi_p1 * Yi)
          Case 2: yav = (chi_p1 * KqnZ1) / (chi_i * Kqn) ' Yi/Y0 = (Kz1*chi_p1)/(Ki/chi_p1_i)
          Case 3: yav = xav * (chi_i * Kqn) / (chi_p1 * KqnZ1) 'ei/e0 =pT0i* Ki/chi_p1_i) / (Kz1*chi_p1)
          Case 4: yav = Kqn * chi_i * xav ' ei/T0 = Ki* chi_p1*pT0i
          Case 5
            If FrCHI.Tag = 2 And no_F > 0 Then 'Chi variable
              yav = dX_Ti
            Else
              yav = aX
            End If
         End Select
         xap = (xav - xn) * 100 / (xx - xn)
         yap = (yav - yn) * 100 / (yx - yn)
         O.Line -(xap, yap), coul
      Next 'i
    End If 'If ChT1T2(arti) = 1 Then 'c'est ok
Next 'arti

'__il faut d'abord trace les courbes selon choix puis les points (à cause differentes unites en Y
If Fresc.Tag = 5 Then
    ' trait dX_Max dans insert /encart
      O.DrawWidth = 1: O.DrawStyle = 2
      ymi = (L_WS - yn) * 100 / (yx - yn)
      O.Line (0, ymi)-(100, ymi), QBColor(0)
      ' O.DrawWidth = e_tt * 2
   ' trait dX_T
    ymi = (aX - yn) * 100 / (yx - yn)
    O.Line (0, ymi)-(10, ymi), QBColor(0) 'coul
```

```
    'equation
    If FrCHI.Tag = 2 And no_F > 0 Then AAPH_STRAIN_CONSIGNE_affi no_F
ElseIf iLeg = 1 Then
    O.FontSize = 11
    O.FontBold = -1
   'legende 'legende en haut à gauche
    For i = 1 To 9
        Select Case i
          Case 1: xav = T0_CC: tx1 = "T0_CC"
          Case 2: xav = Abs(X3): tx1 = "|X3|"

          Case 3: xav = chi_p1: tx1 = "Chi_p1"
          Case 4: xav = chi_p1 + KqnZ1: tx1 = "D"

          Case 5: xav = KqnZ1: tx1 = "Kc_MAIN"
          Case 6: xav = qZ1: tx1 = "-->   qc_MAIN"

          Case 7: xav = T1T2met("a_q_REG"): tx1 = "<--  qc_REG"
          Case 8: qOK = T1T2met("a_q_REG"): xav = calcul_Kqn(qOK): tx1 = "Kc_REG"

          Case 9: xav = T1T2met("r2_q_REG"): tx1 = "r²_q"
        End Select

        O.CurrentX = IIf(i = 6 Or i = 7, 60, 50) - 15
        O.CurrentY = Choose(i, 107, 104, 101, 98, 95, 95, 92, 92, 89, 86, 83, 80)

        O.ForeColor = QBColor(Choose(i, 0, 12, 12, 12, 9, 9, 3, 3, 3, 5, 8, 7))
        tx1 = tx1 + " = " + Format(xav, "0.###")
        Select Case i
          Case 1: tx1 = tx1 + " kPa"
          Case Is < 5: tx1 = tx1 + " nm-1"
        End Select
        O.Print tx1
    Next 'i

    tx1 = IIf(T1T2met("facteur") = "temperature", " * pT0_i", " * Log(pT0_i)")
    O.FontSize = 12
    O.CurrentX = 45 - 10: O.CurrentY = 85
    O.Print "REGR : q_i = " + Format(T1T2met("a_q_REG"), "#.###") + " + " + Format(T1T2met("b_q_REG"), "0.###") + tx1
    O.CurrentX = 45 - 10: O.CurrentY = 81
    O.Print "MAIN : q_i = " + Format(T1T2met("a_q_MAIN"), "#.###") + " + " + Format(T1T2met("b_q_MAIN"), "0.###") + tx1
    AAPH_ART_legende

End If

End Sub
```

#### Sub AAPH_STRAIN_CONSIGNE_calcul(Iz As Integer)

```
 'Iz = No_F de la fonction modelisant consigne vers dX_Ton calul Y

 'xav = pT0i

 Select Case Iz

  Case 1 'montée exponentille : pCA,Pi
      dX_Ti = aX * (1 - Exp(-bo1_pT * xav) * (1 + bo2_pT * xav))

  Case 2 ' descente exponentielle de dX_Max à dX_T : montée en isométrie
      dX_Ti = aX + (L_WS - aX) * Exp(-bo1_pT * xav) * (1 + bo2_pT * xav)

  Case 3 ' descente exponentielle de dX_Max àdX_T avec initaition psauo periodique : montée en isometrie
       dX_Ti = aX + (L_WS - aX) * Exp(-bo1_pT * xav) * Cos(bo2_pT * xav)

  Case 4 'sigmoide descendante de dX_T à 0 (relaxation apres tetanisation )
       dX_Ti = aX / (1 + Exp(-bo1_pT * (xav - bo2_pT)))

  Case 5 'SIGMOIDE montante : courbe FV
     dX_Ti = L_WS + (aX - L_WS) / (1 + Exp(-bo1_pT * (xav - bo2_pT)))


  Case Else: Stop

 End Select

  If dX_Ti > 17 Then Stop

End Sub
```

#### Sub AAPH_STRAIN_CONSIGNE_affi(z1 As Integer)

```
Dim e_ht As Integer
e_ht = 4
 O.FontSize = 12
 xap = 30: yap = 12
 k = Choose(z1, 1, 2, 2, 3, 4)
 If k = 2 Or k = 4 Then O.CurrentX = xap + 15: O.CurrentY = yap + 2 * e_ht: O.Print "dX_Max - dX_T = " + Format(L_WS - aX) + " nm"

 O.CurrentX = xap: O.CurrentY = yap: O.Print "Langda_" + Format(k) + " = " + Format(bo1_pT)
 If bo2_pT <> 0 Then O.CurrentX = xap: O.CurrentY = yap - e_ht: O.Print "Omega_" + Format(k) + " = " + Format(bo2_pT)

 Select Case z1
  Case 1 'exp montante : : pCA,Pi
      If bo2_pT = 0 Then
         tx1 = "dX_Ti = dX_T * (1- Exp(-Langda_1 * pT0i )"
      Else
         tx1 = "dX_Ti = dX_T + [1- Exp(-Langda_1 * pT0i) * (1 + Omega_1 * pT0i)]"
      End If
      tx2 = "EXPO montante : CAS Géneral"

  Case 2 'descente exponentielle de dX_Max à dX_T : montée en isométrie
     tx1 = "dX_Ti = dX_T + (dX_Max - dX_T) * Exp(-Langda_2 * pT0i) * (1 + Omega_2 * pT0i)"
     tx2 = "EXPO descendante : CAS Géneral"

  Case 3 'descente exponentielle de dX_Max àdX_T avec initaition psauo periodique : montée en isometrie
     tx1 = "dX_Ti = dX_T + (dX_Max - dX_T) * Exp(-Langda_2 * pT0i) * Cos(Omega_2 * pT0i )"
     tx2 = "EXPO descendante + COS (oscillations)"

  Case 4 ' descendante de dX_T à 0 (relaxation apres tetanisation )
      tx1 = "dX_Ti = dX_T / [ 1 + exp(-Langda_3 * (pT0i - Omega_3)) ]"
      tx2 = "SIGMOIDE descendante"

  Case 5 'SIGMOIDE montante de dX_T à dX_Max : courbe FV
      tx1 = "dX_Ti = dX_Max+(dX_T-dx_Max) / [1+exp(-Langda_4 * (pT0i -Omega_4))] "
      tx2 = "sigmoide montante"

  Case Else: Stop
End Select

O.CurrentX = xap: O.CurrentY = yap - 2 * e_ht
O.Print "Eq :  " + tx1
O.CurrentX = xap - 7: O.CurrentY = yap + e_ht: O.Print "No fonction : " + Format(z1) + "   (" + tx2 + ")"


End Sub
```
