## Supplementary Chapter. Calculation of T1 with its two elastic and viscous components for "Mechanical model of muscle contraction. 4. Theoretical calculations of the tension during the isometric tetanus plateau and the tension exerted at the end of phase 1 of a length step"

### S4.J Supplementary Chapter of Paper 4

### Calculation of the tension at the end of phase 1 of a length step (T1) with its two elastic and viscous components

#### J.1 Viscosity force

The Stokes-Einstein formula provides an expression of the viscosity force ($F_{Visc}$) applying to a sphere moving in a fluid at velocity V. The force $F_{Visc}$, colinear to V and opposing the displacement, is written classically:

$$F_{Visc} = -(6 \cdot \pi \cdot \eta \cdot r) \cdot V$$

where $\eta$ is the viscosity coefficient of the liquid in which the spherical molecule moves (for example: $\eta_{water}(20°C)=10^{-3}$ Pa); r is the radius of the sphere.

In general, a body moving in a viscous fluid at the absolute velocity V is subjected to a force $F_{Visc}$, opposite to V and varying, as a first approximation, linearly with V such that :

$$F_{Visc} = -\phi \cdot V \qquad \text{(J1)}$$

where $\phi$ is the proportionality coefficient between $F_{Visc}$ and V, which depends on the experimental conditions (geometry of the material body, composition of the viscous liquid, temperature, etc).

#### J.2 Viscosity forces during shortening ($\Delta L<0$) or lengthening ($\Delta L>0$) of a muscle fiber

An isolated muscle fiber is composed of $N_m$ identical myofibril. Each myofibril of the fiber consists of $N_S$ sarcomeres arranged in series; see Fig F1a in Supplement S2.F to Paper 2. Each sarcomere is numbered from 1 to $N_S$ with the index s. A sarcomere is divided into two half-sarcomeres (Fig F1b), one located on the left (hsL) and the other on the right (hsR). Each half-sarcomere (hs) is numbered from 1 to $2N_S$, the even numbers corresponding to hsR and the odd numbers to hsL.

After being isometrically tetanized, the fiber is shortened at a constant speed V by a length step ($\Delta L$) during $\tau_{p1}$, the time of phase 1. We consider the material sets {Z-disk$_s$+2Afil} and {M-disk$_s$+2Mfil} with respect to sarcomere n° s; the solid {Z-disk$_s$+2Afil} is composed of the Z-disk n° s associated with actin filaments (2Afil) located to the right and left; the solid {M-disk$_s$+2Mfil} is composed of the M-disk n° s associated with myosin filaments (2Mfil) located to the right and left; see paragraph F.1 of Supplement S2.F. A change in fiber length at constant velocity implies that the absolute or relative speeds of {Z-disk$_s$+2Afil} and {M-disk$_s$+2Mfil} are also constant; see paragraph F.2 of Supplement S2.F.

The absolute velocities of {Z-disk$_s$+2Afil} and {M-disk$_s$+2Mfil} are calculated in (F6) and (F8) in supplement S2.F, equations that are reformulated below:

$$V_{Zs} = \frac{\Delta L_{Zs}}{\tau_{p1}} = \frac{\sum_{h=1}^{2s} \Delta X_h}{\tau_{p1}} \quad \text{(J2a)}$$

$$V_{Ms} = \frac{\Delta L_{Ms}}{\tau_{p1}} = \frac{\sum_{h=1}^{2s-1} \Delta X_h}{\tau_{p1}} \quad \text{(J2b)}$$

where h is the index of a hs; $\Delta X_h$ is the algebraic value of the length change of hs n° h at the end of phase 1.

During $\tau_{p1}$, the viscosity forces that slow down the displacement of the assemblies {Z-disk$_s$+2Afil} and {M-disk$_s$+2Mfil} are calculated according to (J1):

$$T_{Visc,Zs} = - \phi_{Zs} \cdot V_{Zs} \quad \text{(J3a)}$$

$$T_{Visc,Ms} = - \phi_{Ms} \cdot V_{Ms} \quad \text{(J3b)}$$

where $\phi_{Zs}$ and $\phi_{Ms}$ are the proportionality coefficients characterizing, respectively, the 2 aforementioned sets in the presence of viscosity.

We make an additional hypothesis: the proportionality constants between viscosity force and displacement velocity of {Z-disk$_s$+2Afil} and {M-disk$_s$+2Mfil} are equal. Otherwise formulated:

$$\phi_{hs} = \phi_{Zs} = \phi_{Ms} \quad \text{(J4)}$$

where $\phi_{hs}$ is the characteristic proportionality coefficient common to both solids.

The parameter ν is introduced:

$$\nu = \frac{\phi_{hs}}{T0_m \cdot \tau_{p1}} \quad \text{(J5)}$$

where $T0_m$ is the constant value exerted at the 2 extremities of the myofibril (Fig J1a) and on the 2 edges of each hs during the isometric tetanus plateau, value calculated in (I18) in Supplement S4.I; ν has as unit the inverse of a length and will be expressed in $nm^{-1}$.

Equations (J3a) and (J3b) are recombined according to (J2a), (J2b), (J4) and (J5) in:

$$T_{Visc,Zs} = - T0_m \cdot \nu \cdot \Delta L_{Zs} \quad \text{(J6a)}$$

$$T_{Visc,Ms} = - T0_m \cdot \nu \cdot \Delta L_{Ms} \quad \text{(J6b)}$$

#### J.3 Calculation of the tension at the end of phase 1 in the presence of viscosity when all hs shortenings belong to Zone 1

***Recalls of results given in paragraph I.6 of Supplement S4.I***

*Zone 1 is defined by the interval [-$\delta X_{z1}$ ; 0] where $\delta X_{z1}$ is a value between $\delta X_E$ and $2\cdot\delta X_E$ (Fig I3), and where $\delta X_E$ is a linear range defined in (I23). In the absence of viscosity, the tension ($T1_{Elas,z1}$) exerted on both sides of a hs comes exclusively from the forces deriving from the elastic energy of the WS myosin heads and is formulated in Zone 1 with (I52a):*

$$T1_{Elas,z1} = T0_m \cdot (1 + \chi_{z1} \cdot \Delta X) \tag{J7}$$

*where $\chi_{z1}$ is the coefficient of elastic origin stiffness which is written according to (I52b):*

$$\chi_{z1} = \frac{1}{|X_{down}|} = \frac{1}{\delta X_{Max} - \delta X_T / 2} \tag{J8}$$

*and where $\Delta X$ is the same shortening of the $N_{hs}$ hs of a myofibril in the absence of viscosity, equal according to (I39) to:*

$$\Delta X = \frac{\Delta L}{N_{hs}} \tag{J9}$$

The behavioural pattern of a myofibril consisting of n hs in series is now studied in the presence of viscosity. We follow a recurrence reasoning by specifying that the shortenings of the n hs all belong to Zone 1.

We are interested in the end of phase 1, the precise moment when the shortening of the fiber at constant speed V reaches the ΔL value of the length step and the fiber is in isometry because the shortening stops (Fig 1 of Paper 4). In the presence of viscosity, the forces applying to the assemblies {Z-disk$_s$+2Afil} and {M-disk$_s$+2Mfil} are of two types:

1/ an elastic input formulated in (J7), valid equation since the fiber is in isometry

2/ a viscous input created by the shortening speed formulated in (J6a) and (J6b)

The hs n° 1 associated with the Z-disk n° 0 which is fixed is the most left-handed hs (Fig J1). The hs n° 1 is considered as the most proximal hs with respect to the point of application of the tension measured using a force sensor whose signal is delivered by a transducer.

The hs n° n associated with the Z-disk n° n is located at the moving end of the fiber where the tension imposed by the experimenter is applied. The hs n° n is the most right-handed hs (Fig J1) considered as the most distal hs with respect to the point of application of the measured tension.

To simplify, the coefficient $\chi_{z1}$ will be noted $\chi$, the tension $T0_m$ exerted at the 2 ends of the myofibril during the isometric tetanus plateau will be noted T0' (Fig J1a) and the tension $T1_m$ exerted at the 2 ends of the myofibril at the end of phase 1 will be noted T1' (Fig J1b).

***J.3.1 n = 1 with $\Delta L = \Delta X_1$***

At the end of phase 1, hs n° 1 is shortened by $\Delta X_1$ in Zone 1 and T1', the resultant of the forces applied to the {$M_1$+Mfil} solid is the sum, on the one hand, of the tension exerted by the WS myosin heads in hs n° 1, and on the other hand, of the viscosity force. The linear momentum principle (*LMP*) is applied to {$M_1$+Mfil}. Equations (J6b) and (J7) lead to:

$$0 = T1' - T0' \cdot (1 + \chi \cdot \Delta X_1) - T0' \cdot \nu \cdot \Delta L$$

$$\Rightarrow \quad \frac{T1'}{T0'} = [1 + (\chi + \nu) \cdot \Delta L]$$

The myofibril composed of 1 hs behaves like a linear spring.

***J.3 2 n = 2 with $\Delta L = \Delta X_1 + \Delta X_2$***

At the end of phase 1, hs n° 1 and 2 shorten by $\Delta X_1$ and $\Delta X_2$ in Zone 1

*LMP* to {$M_1$+2Mfil}: $0 = -T0' \cdot (1 + \chi \cdot \Delta X_1) + T0' \cdot (1 + \chi \cdot \Delta X_2) - T0' \cdot \nu \cdot \Delta X_1$

$$\Rightarrow \quad \Delta X_2 = \left(1 + \frac{\nu}{\chi}\right) \cdot \Delta X_1$$

$$\Rightarrow \quad \Delta L = \Delta X_1 + \Delta X_2 = \left(2 + \frac{\nu}{\chi}\right) \cdot \Delta X_1$$

$$\Rightarrow \quad \Delta X_2 = \left(\frac{1 + \frac{\nu}{\chi}}{2 + \frac{\nu}{\chi}}\right) \cdot \Delta L$$

*LMP* to {$Z_1$+Afil}: $0 = T1' - T0' \cdot (1 + \chi \cdot \Delta X_2) - T0' \cdot \nu \cdot \Delta L$

$$\Rightarrow \quad \frac{T1'}{T0'} = [1 + \chi \cdot \Delta X_2 + \nu \cdot \Delta L] = \left[1 + \left(\chi \cdot \frac{1 + \frac{\nu}{\chi}}{2 + \frac{\nu}{\chi}} + \nu\right) \cdot \Delta L\right]$$

The myofibril composed of 2 hs in series behaves like a linear spring.

Note: $\nu > 0 \quad \Rightarrow \quad |\Delta X_1| < |\Delta X_2|$

The shortening value for the most proximal hs ($\Delta X_1$) is equal to:

$$\Delta X_1 = \frac{1}{2 + \frac{\nu}{\chi}} \cdot \Delta L$$

For the most distal hs ($\Delta X_2$):

$$\Delta X_2 = \frac{1 + \frac{\nu}{\chi}}{2 + \frac{\nu}{\chi}} \cdot \Delta L$$

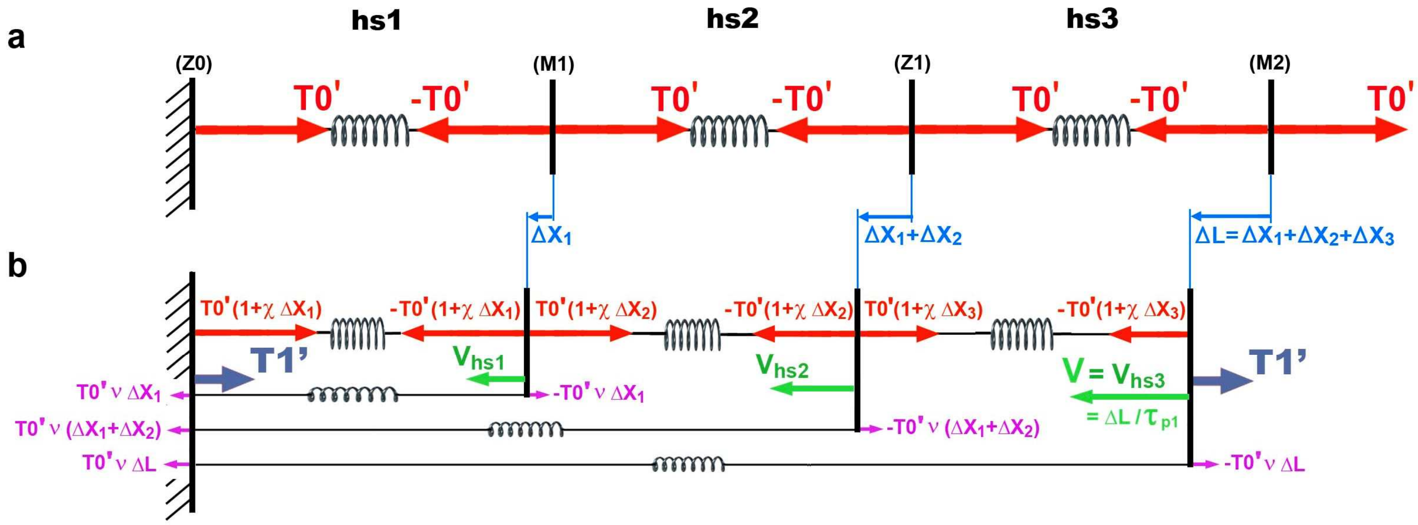


**Fig J1. Modeling of a myofibril composed of 3 hs before and after a shortening.**

(a) Myofibril during the isometric tetanus plateau prior to shortening.

(b) Myofibril at the end of the constant velocity shortening of a length step, i.e. at the end of phase 1.

##### *J.3.3 n= 3 with $\Delta L = \Delta X_1+\Delta X_2+\Delta X_3$*

At the end of phase 1, hs n° 1, 2 and 3 are shortened by $\Delta X_1$, $\Delta X_2$ and $\Delta X_3$ in Zone 1 (Fig J1b).

*LMP* to {$M_1$+2Mfil}: $0 = -T0' \cdot (1+\chi \cdot \Delta X_1) + T0' \cdot (1+\chi \cdot \Delta X_2) - T0' \cdot \nu \cdot \Delta X_1$

$$\Rightarrow \quad \Delta X_2 = \left(1+\frac{\nu}{\chi}\right) \cdot \Delta X_1$$

$$\Rightarrow \quad \Delta X_1 + \Delta X_2 = \left(2+\frac{\nu}{\chi}\right) \cdot \Delta X_1$$

*LMP* to {$Z_1$+2Afil}: $0 = -(1+\chi \cdot \Delta X_2) + (1+\chi \cdot \Delta X_3) - \nu \cdot (\Delta X_1 + \Delta X_2)$

$$\Rightarrow \quad \Delta X_3 = \Delta X_2 + \frac{\nu}{\chi} \cdot (\Delta X_1 + \Delta X_2) = \left(1+\frac{3\nu}{\chi}+\frac{\nu^2}{\chi^2}\right) \cdot \Delta X_1$$

$$\Rightarrow \quad \Delta L = \Delta X_1 + \Delta X_2 + \Delta X_3 = \left(3+\frac{4\nu}{\chi}+\frac{\nu^2}{\chi^2}\right) \cdot \Delta X_1$$

*LMP* to {$M_2$+Mfil}: $0 = T1'/T0' - (1+\chi \cdot \Delta X_3) - \nu \cdot \Delta L$

$$\Rightarrow \quad \frac{T1'}{T0'} = [1+\chi \cdot \Delta X_3 + \nu \cdot \Delta L] = \left[1+\left(\chi \cdot \frac{1+\frac{3\nu}{\chi}+\frac{\nu^2}{\chi^2}}{3+\frac{4\nu}{\chi}+\frac{\nu^2}{\chi^2}} + \nu\right) \cdot \Delta L\right]$$

The myofibril composed of 3 hs in series behaves like a linear spring (Fig J1b).

Note: $\nu > 0 \;\; \Rightarrow \;\; |\Delta X_1| < |\Delta X_2| < |\Delta X_3|$

The shortening value for the most proximal hs ($\Delta X_1$) is equal to:

$$\Delta X_1 = \frac{1}{3+\frac{4\nu}{\chi}+\frac{\nu^2}{\chi^2}} \cdot \Delta L$$

For the most distal hs ($\Delta X_3$):

$$\Delta X_3 = \frac{1+\frac{3\nu}{\chi}+\frac{\nu^2}{\chi^2}}{3+\frac{4\nu}{\chi}+\frac{\nu^2}{\chi^2}} \cdot \Delta L$$

##### *J.3.4 n= 4 with ΔL = Δ X₁+ΔX₂+ΔX₃+ΔX₄*

By applying the *LMP* at the end of phase 1 to the 4 solids, {$M_1$+2Mfil}, {$Z_1$+2Afil}, {$M_2$+2Mfil} and {$Z_2$+Afil}, we obtain, according to the same reasoning as the 3 previous cases:

$$\frac{T1'}{T0'} = \left[1+\chi\cdot\Delta X_4+\nu\cdot\Delta L\right] = \left[1+\left(\chi\cdot\frac{1+6\frac{\nu}{\chi}+5\frac{\nu^2}{\chi^2}+\frac{\nu^3}{\chi^3}}{4+10\frac{\nu}{\chi}+6\frac{\nu^2}{\chi^2}+\frac{\nu^3}{\chi^3}}+\nu\right)\cdot\Delta L\right]$$

The myofibril composed of 4 hs in series behaves like a linear spring.

Note: $\nu > 0 \Rightarrow |\Delta X_1| < |\Delta X_2| < |\Delta X_3| < |\Delta X_4|$

The shortening value for the most proximal hs ($\Delta X_1$) is equal to:

$$\Delta X_1 = \frac{1}{4+10\frac{\nu}{\chi}+6\frac{\nu^2}{\chi^2}+\frac{\nu^3}{\chi^3}}\cdot\Delta L$$

For the most distal hs ($\Delta X_4$):

$$\Delta X_4 = \frac{1+6\frac{\nu}{\chi}+5\frac{\nu^2}{\chi^2}+\frac{\nu^3}{\chi^3}}{4+10\frac{\nu}{\chi}+6\frac{\nu^2}{\chi^2}+\frac{\nu^3}{\chi^3}}\cdot\Delta L$$

##### *J.3.5 n= 5 with ΔL = ΔX₁+ΔX₂+ΔX₃+ΔX₄+ΔX₅*

By applying the *LMP* at the end of Phase 1 to the 5 solids, {{$M_1$+2Mfil}, {$Z_1$+2Afil}, {$M_2$+2Mfil}, {$Z_2$+2Afil} and {$M_3$+Mfil}, we obtain:

$$\frac{T1'}{T0'} = [1+\chi\cdot\Delta X_5 + \nu\cdot\Delta L] = \left[1+\left(\chi\cdot\frac{1+10\frac{\nu}{\chi}+15\frac{\nu^2}{\chi^2}+7\frac{\nu^3}{\chi^3}+\frac{\nu^4}{\chi^4}}{5+20\frac{\nu}{\chi}+21\frac{\nu^2}{\chi^2}+8\frac{\nu^3}{\chi^3}+\frac{\nu^4}{\chi^4}}+\nu\right)\right]\cdot\Delta L$$

The myofibril composed of 5 hs in series behaves like a linear spring.

Note: $\nu > 0 \Rightarrow |\Delta X_1| < |\Delta X_2| < |\Delta X_3| < |\Delta X_4| < |\Delta X_5|$

The shortening value for the most proximal hs ($\Delta X_1$) is equal to:

$$\Delta X_1 = \frac{1}{5+20\frac{\nu}{\chi}+21\frac{\nu^2}{\chi^2}+8\frac{\nu^3}{\chi^3}+\frac{\nu^4}{\chi^4}}\cdot\Delta L$$

For the most distal hs ($\Delta X_5$):

$$\Delta X_5 = \frac{1+10\frac{\nu}{\chi}+15\frac{\nu^2}{\chi^2}+7\frac{\nu^3}{\chi^3}+\frac{\nu^4}{\chi^4}}{5+20\frac{\nu}{\chi}+21\frac{\nu^2}{\chi^2}+8\frac{\nu^3}{\chi^3}+\frac{\nu^4}{\chi^4}}\cdot\Delta L$$

##### *J.3.6 n= 6 with ΔL = ΔX₁+ΔX₂+ΔX₃+ΔX₄+ΔX₅ +ΔX₅*

By applying the *LMP* at the end of phase 1 to the 6 solids {$M_1$+2Mfil}, {$Z_1$+2Afil}, {$M_2$+2Mfil}, {$Z_2$+2Afil}, {$M_3$+2Mfil} and {$Z_3$+Afil}, we obtain:

$$\frac{T1'}{T0'} = [1+\chi\cdot\Delta X_6 + \nu\cdot\Delta L] = \left[1+\left(\chi\cdot\frac{1+15\frac{\nu}{\chi}+35\frac{\nu^2}{\chi^2}+28\frac{\nu^3}{\chi^3}+9\frac{\nu^4}{\chi^4}+\frac{\nu^5}{\chi^5}}{6+35\frac{\nu}{\chi}+56\frac{\nu^2}{\chi^2}+36\frac{\nu^3}{\chi^3}+10\frac{\nu^4}{\chi^4}+\frac{\nu^5}{\chi^5}}+\nu\right)\cdot\Delta L\right]$$

The myofibril composed of 6 hs in series behaves like a linear spring.

Note: $\nu > 0 \;\Rightarrow\; |\Delta X_1| < |\Delta X_2| < |\Delta X_3| < |\Delta X_4| < |\Delta X_5| < |\Delta X_6|$

The shortening value for the most proximal hs ($\Delta X_1$) is equal to:

$$\Delta X_1 = \frac{1}{6+35\frac{\nu}{\chi}+56\frac{\nu^2}{\chi^2}+36\frac{\nu^3}{\chi^3}+10\frac{\nu^4}{\chi^4}+\frac{\nu^5}{\chi^5}}\cdot\Delta L$$

For the most distal hs ($\Delta X_6$):

$$\Delta X_6 = \frac{1+15\frac{\nu}{\chi}+35\frac{\nu^2}{\chi^2}+28\frac{\nu^3}{\chi^3}+9\frac{\nu^4}{\chi^4}+\frac{\nu^5}{\chi^5}}{6+35\frac{\nu}{\chi}+56\frac{\nu^2}{\chi^2}+36\frac{\nu^3}{\chi^3}+10\frac{\nu^4}{\chi^4}+\frac{\nu^5}{\chi^5}}\cdot\Delta L$$

***J.3.7 General term n with $\Delta L = \Delta X_1+\Delta X_2+\Delta X_3+ \ldots +\Delta X_n$***

At the end of phase 1, the n hs are shortened by $\Delta X_1$, $\Delta X_2$, $\Delta X_3$, … , $\Delta X_n$, respectively.

By recurrence, we obtain:

$$\frac{T1'}{T0'} = \left[1+\left(\chi\cdot\underbrace{\frac{1+N_{1,n}\frac{\nu}{\chi}+N_{2,n}\frac{\nu^2}{\chi^2}+N_{3,n}\frac{\nu^3}{\chi^3}+\ldots+N_{n-1,n}\frac{\nu^{n-1}}{\chi^{n-1}}}{n+D_{1,n}\frac{\nu}{\chi}+D_{2,n}\frac{\nu^2}{\chi^2}+D_{3,n}\frac{\nu^3}{\chi^3}+\ldots+D_{n-1,n}\frac{\nu^{n-1}}{\chi^{n-1}}}}_{R_{\chi,n}}+\nu\right)\cdot\Delta L\right] \tag{J10}$$

where the n coefficients to the numerator of $R_{\chi,n}$ are calculated according to:

$$N_{0,n} = 1 \tag{J11a}$$

$$N_{1,n} = \frac{n\cdot(n-1)}{2!} \tag{J11b}$$

$$N_{2,n} = N_{2,n-1} + D_{1,n-1} = \frac{(n-2)\cdot(n-1)\cdot n\cdot(n+1)}{4!} \tag{J11c}$$

$$N_{3,n} = N_{3,n-1} + D_{2,n-1}$$

$$\ldots$$

$$N_{k,n} = N_{k,n-1} + D_{k-1,n-1}$$

$$\ldots$$

$$N_{n-1,n} = 1$$

and where the n coefficients to the denominator of $R_{\chi,n}$ are formulated :

$$D_{0,n} = n \tag{J12a}$$

$$D_{1,n} = D_{1,n-1} + N_{1,n} = \frac{n\cdot(n^2-1)}{3!} \tag{J12b}$$

$$D_{2,n} = D_{2,n-1} + N_{2,n} = \frac{(n-2)\cdot(n-1)\cdot n\cdot(n^2+3n+2)}{5!} \tag{J12c}$$

$$D_{3,n} = D_{3,n-1} + N_{3,n}$$

.

$$\ldots$$

$$D_{k,n} = D_{k,n-1} + N_{k,n}$$

$$\ldots$$

$$D_{n-1,n} = 1$$

The myofibril composed of n hs in series behaves like a linear spring.

The shortening value for the most proximal hs ($\Delta X_1$) is equal to:

$$\Delta X_1 = \frac{1}{n + D_{1,n}\frac{\nu}{\chi} + D_{2,n}\frac{\nu^2}{\chi^2} + D_{3,n}\frac{\nu^3}{\chi^3} + ... + D_{n-1,n}\frac{\nu^{n-1}}{\chi^{n-1}}} \cdot \Delta L \quad \text{(J13a)}$$

The shortening value for the next hs ($\Delta X_2$) is equal to:

$$\Delta X_2 = \frac{1 + N_{1,n}\frac{\nu}{\chi}}{n + D_{1,n}\frac{\nu}{\chi} + D_{2,n}\frac{\nu^2}{\chi^2} + D_{3,n}\frac{\nu^3}{\chi^3} + ... + D_{n-1,n}\frac{\nu^{n-1}}{\chi^{n-1}}} \cdot \Delta L$$

The shortening value for the next hs ($\Delta X_3$) is equal to:

$$\Delta X_3 = \frac{1 + N_{1,n}\frac{\nu}{\chi} + N_{2,n}\frac{\nu^2}{\chi^2}}{n + D_{1,n}\frac{\nu}{\chi} + D_{2,n}\frac{\nu^2}{\chi^2} + D_{3,n}\frac{\nu^3}{\chi^3} + ... + D_{n-1,n}\frac{\nu^{n-1}}{\chi^{n-1}}} \cdot \Delta L$$

And so on until the most distal hs ($\Delta X_n$):

$$\Delta X_n = R_{\chi,n} \cdot \Delta L \quad \text{(J13b)}$$

In the presence of viscosity ($\nu > 0$), we check:

$$|\Delta X_1| < |\Delta X_2| < |\Delta X_3| < \ldots < |\Delta X_k| < \ldots < |\Delta X_{n-1}| < |\Delta X_n| \quad \text{(J14)}$$

**Remarks**

$N_{1,n}$ formulated in (J11b) is the sum of the first (n-1) integers, i.e. a polynomial of degree 2, and $D_{1,n}$ formulated by (J12b) is the sum of the sum of the first (n-1) integers, i.e. a polynomial of degree 3.

$N_{2,n}$ formulated in (J11c) is the sum of the sum of the sum of the first (n-2) integers, i.e. a polynomial of degree 4, and $D_{2,n}$ formulated by (J12c) is the sum of the sum of the sum of the sum of the first (n-2) integers, i.e. a polynomial of degree 5.

$N_{3,n}$ is the sum of the sum of the sum of the sum of the sum of the first (n-3) integers, i.e. a polynomial of degree 6, and $D_{3,n}$ is the sum of the sum of the sum of the sum of the sum of the sum of the first (n-3) integers, i.e. a polynomial of degree 7.

By recurrence, we deduce that $N_{k,n}$ is a polynomial of degree (2k) and that $D_{k,n}$ is a polynomial of degree (2k+1).

***J.3.8 Cases where n is large (n>250)***

According to the conclusions of the previous paragraph, when n becomes large (n > 250), there is an integer bound, noted m, close to n/2 such that the coefficients $N_{k,n}$ and $D_{k,n}$ with $1 \le k \le m$ are approached by powers of n, and such that the following coefficients $N_{k,n}$ and $D_{k,n}$ with $m < k \le n$ are considered negligible.

So:

$$N_{1,n} \approx \frac{n^2}{2!}$$

$$N_{2,n} \approx \frac{n^4}{4!}$$

$$\dots$$

$$N_{m,n} \approx \frac{n^{2m}}{(2m)!}$$

$$D_{1,n} \approx \frac{n^3}{3!}$$

$$D_{2,n} \approx \frac{n^5}{5!}$$

$$\dots$$

$$D_{m,n} \approx \frac{n^{2m+1}}{(2m+1)!}$$

In the presence of viscosity ($\nu$>0), we propose to express the ratio $\nu/\chi$ in the form of a negative power of n:

$$\frac{\nu}{\chi} = n^{-q} \qquad \text{(J15)}$$

where q is a real positive.

By replacing the ratio $\nu/\chi$ by $n^{-q}$ and the integer m by n/2, equality (J.10) is reformulated with the previous approximations:

$$\frac{T1'}{T0'} \approx \left[ 1 + \left( \chi \cdot \frac{1 + \frac{n^{(2-q)}}{2!} + \frac{n^{2(2-q)}}{4!} + \frac{n^{3(2-q)}}{6!} + \dots + \frac{n^{n(2-q)}}{(n)!}}{n + \frac{n^{(2-q)+1}}{3!} + \frac{n^{2(2-q)+1}}{5!} + \frac{n^{3(2-q)+1}}{7!} + \dots + \frac{n^{n(2-q)+1}}{(n+1)!}} + \nu \right) \cdot \Delta L \right] \qquad \text{(J16)}$$

The entire series developments of the hyperbolic sine and cosine are written:

$$ch(x) = \frac{e^{x} + e^{-x}}{2} = 1 + \frac{x^{2}}{2!} + \frac{x^{4}}{4!} + \frac{x^{6}}{6!} + ... + \frac{x^{k}}{(n)!} + ....$$

$$sh(x) = \frac{e^{x} - e^{-x}}{2} = x + \frac{x^{3}}{3!} + \frac{x^{5}}{5!} + \frac{x^{7}}{7!} + ... + \frac{x^{k+1}}{(k+1)!} + ....$$

The hyperbolic cotangent function is classically defined:

$$coth(x) = \frac{ch(x)}{sh(x)} = \frac{e^{2x} + 1}{e^{2x} - 1}$$

A new formulation of (J10) and (J16) is deduced from the previous expressions; so with (J15):

$$\frac{T1'}{T0'} \approx \left[ 1 + \left( \chi \cdot \underbrace{\left[ \frac{1}{n} \cdot n^{1-\frac{q}{2}} \cdot coth\left( n^{1-\frac{q}{2}} \right) \right]}_{Q_{q,n}} + \nu \right) \cdot \Delta L \right] \quad (J17)$$

The coefficient $R_{\chi,n}$ determined in (J.10) is approximated by the parameter $Q_{q,n}$ characterized in (J.17):

$$Q_{q,n} = \frac{1}{n} \cdot n^{1-\frac{q}{2}} \cdot coth\left( n^{1-\frac{q}{2}} \right) \approx R_{\chi,n} \quad (J18)$$

The equality (J18) is tested for 3 values of q (Fig J2). All numerical simulations where the coefficient $R_{\chi,n}$ is compared to $Q_{q,n}$ validate (J18).

Parameter K is introduced as follows:

$$K = n^{1-q/2} \cdot coth\left( n^{1-q/2} \right) + n^{1-q} \quad (J19)$$

The equations (J15) and (J19) allow the rewriting of (J17):

$$\frac{T1'}{T0'} \approx \left( 1 + K \cdot \chi \cdot \frac{\Delta L}{n} \right) \quad (J20)$$

By definition, the average shortening ($\overline{\Delta X}$) is equal to:

$$\overline{\Delta X} = \frac{\Delta L}{N_{hs}} \quad (J21)$$

where $N_{hs}$ is the number of hs per myofibril.

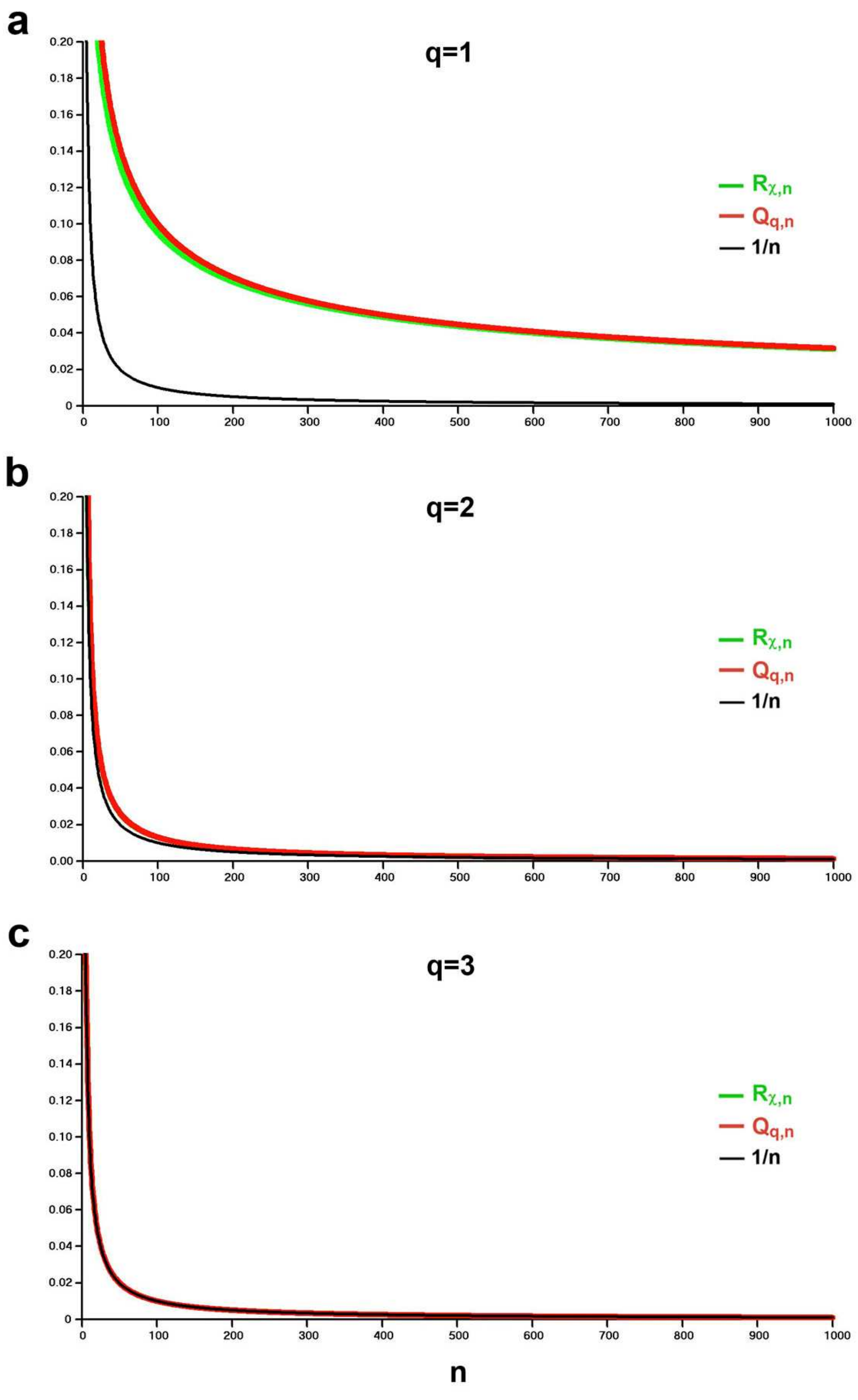


**Fig J2. Approximation of $R_{\chi,n}$ par $Q_{q,n}$ where n is the number of hs, varying from 1 to 1000.**
(a) q=1. (b) q=2. (c) q=3.

Since this paragraph is specifically devoted to Zone 1, q and K are renamed to $q_{z1}$ and $K_{z1}$. Returning to the original ratings $\chi_{z1}$, $T0_m$ and $T1_m$ instead of $\chi$, T0' and T1', the relative tension at the end of phase 1 (pT1) is rewritten according to (J20) and (J21) with n= $N_{hs}$:

$$pT1 = \frac{T1}{T0} = \frac{T1_m}{T0_m} \approx \left[1 + \left(\chi_{z1} \cdot K_{z1}\right) \cdot \overline{\Delta X}\right] \qquad (J22)$$

where T1 is the tension applied to the two extremities of the fiber at the end of phase 1; T0 is the tension of the fiber during the isometric tetanus plateau ; $K_{z1}$ is a multiplier coefficient greater than 1 calculated from (J19) with $n=N_{hs}$:

$$K_{z1} = N_{hs}^{1-q_{z1}/2} \cdot \coth\left(N_{hs}^{1-q_{z1}/2}\right) + N_{hs}^{1-q_{z1}} \qquad (J23)$$

The parameter $K_{z1}$ depends mainly on 2 factors: the number of hs per myofibril ($N_{hs}$) and the viscosity through the coefficient $q_{z1}$ determined from the equality (J15) according to:

$$q_{z1} = \frac{Ln\chi_{z1} - Ln\nu}{LnN_{hs}} \qquad (J24)$$

Figure J3 shows 6 plots of the coefficient K (or $K_{z1}$) as a function of the real exponent q (or $q_{z1}$) for 6 values of $N_{hs}$. The 6 plots pass through the coordinate point (q=2; K =1.31). In the absence of viscosity, i.e. as soon as $q \geq 2.3$, the coefficient K tends towards 1.

**Note**: Most experiments verify the conditions: $N_{hs} \geq 4500$ and $q_{z1} \geq 1.7$; in this case, the factor $N_{hs}^{1-q_{z1}}$ present in the right member of (J23) becomes negligible. It will nevertheless be kept in all our algorithmic calculations.

##### *J.3.9 Inequality of hs shortening in the presence of viscosity*

The series of inequalities given in (J14) indicate that in the presence of viscosity, the hs shortening increases in modulus from the fixed end of the myofibril (h=1) to the movable end ($h=N_{hs}$). By recurrence, the n coefficients ($N_{h-1,n}$) present to the numerator of the parameter $R_{\chi,n}$ introduced in (J10) to sub-paragraph J.3.7 are calculated algorithmically, with h as an integer varying from 1 to n; the first three values, $N_{0,n}$, $N_{1,n}$ and $N_{2,n}$, are written in (J11a), (J11b) and (J11c) and the recurrence formulas of the following coefficients appear thereafter. The n coefficients ($D_{h-1,n}$) present to the denominator of $R_{\chi,n}$ are calculated in a similar way: the first three values, $D_{0,n}$, $D_{1,n}$ and $D_{2,n}$, are formulated in (J12a), (J12b) and (J12c) and the recurrence formulas of the following coefficients appear next.

These data make it possible to determine the n shortenings of the n hs of a myofibril; see expressions of $\Delta X_1$ and $\Delta X_n$ with (J13a) and (J13b) and those of $\Delta X_2$ and $\Delta X_3$ between (J13a) and (J13b).

In Fig J3 we present 3 evolutions of the hs shortening relative to the fiber shortening for 3 values of $q_{z1}$ in the particular case where $n=N_{hs}=1000$.

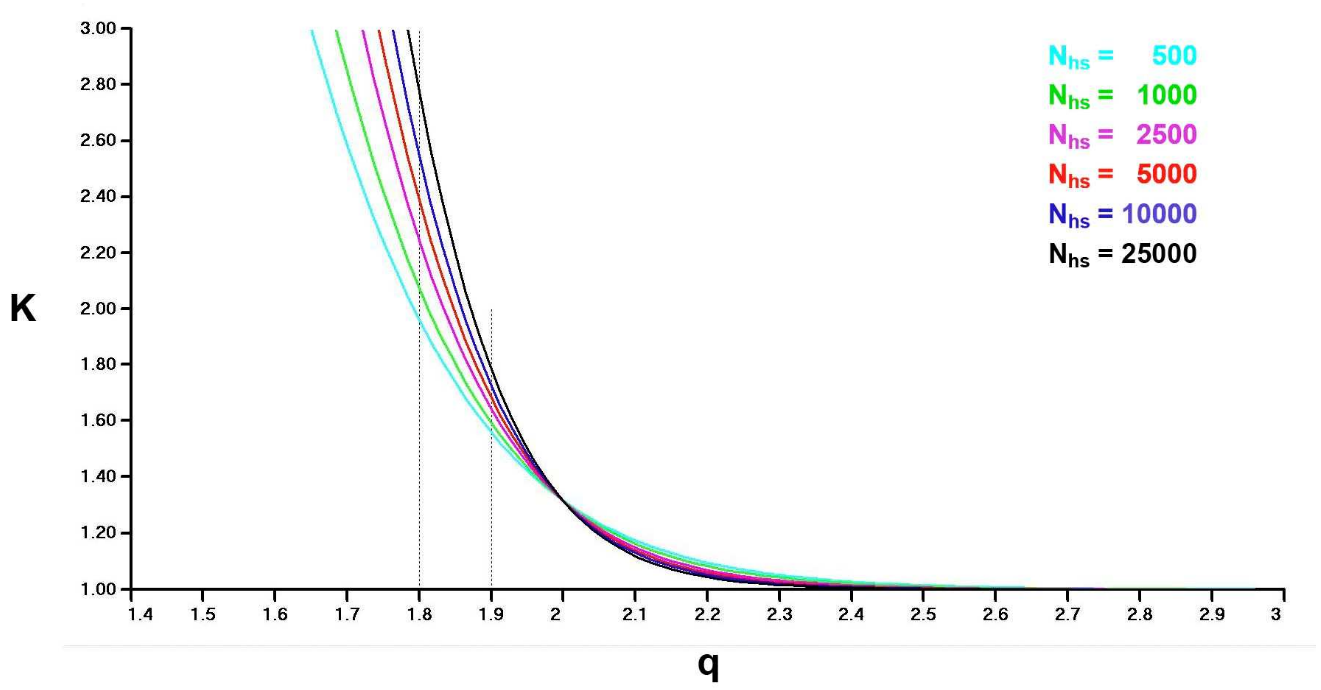


**Fig J3. Curves of the multiplier coefficient K determined in (J19) as a function of the viscous parameter q for 6 values of the number of half-sarcomeres (n=$N_{hs}$).**

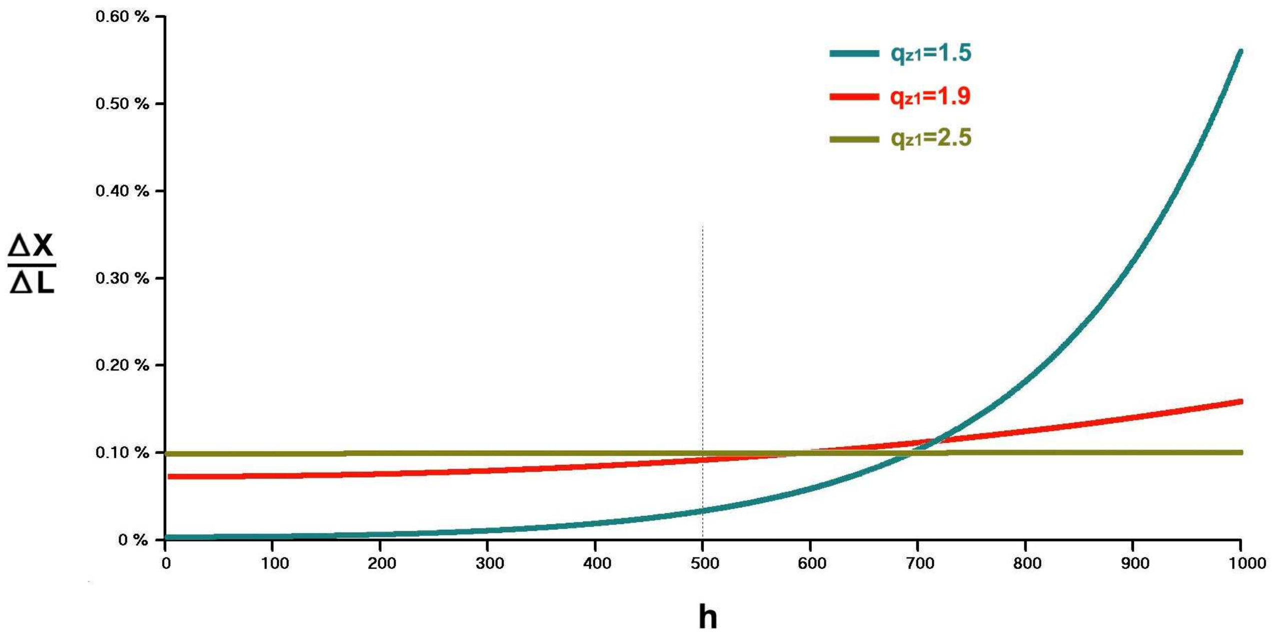


**Fig J4. Evolution of the relative shortening of hs n° h ($\Delta X_h/\Delta L$) with the index h incremented from 1 to 1000 for 3 values of $q_{z1}$; $\Delta L$ is the shortening of the fiber.**

The three cases in Fig J4 are analyzed:

**1/ $q_{z1}$=2.5** (khaki line): value greater than 2.3, the lower bound given as a condition of the absence of viscosity; all shortenings are equal at the end of phase 1 according to the relationship (J9). As soon as $q_{z1} \geq 2.3$, $K_{z1}$ tends towards 1 and the linear equation (J22) tends towards the linear equation (J7) where viscosity is nil.

**2/ $q_{z1}$=1.9** (red line): the influence of viscosity is significant, but all hs shorten with a value that does not deviate significantly from the average value $\overline{\Delta X}$.

**3/ $q_{z1}$=1.5** (blue line): the differences are very marked; the proximal hs of numbers 1 to 100 have zero shortening while the distal hs of numbers 900 to 1000 are shortened with a length 3 to 5 times greater than the average value. The distal hs of n° 600 to 1000 receive 90% of the total shortening (ΔL).

##### *J.3.10 Zone 1 TRUE*

With the definition of the hyperbolic cosine, the shortening of the most proximal hs ($\Delta X_1$) is determined from the relationships (J13a), (J18), (J19), (J21) and (J23):

$$\Delta X_1 = \frac{K_{z1}}{ch\left(N_{hs}{}^{1-q_{z1}/2}\right)} \cdot \overline{\Delta X} \tag{J25a}$$

Similarly, the shortening of the most distal hs ($\Delta X_{Nhs}$) is evaluated according to (J13b), (J18), (J19), (J21) and (J23):

$$\Delta X_{N_{hs}} = K_{z1} \cdot \overline{\Delta X} \tag{J25b}$$

The most distal hs is the first one whose shortening passes into Zone 2 and the corresponding mean shortening ($Bz1_{min}$) checks (J25b) such that:

$$\Delta X_{N_{hs}} = -\delta X_{z1} = K_{z1} \cdot Bz1_{min}$$

Consequently:

$$Bz1_{min} = \frac{-\delta X_{z1}}{K_{z1}} \tag{J26a}$$

We define "Zone 1 TRUE" in relation to the average shortening ($\overline{\Delta X}$) when $\overline{\Delta X}$ is between $Bz1_{min}$ and 0 (Fig J5a), case where individual shortenings of the $N_{hs}$ hs of the myofibril are all in Zone 1 (Fig J5b).

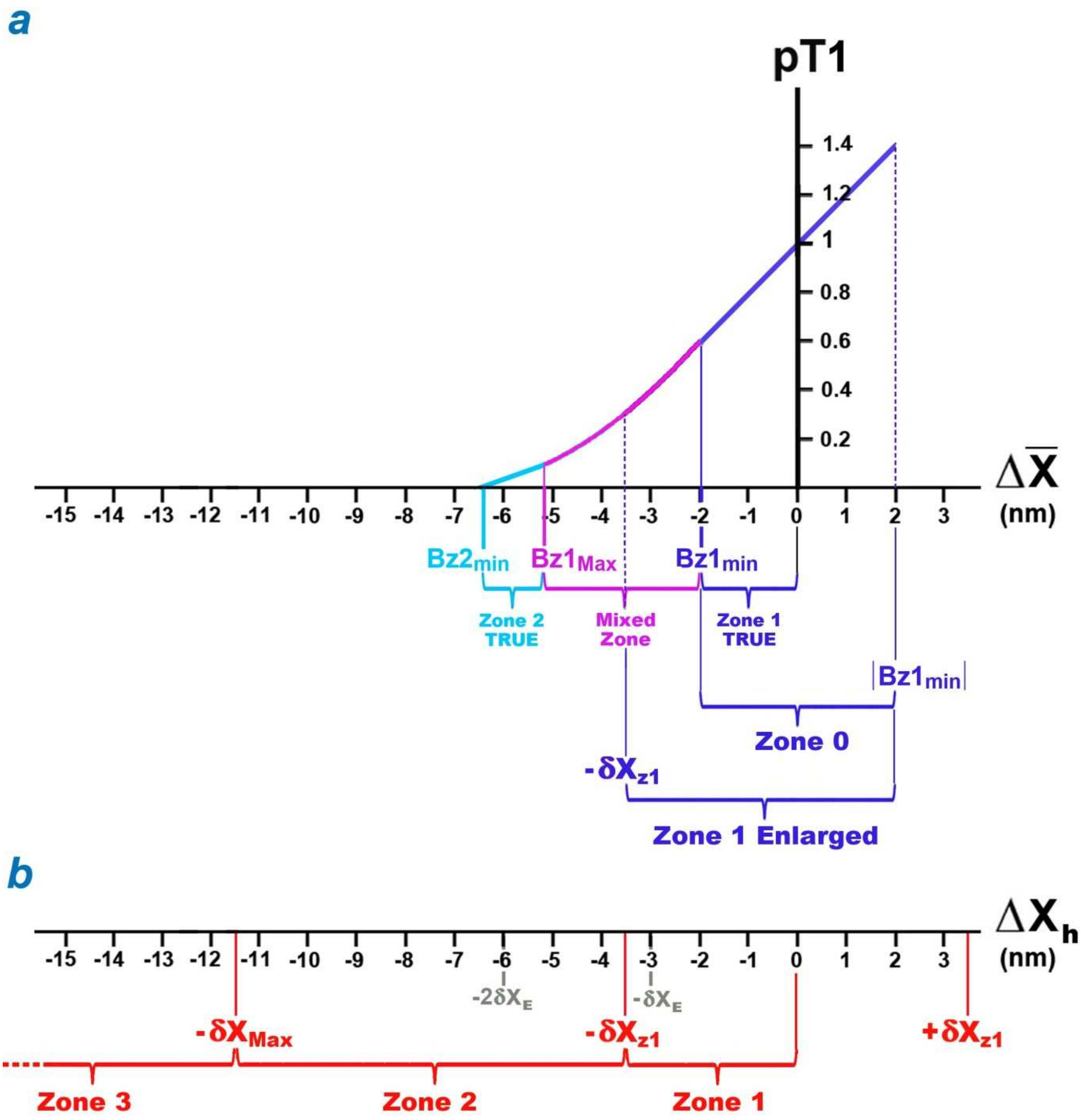


**Fig J5. (a) Characterization of "Zone 1 TRUE", "Zone 2 TRUE", "Mixed Zone", "Zone O" and "Zone 1 Enlarged" in relation to the average shortening ( $\overline{\Delta X}$ ). (b) Reminder of the definitions of Zones 1, 2 and 3 with regard to the shortening of hs n° h ($\Delta X_h$), h integer index varying from 1 to $N_{hs}$.**

The most proximal hs is the last one whose shortening ($\Delta X_1$) passes into Zone 2 and the corresponding mean shortening ($Bz1_{Max}$) checks (J25a) such that:

$$\Delta X_1 = -\delta X_{z1} = \frac{K_{z1}}{ch\left(N_{hs}{}^{1-q_{z1}/2}\right)} \cdot Bz1_{Max}$$

With (J26a), it is deduced:

$$Bz1_{Max} = Bz1_{min} \cdot ch\left(N_{hs}{}^{1-q_{z1}/2}\right) \tag{J26b}$$

We verify:

$$Bz1_{Max} \leq -\delta X_{z1} \leq Bz1_{min} \tag{J26c}$$

In the absence of viscosity ($q_{z1} \geq 2.3$), (J26c) becomes a relationship of strict equalities.

#### J.4 Calculation of the tension at the end of phase 1 in the presence of viscosity when all hs shortenings belong to Zone 2

##### *J.4.1 Recalls and calculations*

***Recalls of results given in paragraph I.6 of Supplement S4.I***

*Zone 2 is defined by the interval [-$\delta X_{Max}$ ; -$\delta X_{z1}$]. In the absence of viscosity, the tension ($T1_{Elas,z2}$) exerted on both edges of a hs comes exclusively from the forces deriving from the elastic energy of the WS myosin heads and is formulated in Zone 2 according to (I53a):*

$$\frac{T1_{Elas,z2}}{T0_m} = \chi_{z2} \cdot \left(\delta X_{Max} + \Delta X\right) \tag{J27}$$

*where $\chi_{z2}$ is the coefficient of elastic origin stiffness in Zone 2 which is written with (I53b):*

$$\chi_{z2} = \frac{\left(1 - \chi_{z1} \cdot \delta X_{z1}\right)}{\delta X_{Max} - \delta X_{z1}} \tag{J28}$$

In the presence of viscosity, for the shortenings of all hs to be in Zone 2, the following condition from (J25b) must be checked:

$$\Delta X_1 \leq Bz1_{Max} \tag{J29}$$

where $\Delta X_1$ is the shortening of the most proximal hs (h=1).

Since the constituent assemblies of a hs are identical, the viscosity is always characterized by the parameter ν defined in (J15).

By adopting a recurrence reasoning similar to that used for shortenings in Zone 1 in paragraph J.3 where equation (J7) is replaced by (J27), we obtain in Zone 2 when the condition (J29) is established:

$$\frac{T1_m}{T0_m} = \left[ \chi_{z2} \cdot \delta X_{Max} + \left( \chi_{z2} \cdot \underbrace{\frac{1 + N_{1,n}\dfrac{\nu}{\chi_{z2}} + N_{2,n}\dfrac{\nu^2}{\chi_{z2}^2} + N_{3,n}\dfrac{\nu^3}{\chi_{z2}^3} + \ldots + N_{n-1,n}\dfrac{\nu^{n-1}}{\chi_{z2}^{n-1}}}{n + D_{1,n}\dfrac{\nu}{\chi_{z2}} + D_{2,n}\dfrac{\nu^2}{\chi_{z2}^2} + D_{3,n}\dfrac{\nu^3}{\chi_{z2}^3} + \ldots + D_{n-1,n}\dfrac{\nu^{n-1}}{\chi_{z2}^{n-1}}}}_{R_{\chi_{z2},n}} + \nu \right) \cdot \Delta L \right]$$

(J30)

where n is the number of hs per myofibril.

The series of inequalities proposed in (J14) is always verified in Zone 2. The value of the shortening of the most proximal hs (h=1) is equal to:

$$\Delta X_1 = \frac{\Delta L}{n + D_{1,n}\dfrac{\nu}{\chi_{z2}} + D_{2,n}\dfrac{\nu^2}{\chi_{z2}^2} + D_{3,n}\dfrac{\nu^3}{\chi_{z2}^3} + \ldots + D_{n-1,n}\dfrac{\nu^{n-1}}{\chi_{z2}^{n-1}}} \qquad \text{(J31a)}$$

And that of the shortening of the most distal hs (h=n) is worth:

$$\Delta X_n = R_{\chi_{z2},n} \cdot \Delta L \qquad \text{(J31b)}$$

##### *J.4.2 Case for n > 250*

The ratio $\nu/\chi_{z2}$ is formulated as a negative power of n:

$$\frac{\nu}{\chi_{z2}} = n^{-q_{z2}} \qquad \text{(J32)}$$

where $q_{z2}$ is a real positive.

By following the inductive method proposed in sub-paragraph J.3.8 for Zone 1, the relative value of the fiber tension at the end of phase 1 is obtained when all shortenings are in Zone 2:

$$pT1 = \frac{T1}{T0} = \frac{T1_m}{T0_m} = \left[\chi_{z2} \cdot \left(\delta X_{Max} + K_{z2} \cdot \overline{\Delta X}\right)\right] \qquad \text{(J33)}$$

where $K_{z2}$ is a multiplier coefficient greater then 1, calculated with $q_{z2}$ according to (J19) exactly as $K_{z1}$ was with $q_{z1}$:

$$K_{z2} = N_{hs}^{1-q_{z2}/2} \cdot \coth\left(N_{hs}^{1-q_{z2}/2}\right) + N_{hs}^{1-q_{z2}} \qquad \text{(J34)}$$

We recall that $\overline{\Delta X}$ is the average shortening defined in (J21).

With $n=N_{hs}$, the parameter $q_{z2}$ is determined according to (J32):

$$q_{z2} = \frac{Ln\chi_{z2} - Ln\nu}{LnN_{hs}} \quad \text{(J35)}$$

Comparing (J35) to (J23), we note:

$$q_{z2} = q_{z1} + \frac{1}{LnN_{hs}} \cdot Ln\frac{\chi_{z2}}{\chi_{z1}} \quad \text{(J36)}$$

The elastic stiffness coefficient $\chi_{z2}$ being inferior to $\chi_{z1}$, the equality (J36) implies that $q_{z2}$ is inferior to $q_{z1}$ and consequently that $K_{z2}$ is superior to $K_{z1}$ (Fig J3). It is concluded that the more the stiffness coefficient $\chi$ resulting from the elasticity of the WS myosin heads decreases, the greater the influence of viscosity is. This finding is verified experimentally and explains why viscosity is considered present when the fiber is at rest and neglected when the fiber is stimulated [1].

##### *J.4.3 Zone 2 TRUE*

By comparing the formulations (J13a) and (J31a), the calculation of the most proximal hs ($\Delta X_1$) is carried out in Zone 2 on the model of the one determined in Zone 1 with (J25a):

$$\Delta X_1 = \left( \frac{K_{z2}}{ch\left(N_{hs}{}^{1-q_{z2}/2}\right)} \right) \cdot \overline{\Delta X} \quad \text{(J37a)}$$

By comparing the formulations (J13b) and (J31b), the calculation of the most proximal hs ($\Delta X_{Nhs}$) is carried out in Zone 2 on the model of the one determined in Zone 1 with (J25b):

$$\Delta X_{N_{hs}} = K_{z2} \cdot \overline{\Delta X} \quad \text{(J37b)}$$

The most distal hs is the first one whose shortening ($\Delta X_{Nhs}$) passes into Zone 3 ($\Delta X < -\delta X_{Max}$) and the corresponding mean shortening ($Bz2_{min}$) checks for equality (J37b), i.e.:

$$\Delta X_{N_{hs}} = -\delta X_{Max} = K_{z2} \cdot Bz2_{min}$$

Consequently:

$$Bz2_{min} = \frac{-\delta X_{Max}}{K_{z2}} \quad \text{(J38a)}$$

We define "Zone 2 TRUE" in relation to the average shortening ($\overline{\Delta X}$) when $\overline{\Delta X}$ is between $Bz2_{min}$ and $Bz1_{Max}$ (Fig J5a), case where individual shortenings of the $N_{hs}$ hs of the myofibril are all in Zone 2 (Fig J5b).

The most proximal hs is the last one whose shortening ($\Delta X_1$) passes into Zone 3 and the corresponding mean shortening ($Bz2_{Max}$) checks for equality (J37a), i.e.:

$$\Delta X_1 = -\delta X_{Max} = \frac{K_{z2}}{ch\left(N_{hs}{}^{1-q_{z2}/2}\right)} \cdot Bz2_{Max}$$

With (J38a), it is deduced:

$$Bz2_{Max} = Bz2_{min} \cdot ch\left(N_{hs}{}^{1-q_{z2}/2}\right) \quad (J38b)$$

We verify:

$$Bz2_{Max} \leq -\delta X_{Max} \leq Bz2_{min} \quad (J39)$$

In the absence of viscosity with $q_{z2} \geq 2.3$, (J39) becomes a relationship of strict equality.

Note that $Bz2_{min}$ is the value of the average shortening ($\overline{\Delta X}$) for which the equation (J33) cancels out, i.e. the tension across the fiber becomes zero (Fig J5a).

The examples given in Fig J4 for $q_{z1}$ are just as valid for $q_{z2}$.

##### *J.4.4 Mixed Zone between Zone 1 True and Zone 2 True*

When the average shortening ($\overline{\Delta X}$) is between the two abscissa $Bz1_{Max}$ and $Bz1_{min}$, calculated in (J26b) and (J26a), the shortenings of the most distal hs have passed into Zone 2 while the shortenings of the other hs are still in Zone 1. The two terminals $Bz1_{Max}$ and $Bz1_{min}$ frame a Mixed Zone between Zone 1 True and Zone 2 True (Fig J5).

A weight ($p_M$) is introduced:

$$p_M = \left(\frac{\overline{\Delta X} - Bz1_{min}}{Bz1_{Max} - Bz1_{min}}\right) \quad (J40)$$

where $\overline{\Delta X}$ is between $Bz1_{Max}$ and $Bz1_{min}$.

In the Mixed Zone, the tension at the end of phase 1 (pT1) is a function of $\overline{\Delta X}$, estimated in first approximation as the barycentre of the two tensions pT1 calculated in Zone 1 True and Zone 2 True according to relationships (J22) and (J33) to which the weights ($1-p_M$) and $p_M$ are assigned, respectively:

$$pT1 = (1 - p_M) \cdot \left[1 + \chi_{z1} \cdot K\chi_{z1} \cdot \overline{\Delta X}\right] + p_M \cdot \chi_{z2} \cdot \left[\delta X_{Max} + K\chi_{z2} \cdot \overline{\Delta X}\right] \quad (J41)$$

The relative tension (pT1) is represented in the Mixed Zone by a purple parabolic arc (Fig J5a).

#### J.5 Calculation of the tension at the end of phase 1 in the presence of viscosity when all elongations and shortenings of hs belong to Zone O

A fiber lengthening ($\Delta L$) is considered such that all the hs lengthenings ($\Delta X_h$) belong to the linear range [0 ; $+\delta X_{z1}$]; see Fig J5b.

At the end of phase 1, the angular position $\theta$ of the lever of a small part of the WS heads is located beyond the maximum position $\theta_{up}$. We postulate that the linear contribution to the $T1_{Elas}$ tension of a hs is compensated by the linear elastic stretching of the rigid segment S2 and it is assumed that:

$$\chi_{stretch} \approx \chi_{z1} \tag{J42}$$

By using the same ratings as in the previous paragraph, we directly study the case n=3 with:

$$\Delta L = \Delta X_1 + \Delta X_2 + \Delta X_3$$

At the end of phase 1, hs n° 1, 2 and 3 are extended by $\Delta X_1$, $\Delta X_2$ and $\Delta X_3$ (Fig J6). The linear moment principle (*LMP*) is applied to the 3 solids {$M_1$+2Mfil}, {$Z_1$+2Afil) and {$M_2$+Mfil}:

*LMP* to {$M_1$+2Mfil}: $0 = -(1+\chi\cdot\Delta X_1)+(1+\chi\cdot\Delta X_2)-\nu\cdot\Delta X_1$

*LMP* to {$Z_1$+2Afil): $0 = -(1+\chi\cdot\Delta X_2)+(1+\chi\cdot\Delta X_3)-\nu\cdot(\Delta X_1+\Delta X_2)$

*LMP* to {$M_2$+Mfil}: $0 = T'/T0' - (1+\chi\cdot\Delta X_3)-\nu\cdot\Delta L$

The equations being similar to that of sub-paragraph J.2.3, they lead to the same result:

$$\frac{T'}{T0'} = [1+\chi\cdot\Delta X_3+\nu\cdot\Delta L] = \left[1+\left(\chi\cdot\frac{1+\frac{3\nu}{\chi}+\frac{\nu^2}{\chi^2}}{3+\frac{4\nu}{\chi}+\frac{\nu^2}{\chi^2}}+\nu\right)\cdot\Delta L\right]$$

The myofibril composed of 3 hs in series behaves like a linear spring with a coefficient identical to that determined for the shortening of the case n=3 studied in sub-paragraph J.3.3.

The general case leads by recurrence to the same equation formulated in (J.10). The set of inequalities presented in (J14) remains valid. It can be deduced that all equations or equalities from (J15) to (J24) apply. The relative tension exerted at the ends of the fiber or a myofibril is formulated:

$$pT1 = \frac{T1_m}{T0_m} = \frac{T1_m}{T0_m} \approx \left[1+(\chi_{z1}\cdot K_{z1})\cdot\overline{\Delta X}\right] \tag{J43}$$

where $K_{z1}$ is determined in (J23); $\overline{\Delta X}$ is the average shortening or elongation defined algebraically in (J21).

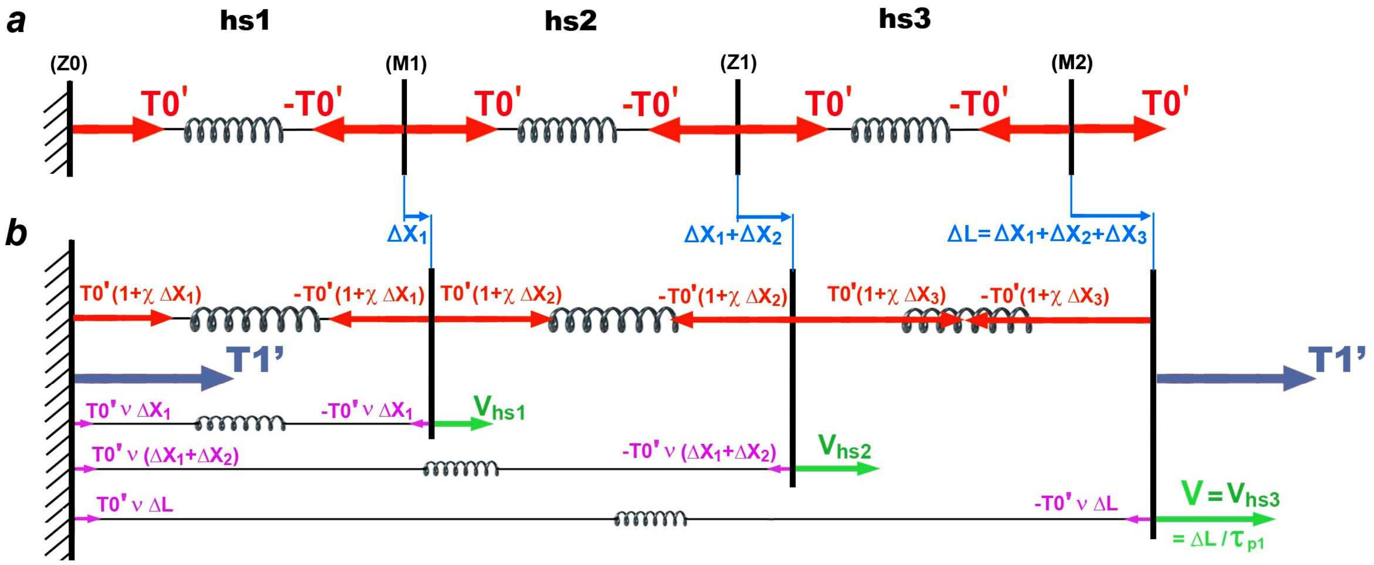


**Fig J6. Modeling of a myofibril composed of 3 hs before and after a lengthening ($\Delta L > 0$).**

(a) Myofibril during the isometric tetanus plateau before elongation.

(b) Myofibril at the end of the constant velocity elongation of a length step, i.e. at the end of phase 1.

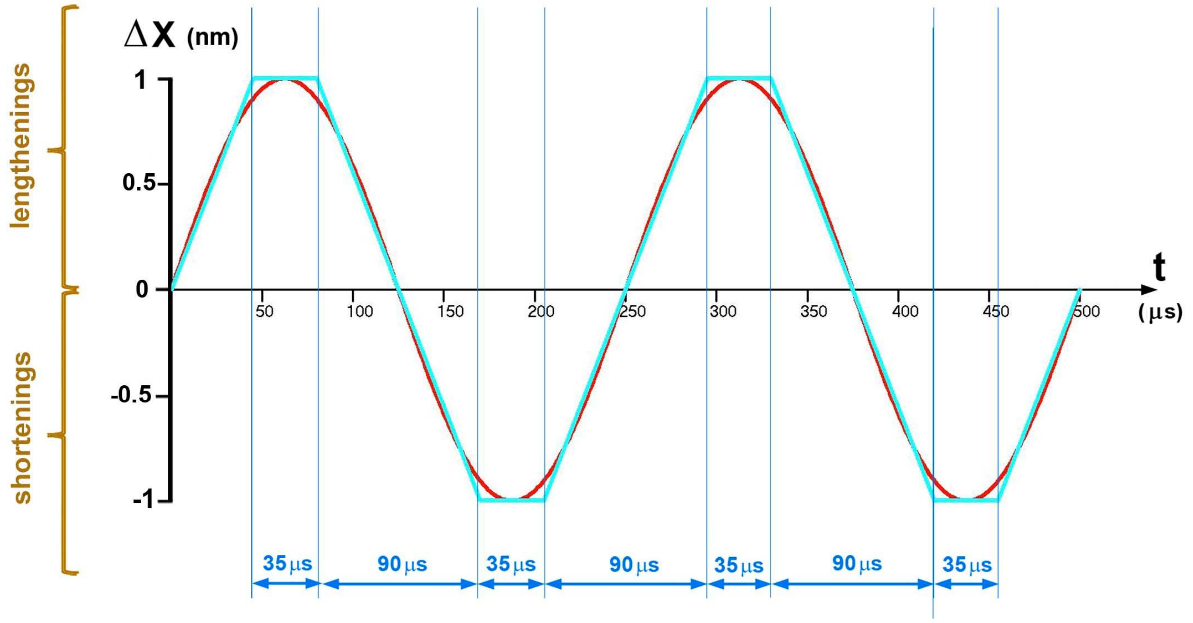


**Fig J7. Modeling of a sinusoid with an amplitude of 2 nm peak to peak and a frequency of 4 kHz (red line) by a succession of straight line segments (light blue line).**

The most distal hs (h=$N_{hs}$) is the first one whose elongation ($\Delta X_{Nhs}$) becomes greater than $+\delta X_{z1}$. The average lengthening ($Sz1_{min}$) corresponding to this value checks:

$$\Delta X_{Nhs} = +\delta X_{z1} = K_{z1} \cdot Sz1_{min}$$

Consequently:

$$Sz1_{min} = \frac{\delta X_{z1}}{K_{z1}} \qquad \text{(J44)}$$

It is noted with (J25a):

$$Sz1_{min} = |Bz1_{min}| \qquad \text{(J45)}$$

Equations (J22) and (J43) are identical. The "Zone O" is defined in relation to the average of the hs length change ($\overline{\Delta X}$) when $\overline{\Delta X}$ is between $Bz1_{min}$ and $|Bz1_{min}|$ (Fig J5a), where the shortenings of the $N_{hs}$ hs are all in the interval [$-\delta X_{z1}$ ; 0] characterizing Zone 1 and where the elongations of the $N_{hs}$ hs are all in the interval [0 ; $+\delta X_{z1}$] (Fig J5b).

The letter O corresponds to the first letter of the word "oscillation". One type of experimentation consists in subjecting the muscle fiber to forced sinusoidal movements of constant amplitude and frequency. As a general rule, the peak-to-peak amplitude is equal to or less than 2 nm and belongs to the Zone O. A sinusoid with a frequency of 4 kHz can be modelled by straight line segments of the same amplitude over a period of 90 µs, spaced by time plateaus of 35 µs (Fig J7). We thus obtain a succession of phases 1 of length steps whose duration is a constant equal to:

$$\tau_{p1,4KHz} \approx 90\ \mu s \qquad \text{(J46)}$$

It becomes possible to calculate pT1 with the equation given in (J22) and (J43) for any non-zero slope segment modelling the sinusoid with T0 as the reference tension equal to the tension in the absence of oscillations, on the one hand, and to the mean tension after rectification in the presence of oscillations, on the other hand; see Fig 1 in [2].

For a frequency of 2.5 kHz, it is established:

$$\tau_{p1,2.5KHz} \approx 135\ \mu s \qquad \text{(J47)}$$

#### J.6 Application with study of an example

The relative tension (pT1) exerted at the extremidties of the fiber at the end of phase 1 and calculated in (J22), (J33), (J41) and (J43) is concisely formulated as a function of $\overline{\Delta X} \in \left[Bz2_{min}; \left|Bz1_{min}\right|\right]$:

$$pT1 = \left[1 - P_{Elas+Visc}\right] \cdot \left(1 + \chi_{z1} \cdot K_{z1} \cdot \overline{\Delta X}\right) + P_{Elas+Visc} \cdot \left[\chi_{z2} \cdot \left(\delta X_{Max} + K_{z2} \cdot \overline{\Delta X}\right)\right] \qquad (J48)$$

where $P_{Elas+Visc}$ is a coefficient varying between 0 and 1 such that :

$$P_{Elas+Visc} = \mathit{1}_{[Bz2_{min};Bz1_{Max}[}\left(\overline{\Delta X}\right) + \left(\frac{\overline{\Delta X} - Bz1_{min}}{Bz1_{Max} - Bz1_{min}}\right) \cdot \mathit{1}_{[Bz1_{Max};Bz1_{min}]}\left(\overline{\Delta X}\right) \qquad (J49)$$

**Method of calculation**

With the values of the first column (Ford 1977) of Table 1 to Paper 4, we determine $\chi_{z1}$ and $\chi_{z2}$, the two elastic origin slopes, according to (J8) and (J28) as:

$$\chi_{z1} = 0.133 \text{ nm}^{-1}$$

$$\chi_{z2} = 0.062 \text{ nm}^{-1}$$

The straight line segment (**dark blue line**) empirically plotted by the trial-error method represents the relationship of pT1 as a function of $\overline{\Delta X}$ between $Bz1_{min}$ and $|Bz1_{min}|$. It crosses the abscissa axis to the value $\overline{\Delta X}$ = -5.56 nm and has a slope according to (J48):

$$(K_{z1} \cdot \chi_{z1}) = 1/5.56 = 0.18 \text{ nm}^{-1} \quad \Rightarrow \quad K_{z1} = 0.18/0.133 = 1.35$$

The interpolation of $K_{z1}$ using the expression (J23) give:

$$q_{z1} = 1.99$$

From the previous data, the following calculations are made:

$$Bz1_{min} = -3 \text{ nm}$$

$$Bz1_{Max} = -4.8 \text{ nm}$$

$$q_{z2} = q_{z1} + \frac{Ln(0.062/0.133)}{Ln5500} = 1.9$$

$$K_{z2} = 1.7$$

$$Bz2_{min} = -6.8 \text{ nm}$$

$$Bz2_{Max} = -16.7 \text{ nm}$$

**Zone O (dark blue line) ≡ [** -3 nm ; + 3nm]

$$pT1 = 1 + 0.18 \cdot \overline{\Delta X}$$

Equation represented by a straight line segment extendable to $-\delta X_{z1}$, i. e. -4 nm

**Mixed Zone (purple line) ≡ [ -4.8 nm ; -3 nm [**

$$pT1 = \left(\frac{4.8 + \overline{\Delta X}}{1.8}\right) \cdot \left(1 + 0.18 \cdot \overline{\Delta X}\right) - \left(\frac{3 + \overline{\Delta X}}{1.8}\right) \cdot \left(0.715 + 0.105 \cdot \overline{\Delta X}\right)$$

Equation represented by a parabolic arch

**Zone 2 True (light blue line) ≡ [ -6.8 nm ; -4.8 nm [**

$$pT1 = 0.715 + 0.105 \cdot \overline{\Delta X}$$

Equation represented by a straight line segment

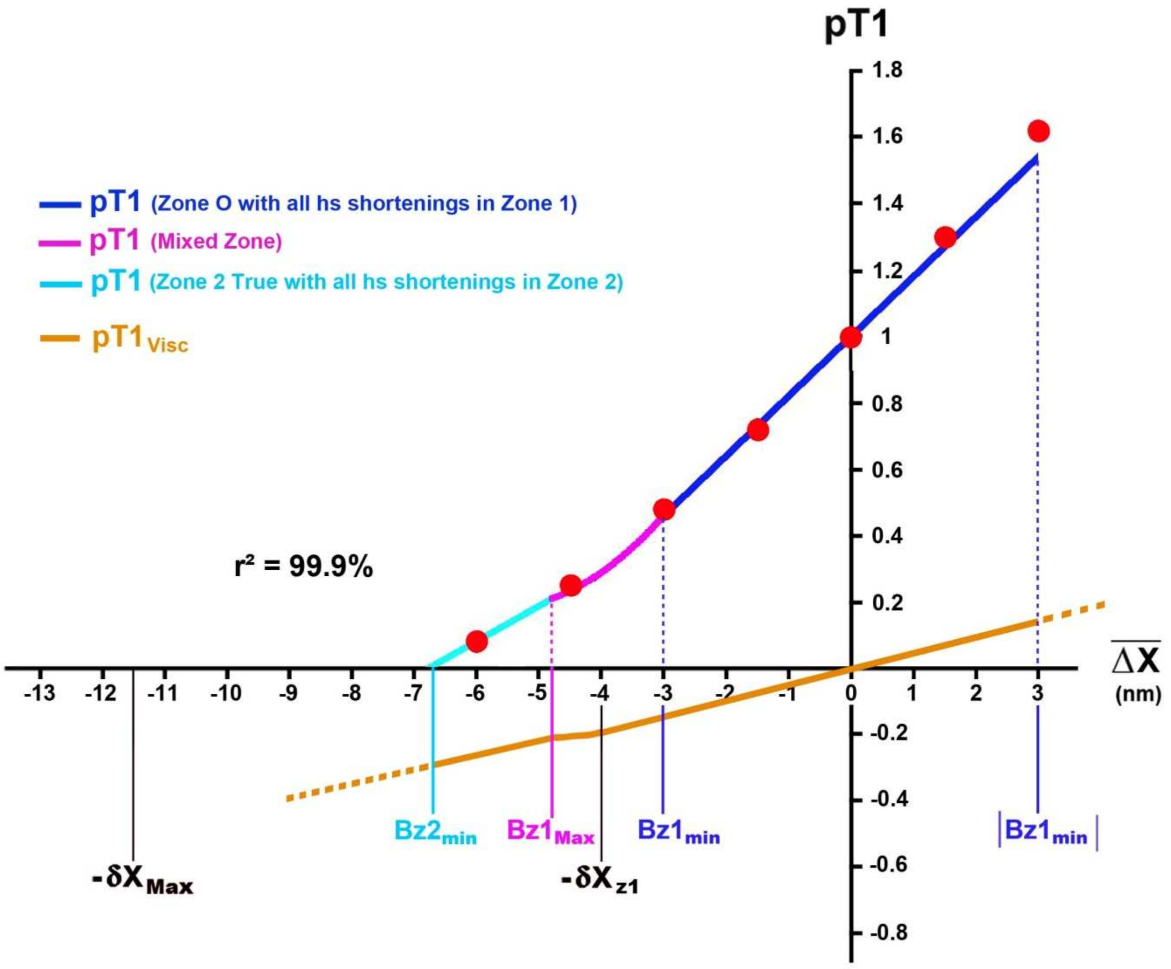


**Fig J8. Relation of pT1 as a function of the average shortening of a hs ($\overline{\Delta X}$) according to the equations of the model: straight line segment for Zone O between $Bz1_{min}$ and $|Bz1_{min}|$ (dark blue line), convex parabolic arc for Mixed Zone between $Bz1_{Max}$ and $Bz1_{min}$ (purple line), and straight line segment for Zone 2 True between $Bz2_{min}$ and $Bz1_{Max}$ (light blue line). The orange line represents the relationship of the force generated by the viscosity ($pT1_{Visc}$) as a function of $\overline{\Delta X}$. The red dots come from Fig 13 in [1].**

**Note**

The linear regression between theoretical and experimental pT1 values gives a determination coefficient of 99.9%.

#### J.7 Explicit relationship of the tension at the end of phase 1 after a length step in Zone 1 Enlarged

As noted in the example in paragraph J.6, the equation of the straight line segment given in (J22) and (J43) relating to the interval [$Bz1_{min}$ ; $|Bz1_{min}|$ ] characterizing Zone O (Fig J8) can be extended to the interval [$-\delta X_{z1}$ ; $|Bz1_{min}|$ ] defining the "Zone 1 Enlarged" (Fig J5a). An experiment, entitled "control" and marked "c", is carried out. The relationship between T1 and $\overline{\Delta X}$ in Zone 1 Enlarged is formulated:

$$T1_{z1} = T0_c \cdot \left(1 + K_{z1} \cdot \chi_{z1} \cdot \overline{\Delta X}\right) \cdot \mathit{1}_{\left[-\delta X_{z1};\left|Bz1_{min}\right|\right]}\left(\overline{\Delta X}\right) \qquad \text{(J50)}$$

where $T0_c$ is the tension during the isometric tetanus plateau for the control experiment.

Equation (J50) represents a straight line with a slope $e_0$ that intersects the abscissa axis in $-Y_0$, so:

$$e_0 = T0_c \cdot \chi_{z1} \cdot K_{z1} \qquad \text{(J51a)}$$

$$Y_0 = \frac{1}{\chi_{z1} \cdot K_{z1}} \qquad \text{(J51b)}$$

Parameters $e_0$ and $Y_0$ are stiffness and strain related to the control experiment.

#### J.8 Forces induced by the presence of viscosity

In Zone 1 Enlarged, the relative tension caused by the viscous original forces at the end of phase 1 ($pT1_{Visc,z1}$) is calculated by subtraction between (J50) and (J7) as:

$$pT1_{Visc,z1} = \left[\chi_{z1} \cdot \left(K_{z1} - 1\right) \cdot \overline{\Delta X}\right] \cdot \mathbf{1}_{\left[-\delta X_{z1};\left|Bz1_{min}\right|\right]}\left(\overline{\Delta X}\right) \qquad \text{(J52a)}$$

In Zone 2 True defined by the range [$Bz2_{min}$ ; $Bz1_{Max}$ ], the relative tension generated by the end of phase 1 ($pT1_{Viscs,z2}$) is determined by subtraction between (J33) and (J27) according to:

$$pT1_{Visc,z2} = \left[\chi_{z2} \cdot \left(K_{z2} - 1\right) \cdot \overline{\Delta X}\right] \cdot \mathbf{1}_{\left[Bz2_{min};Bz1_{Max}\right]}\left(\overline{\Delta X}\right) \qquad \text{(J52b)}$$

In the example of paragraph J.6, the expressions (J52a) and (J52b) are represented by two orange straight line segments (Fig J8). With the data, the slopes of the two segments are calculated:

$$[\chi_{z1} \cdot (K_{z1} - 1)] = 0.133\ nm^{-1} \cdot 0.35 = 0.047\ nm^{-1}$$

$$[\chi_{z2} \cdot (K_{z2} - 1)] = 0.062\ nm^{-1} \cdot 0.7 = 0.044\ nm^{-1}$$

The two slopes have almost identical values, equal to about 35% of $\chi_{z1}$ and 70% of $\chi_{z2}$.

The viscosity force at the end of phase 1 in the range [$Bz1_{Max}$ ; $-\delta X_{z1}$] is an orange parabolic arc (Fig J8) calculated on the model of (J48) and (J49).

#### J.9 Influence of the duration of phase 1 ($\tau_{p1}$)

A control experiment is performed on an isolated muscle fiber under specific conditions. Then a distinct experiment is carried out where all the characteristic parameters of the control experiment are identical except for the duration of phase 1 such that:

$$\boldsymbol{\tau_{p1,d} \neq \tau_{p1,c}}$$

where $\tau_{p1,c}$ and $\tau_{p1,d}$ are the durations of phase 1 for control and distinct experiments, labelled "c" and "d", respectively.

Under equal conditions (and in particular with regard to internal temperature), the proportionality coefficient ($\phi_{hs}$) common to the two solids {Z-disk$_s$+2Afil} and {M-disk$_s$+2Mfil} is a constant. It is checked according to (J5):

$$\phi_{hs} = \frac{\nu_c \cdot T0_c \cdot \tau_{p1,c}}{N_m} = \frac{\nu_d \cdot T0_c \cdot \tau_{p1,d}}{N_m}$$

where $\nu_c$ and $\nu_d$ are the 2 viscous parameters related to the control and distinct experiments, respectively.

As a result:

$$\nu_d = \nu_c \cdot \frac{\tau_{p1,c}}{\tau_{p1,d}} \qquad \text{(J53)}$$

The constitutive coefficient of the presence of viscosity in the distinct experiment in Zone 1 Enlarged ($q_{d,z1}$) is calculated on the model of (J24):

$$q_{d,z1} = \frac{Ln\chi_{z1} - Ln\nu_d}{LnN_{hs}}$$

where $\chi_{z1}$ is the elastic origin stiffness that in accordance with (J8) remains constant between the 2 experiments.

By entering (J53), we obtain:

$$q_{d,z1} = q_{c,z1} + \frac{1}{Ln N_{hs}} \cdot Ln\left(\frac{\tau_{p1,d}}{\tau_{p1,c}}\right) \qquad \text{(J54)}$$

where $q_{c,z1}$ is the characteristic parameter of the presence of viscosity in Zone 1 Enlarged for the control experiment calculated from (J24).

The tension at the end of phase 1 of a length step in the distinct experiment in Zone 1 Enlarged ($T1_{d,z1}$) is formulated on the model of (J50):

$$T1_{d,z1} = T0_c \cdot \left(1 + \chi_{z1} \cdot K_{d,z1} \cdot \overline{\Delta X}\right) \qquad \text{(J55)}$$

where $K_{d,z1}$ is the multiplier coefficient in the distinct experiment calculated according to (J23):

$$K_{d,z1} = N_{hs}^{\,1-q_{d,z1}/2} \cdot \coth\left(N_{hs}^{\,1-q_{d,z1}/2}\right) + N_{hs}^{\,1-q_{d,z1}} \qquad \text{(J56)}$$

***Example with points collected in Fig 19 in [1].***

The data of the **control experiment** are identical to those of Fig J8 and are presented in the first column of Table 1 as:

$$\boldsymbol{\tau_{p1,c} = 0.2\ ms}$$

$$N_{hs} = 5500$$

$$q_c = q_{z1} = 1.99$$

$$\chi_{z1} = 0.133\ nm^{-1}$$

$$\chi_{z2} = 0.062\ nm^{-1}$$

Based on equation (J48), the plot of $pT1_c$ as a function of $\overline{\Delta X}$ appears in Fig J9 as a **dark blue** line in Zone 1 Enlarged, Mixed Zone and Zone 2 True; it is the identical reproduction (except for the choice of colors) of the one in Fig J8. There is good match between the theoretical model and the experimental data (**blue dots**) with a determination coefficient of 99.7%.

The **distinct experiment** is performed under identical experimental conditions except for the duration of phase 1 of the length steps with $\boldsymbol{\tau_{p1,d} = 1\ ms}$**.**

The viscous parameter relating to Zone 1 Enlarged ($q_{d,z1}$) is calculated from (J54):

$$q_{d,z1} = \left[1.99 + \frac{1}{\text{Ln}5500} \cdot \text{Ln}\left(\frac{1\,\text{ms}}{0.2\,\text{ms}}\right)\right] = 2.18$$

In Zone 1 Enlarged, the multiplier coefficient ($K_{d,z1}$) is determined with (J56):

$$K_{d,z1} = 5500^{-0.09} \cdot \coth\left(5500^{-0.09}\right) + 5500^{-1.18} = 1.07$$

The product ($\chi_{z1}$·$K_{d,z1}$) is deduced:

$$(\chi_{z1} \cdot K_{d,z1}) = 0.133\ nm^{-1} \cdot 1.07 = 0.142\ nm^{-1}$$

The viscosity coefficient in Zone 2 True ($q_{d,z2}$) is calculated from (J36):

$$q_{d,z2} = q_{d,z1} + \frac{1}{\text{Ln}N_{hs}} \cdot \text{Ln}\frac{\chi_{z2}}{\chi_{z1}} = \left[2.18 + \frac{1}{\text{Ln}5500} \cdot \text{Ln}\left(\frac{0.062}{0.133}\right)\right] = 2.09$$

In Zone 2 True, the multiplier coefficient ($K_{d,z2}$) is calculated using (J35):

$$K_{d,z2} = 5500^{-0.045} \cdot \coth\left(5500^{-0.045}\right) + 5500^{-1.09} = 1.15$$

The product ($\chi_{z2}$·$K_{d,z2}$) is deduced:

$$(\chi_{z2} \cdot K_{d,z2}) = 0.062\ nm^{-1} \cdot 1.15 = 0.071\ nm^{-1}$$

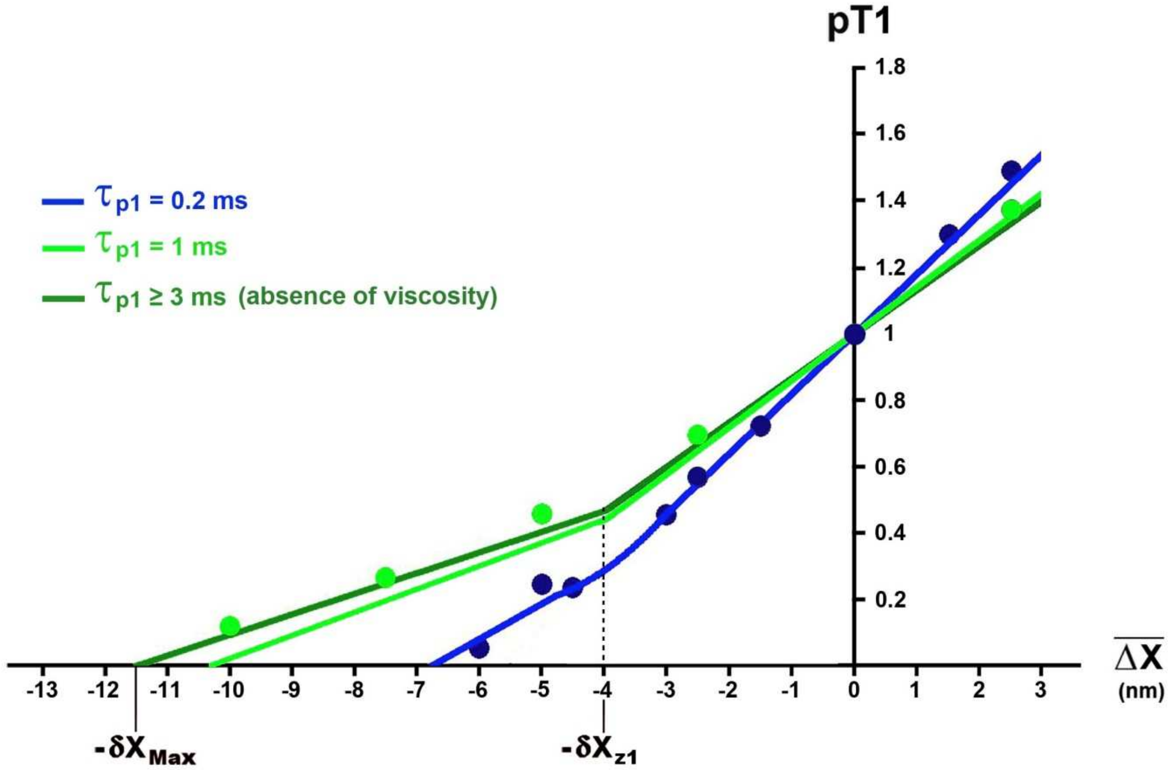


**Fig J9. Relations of the relative tension at the end of phase 1 (pT1) as a function of the average shortening of a hs ($\overline{\Delta X}$) according to duration of phase 1 ($\tau_{p1}$).**

The blue and light green dots that refer to a phase 1 duration of 0.2 ms and 1 ms, respectively, are taken from Fig 19 in [1].

The plot of $pT1_d$, the relative tension at the end of phase 1 with a duration **$\tau_{p1,d}$ = 1 ms**, consists of the two straight segments (**light green** line), the Mixed Zone being extremely small. We note that the values of $pT1_d$ are, on the one hand, significantly higher than those of $pT1_c$, the relative tension at the end of phase 1 for **$\tau_{p1,c}$ = 0.2 ms** and, on the other hand, close to those of $pT1_{Elas}$, the relative tension at the end of phase 1 in the absence of viscosity (**dark green** straight line), whose equations are provided in (J7) and (J27) relative to Zones 1 and 2.

The increase in the duration of phase 1 induces the increase of the parameters q according to (J54) and the decrease of the multiplying coefficients K with (J56). Logically, as the shortening speed decreases, the influence of viscosity decreases with it, and the tension values at the end of phase 1 approximate those determined in the absence of viscosity. Thus the slopes ($\chi_{z1}\cdot K_{d,z1}$) and ($\chi_{z2}\cdot K_{d,z2}$) equal to à 0.142 $nm^{-1}$ and 0.071 $nm^{-1}$ approach respectively the elastic origin slopes $\chi_{z1}$ and $\chi_{z2}$ equal to à 0.133 $nm^{-1}$ and 0.062 $nm^{-1}$.

Conversely, the decrease in the duration of phase 1 implies a greater presence of viscosity with an increase in the multiplying coefficients K.

In Fig J9, the **light green** dots corresponding to **$\tau_{p1,d}$ = 1 ms**, are slightly above the **light green** line. Indeed, since the duration of phase 1 becomes longer than the average time rapid initiation of a WS ($\tau_{startF}$ = 0.7 ms), the rapid tension rise of phase 2 has begun.

At the end of sub-paragraph J.3.8, the value "q=2.3" is proposed to notify the absence of viscosity (Fig J3). The duration of phase 1 corresponding to $q_d$=2.3 is noted "$\tau_{p1,Max}$"; its value is calculated according to (J54) :

$$2.3 = 1.99 + \frac{1}{\mathrm{Ln}5500} \cdot \frac{\tau_{p1,Max}}{0.2\,\mathrm{ms}}$$

So:

$$\tau_{p1,Max} = 0.2\,\mathrm{ms} \cdot e^{[\mathrm{Ln}5500 \cdot (2.3-1.99)]} \approx 3\mathrm{ms}$$

We note that 3 ms is equivalent to the duration of phase 2 of a length step, hence the hiatus:

If phase 1 is short ($\tau_{p1} \leq$ 0.2 ms), the initiation of WS heads is not feasible: phase 2 starts at the end of phase 1 where viscosity is strongly present.

If phase 1 is not brief ($\tau_{p1}$ > 0.2 ms), the initiation of WS heads becomes possible: phases 1 and 2 coexist during $\tau_{p1}$; viscosity becomes less significant during phase 1 but remains effective as long as $\tau_{p1}$ is less than 3 ms.

#### J.10 Influence of tetanus plateau tension under isometric conditions (T0)

An experiment control is carried out where $T0_c$ is the tension of the isometric tetanus plateau. Following this, a "new" experiment labelled "n" is performed where all the characteristic parameters of the control experiment are identical, except for the tension of the isometric tetanus plateau, such that:

$$T0_n \neq T0_c$$

where $T0_n$ is the tension of the isometric tetanus plateau related to the new experiment.

This condition can be effectuated in several ways: lengthening the initial length of the sarcomere, changes in calcium or inorganic phosphate concentration, shortening of the fiber at constant velocity, presence or not of an inhibitor of cross-bridges, etc.

By following a similar path to that of the previous paragraph, i.e. by using the constancy of the parameter $\phi_{hs}$ defined in (J4) and (J5), the constituent coefficient of the presence of the viscosity relative to the new experiment ($q_{n,z1}$) is calculated in Zone 1 Enlarged according to:

$$q_{n,z1} = q_{c,z1} + \frac{1}{LnN_{hs}} \cdot Ln \frac{T0_n}{T0_c} \tag{J57}$$

In Zone 1 Enlarged, the tension at the end of phase 1 of a length step in the new experiment ($T1_{n,z1}$) is formulated with (J50):

$$T1_{n,z1} = T0_n \cdot \left(1 + \chi_{z1} \cdot K_{n,z1} \cdot \overline{\Delta X}\right) \tag{J58}$$

where $K_{n,z1}$ is the multiplier in the new experimentation determined according to (J23):

$$K_{n,z1} = N_{hs}{}^{1-q_{n,z1}/2} \cdot \coth\left(N_{hs}{}^{1-q_{n,z1}/2}\right) + N_{hs}{}^{1-q_{n,z1}} \tag{J59}$$

In Zone 2 True, the following relationship is obtained in support of (J36):

$$q_{n,z2} = q_{n,z1} + \frac{1}{LnN_{hs}} \cdot Ln\left(\frac{\chi_{z2}}{\chi_{z1}}\right) = q_{c,z1} + \frac{1}{LnN_{hs}} \cdot Ln\left(\frac{\chi_{z2} \cdot T0_n}{\chi_{z1} \cdot T0_c}\right) \tag{J60}$$

where $q_{n,z2}$ is the characteristic coefficient of the presence of viscosity for new experiment in Zone 2 True.

From (J33) et with (J60), we deduce $K_{n,z2}$, the multiplier in Zone 2 True.

***Example with points collected in Figs 6 and 11 in [3]***

According to the expression (8) of Paper 4, the tension of the isometric tetanus plateau is a linear function of the number of myosin heads in WS, which itself is a decreasing affine function of the initial length of the sarcomere ($L0_s$) between 2.25 and 3.65 µm. This assertion was discovered experimentally [4] and is theoretically demonstrated in accompanying Paper 3. From Fig 12 in [4], it can be deduced:

$$pT0_n\left(L0_s\right) = \frac{T0_n\left(L0_s\right)}{T0_c} = \mathbf{1}_{[2\mu m;2.25\mu m[}\left(L0_s\right) + \left(2.607 - 0.7143 \cdot L0_s\right) \cdot \mathbf{1}_{[2.25\mu m;3.65\mu m[}\left(L0_s\right) \qquad \text{(J61)}$$

where $T0_n$ is the tension of the isometric tetanus plateau related to the new experiment with $L0_s$ between 2.25 and 3.65 µm; $T0_c$ is the maximum tension for an isometric tetanus plateau when $L0_s$ is between 2 and 2.25 µm related to the control experiment.

**<u>Methodology</u>**

1/ Using (J61), $L0_s$ provides the corresponding value of $pT0_n$ as:

**$L0_s$ = 2.2 µm ⇒ $pT0_c$ = 1**

**$L0_s$ = 2.6 µm ⇒ $pT0_{2.6}$ = 0.75**

**$L0_s$ = 3.1 µm ⇒ $pT0_{3.1}$ = 0.39**

2/ For the control experiment, the value " $q_{c,z1}$ = 1.98" is used:

**$L0_s$ = 2.2 µm ⇒ $q_{2.2,z1} = q_{c,z1}$ = 1.98 ⇒ $K_{2.2,z1} = 5500^{0.01} \cdot \coth(5500^{0.01})$ = 1.37**

The values of $q_{n,z1}$ for **$L0_s$ = 2.6 µm** and **$L0_s$ = 3.1 µm** are determined in Zone 1 Enlarged according to (J57) then the associated parameter K is determined with (J59).

**$L0_s$ = 2.6 µm ⇒ $q_{2.6,z1}$ = 1.98 + 0.116·Ln(0.75) = 1.94 ⇒ $K_{2.6,z1} = 5500^{0.03} \cdot \coth(5500^{0.03})$ = 1.49**

**$L0_s$ = 3.1 µm ⇒ $q_{3.1,z1}$ =1.98 + 0.116·Ln(0.39) = 1.87 ⇒ $K_{3.1,z1} = 5500^{0.065} \cdot \coth(5500^{0.065})$ = 1.86**

3/ Since the slope $\chi_{z1}$ is a constant equal to 0.133 $nm^{-1}$ (first column of Table 1 to Paper 4), the calculations of pT1 in Zone 1 Enlarged are made with (J58) for each of the 3 initial lengths of the sarcomere.

Same goes for Mixed Zone and Zone 2 True with (J60).

The theoretical model is checked with the 2 examples from Figs 6 and 11 in [3] where $q_{c,z1}$ is equal to 1.98 and 1.99, respectively. The relationships of the relative tension at the end of phase 1 (pT1) for each of the $L0_s$ values are plotted in Figs J10a and J10b. A correct match between experimental and calculated values is noted with $r^2$ > 98%.

Thus, the only knowledge of the 2 parameters $q_{c,z1}$ and $N_{hs}$ allows to calculate the relation of pT1 according to $\overline{\Delta X}$ in the 3 zones, Zone 1 Enlarged, Mixed Zone, Zone 2 True, for any initial length of the sarcomere between 2 and 3.65 µm.

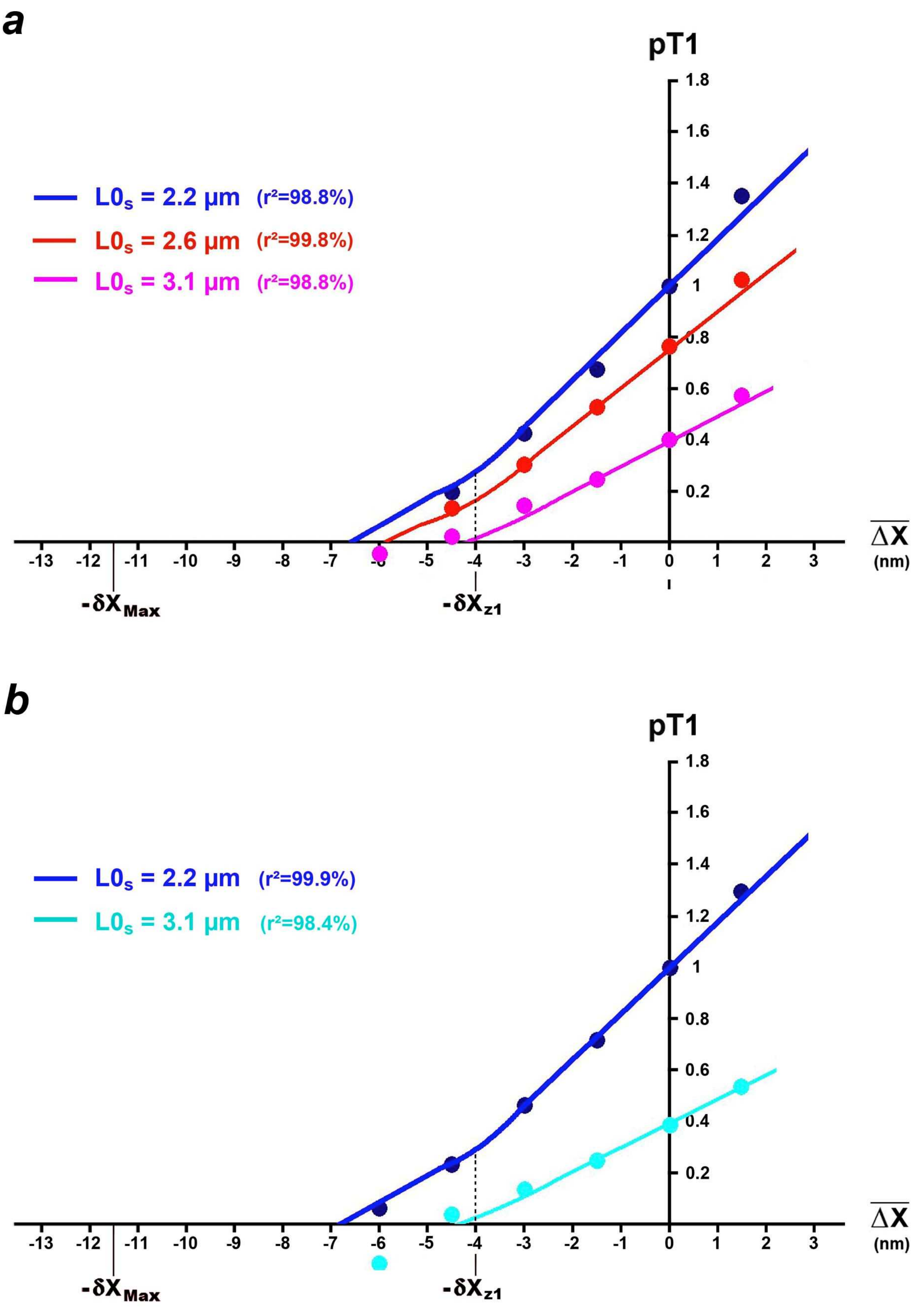


**Fig J10. Relations of pT1 with $\overline{\Delta X}$ according to the initial length of the sarcomere ($L0_s$).**

(a) and (b) Plots based on Figs 6 and 11 in [3], respectively.

**Note**

According to the definition of stiffness given in (J53a), the stiffnesses relating to both lengths 2.2 and 2.6 µm are formulated with (J58) in Zone 1 Enlarged:

$$e_{2.2,z1} = T0_{2.2} \cdot \chi_{z1} \cdot K_{2.2,z1}$$

$$e_{2.6,z1} = T0_{2.6} \cdot \chi_{z1} \cdot K_{2.6,z1}$$

where $e_{2.2,z1}$ is the stiffness relative to the control experiment which in the example corresponds to a initial length of sarcomere between 2 and 2.25 µm, hence the notation "2.2"; $e_{2.6,z1}$ is the stiffness relative to the new experiment for a initial length of sarcomere equal to 2.6 µm, hence the acronym "2.6".

Their ratio is calculated with the previous data:

$$e_{2.6,z1} / e_{2.2,z1} = 0.75 \cdot (1.49/1.37) = 0.81$$

In Zone 2 True, the calculations provide:

$$e_{2.6,z2} / e_{2.2,z2} = 0.75 \cdot (1.95/1.74) = 0.84$$

The ratio of the two stiffnesses relative to these two initial lengths of sarcomere is a quasi-constant independent of the size of the step in length or force. This assertion is confirmed with Fig 3 in [5].

#### J.11 Influence of instantaneous or intermediate tetanus tension ($T0_i$)

##### *J.11.1 Introduction*

Instead of studying the transformations produced by a limited number of "new" experiments, we generalize to a temporal succession of new experiments during which the isometric tetanus tension varies ($T0_i$). Such temporal changes for $T0_i$ can be obtained:

- during the development or redevelopment of the tension of the fiber under isometric conditions
- by increasing the calcium concentration (pCa) with values above 4.5, the normal rate.
- in the presence of a strong binding or working stroke inhibitor: the muscle fiber is tetanized under normal conditions and then at selected times the fiber is infused at different concentrations of an inhibitor product such as *N-benzyl_p-toluene sulphonamide* (BTS).
- with a series of force steps where the shortening velocity of the fiber is steady.
- in the presence of components of the ATP hydrolysis reaction (Pi, ADP or ATP) measured at different concentrations..

Such experimental measurements are often made by subjecting the fiber to forced oscillations, with a frequency equal to 4 kHz and a peak-to-peak amplitude of 2 nm, i.e. a succession of shortenings and elongations of 2 nm (Fig J7) belonging to the Zone O (O for oscillations) defined in paragraph J.5.

##### *J. 11.2 General equations*

The reference isometric tetanus tension ($T0_c$) is the isometric tension of the tetanus plateau of the "control" experiment under specified conditions. When the fiber is tetanized outside these conditions, for example when rising to the plateau, the instantaneous tetanus tension ($T0_i$) is acquired at different times t and each measurement corresponds to an "instantaneous" experiment.

The instantaneous relative tension ($pT0_i$) is equal to:

$$pT0_i(t) = \frac{T0_i(t)}{T0_c} \qquad \text{(J62)}$$

Since most of the length steps in "instantaneous" experimentations belong to Enlarged Zone 1, all calculations will refer to this zone. The instantaneous coefficient ($q_i$) constituting the presence of viscosity in the instantaneous experiment is calculated with (J57) and (J62) as:

$$q_i(t) = q_c + \frac{1}{LnN_{hs}} \cdot Ln\left[pT0_i(t)\right] \qquad \text{(J63)}$$

where $q_c = q_{c,z1}$

The instantaneous tension ($T1_i$) measured at the end of phase 1 of a length step over time ($t+\tau_{p1}$) in the instantaneous experiment is formulated according to (J58):

$$T1_i(t+\tau_{p1}) = T0_i(t)\cdot\left(1+\chi_i(t)\cdot K_i(t)\cdot\overline{\Delta X}\right) \quad \text{(J64)}$$

where $\chi_i$ is the instant coefficient of elastic origin stiffness in Zone 1 estimated from (J8):

$$\chi_i(t) = \frac{1}{\delta X_{Max} - \delta X_{T,i}/2} \quad \text{(J65)}$$

with $\delta X_{T,i}$ as the instant linear range corresponding to the instant angular range $\delta\theta_{T,i}$ on which the orientation of the levers belonging to the WS heads is uniformly distributed at time t.

and where $K_i$ is the instant multiplier coefficient which is calculated according to (J23):

$$K_i(t) = N_{hs}^{1-q_i/2}\cdot\coth\left(N_{hs}^{1-q_i/2}\right)+N_{hs}^{1-q_i} \quad \text{(J66)}$$

Instant tension measurements are performed with short phase 1 durations ($\tau_{p1}$<120 µs) and most often by means of forced oscillations at 4 kHz ($\tau_{p1,4kHz}$ = 90 µs). We proceed with the following approximation "($t+\tau_{p1}$) ≈ t" and the equation (J64) is simplified:

$$T1_i(t) \approx T0_i(t)\cdot\left(1+\chi_i(t)\cdot K_i(t)\cdot\overline{\Delta X}\right) \quad \text{(J67)}$$

##### *J.11.3 Cases where control experimentation and instantaneous experiments have different durations of phase 1*

Instantaneous measurements of $T0_i$ and $T1_i$ are often performed using sinusoidal oscillations. It is demonstrated in paragraph J.5 that such oscillations are approximated by a series of successive length steps (Fig J7). The temporal frequency of the oscillations is expressed in kHz with a corresponding phase 1 duration ($\tau_{p1,osc}$). If the calculation of $q_c$ is not performed with oscillations or if it is performed at a different frequency, it is necessary to make a correction by coupling (J63) with (J54):

$$q_i(t) = \left[q_c + \frac{1}{LnN_{hs}}\cdot Ln\left(\frac{\tau_{p1,osc}}{\tau_{p1,c}}\right)\right] + \frac{1}{LnN_{hs}}\cdot Ln\left(\frac{T0_i(t)}{T0_c}\right) \quad \text{(J68)}$$

The most common frequency is 4 kHz with a corresponding phase 1 duration ($\tau_{p1.4kHz}$) equal to 90 µs according to the equality (J46).

***J.11.4 Cases where control and instantaneous experimentation have different isometric tetanus plateau tensions***

Changes in an experimental factor may affect the value of the tetanus plateau tension, such as intracellular calcium concentration (pCa), presence of inorganic phosphate ([Pi]), pH change or presence of a cross-bridge inhibitor. For example, the rise to an isometric tetanus plateau can be studied under conventional conditions where everything is normal (pCa=4.5, [Pi]=0, pH=7, no inhibitor, etc.), then one or more factors are modified: pCa>4.5, [Pi]>0, pH <7, etc.

The instantaneous viscous coefficient ($q_{i,c}$) under "conventional" conditions is formulated according to (J57):

$$q_{i,c}(t) = q_c + \frac{1}{LnN_{hs}} \cdot Ln\left(\frac{T0_{i,c}(t)}{T0_c}\right) \quad \text{(J69)}$$

where $T0_{i,c}$ and $T0_c$ are, respectively, the instantaneous tetanus tension and the isometric tetanus plateau tension under conventional conditions.

The instantaneous viscous coefficient ($q_{i,n}$) under "new" conditions is written according to (J63):

$$q_{i,n}(t) = q_n + \frac{1}{LnN_{hs}} \cdot Ln\left(\frac{T0_{i,n}(t)}{T0_n}\right) \quad \text{(J70a)}$$

where $T0_{i,n}$ and $T0_n$ are, respectively, the instantaneous tetanus tension and the isometric tetanus plateau tension under new conditions, i.e. after modification of a factor affecting the isometric tetanus plateau tension.

With (J57) and (J69), the equation (J70a) is rearranged:

$$q_{i,n}(t) = q_c + \frac{1}{LnN_{hs}} \cdot Ln\left(\frac{T0_{i,n}(t)}{T0_c}\right) \quad \text{(J70b)}$$

When the instantaneous tensions under new conditions are reported to $T0_c$, equation (J70b) is equivalent to (J63). Consequently, the experimental points under conventional and new conditions must be on the same curve. This forecast will be verified with several examples.

***J. 11.5 Definitions of 6 parameters from the expression (J67)***

The instant stiffness ($e_i$) is written as:

$$e_i(t) = \chi_i(t) \cdot K_i(t) \cdot T0_i(t) \quad \text{(J71a)}$$

The normalized instant stiffness ($e_i / e_0$) is formulated:

$$\frac{e_i(t)}{e_0} = \frac{\chi_i(t) \cdot K_i(t)}{\chi_0 \cdot K_0} \cdot pT0_i(t) \quad \text{(J71b)}$$

where $e_0$ is given in (J51a); $\chi_0$ and $K_0$ are identical to the two parameters of the control experiment, $\chi_{z1}$ and $K_{z1}$, respectively.

The instant stiffness in relation to the reference tension of the isometric tetanus plateau ($e_i / T0_c$) is:

$$\frac{e_i(t)}{T0_c} = \chi_i(t) \cdot K_i(t) \cdot pT0_i(t) \quad \text{(J71c)}$$

The instant compliance ($C_i$), i.e. the inverse of instant stiffness, is written as:

$$C_i(t) = \frac{1}{\chi_i(t) \cdot K_i(t) \cdot T0_i(t)} \quad \text{(J71d)}$$

The instant strain ($Y_i$) is formulated:

$$Y_i(t) = \frac{1}{\chi_i(t) \cdot K_i(t)} \quad \text{(J71e)}$$

The normalized instant strain ($Y_i / Y_0$) is written as:

$$\frac{Y_i(t)}{Y_0} = \frac{\chi_0 \cdot K_0}{\chi_i(t) \cdot K_i(t)} \quad \text{(J71f)}$$

where $Y_0$ is determined in (J51b).

From the previous equations, the following equations are derived:

$$e_i(t) \cdot Y_i(t) = T0_i(t) \quad \text{(J72a)}$$

$$e_0 \cdot Y_0 = T0_c \quad \text{(J72b)}$$

$$\chi_0 \cdot Y_0 \cdot K_0 = 1 \quad \text{(J72c)}$$

With this reminder, the 6 parameters defined from (J71a) to (J71f) are interdependent and can be deduced from each other.

We also note that the 5 parameters, $e_i$, $Y_i$, $e_i/e_0$, $Y_i / Y_0$ and $Y_i / T0_c$, tend towards zero when $pT0_i$ tends towards zero, i.e. when the fiber is close to rest or super-relaxing:

- $e_i$, $e_i/e_0$, and $e_i / T0_c$ because the influence of the term $T0_i$ overrides that of others

- $Y_i$ and $Y_i / Y_0$ because the term $1/K_i$ tends towards zero

##### *J.11.6 Instant slope of elastic origin*

With (J71b) and (J71e), the instant elastic slope ($\chi_i$) is calculated relative to $e_i/e_0$ and $Y_i$ as:

$$\chi_i(t) = \frac{e_i(t)}{e_0} \cdot \left( \frac{\chi_0 \cdot K_0}{K_i(t) \cdot pT0_i(t)} \right) \qquad \text{(J73a)}$$

$$\chi_i(t) = \frac{1}{Y_i(t)} \cdot \left( \frac{1}{K_i(t)} \right) \qquad \text{(J73b)}$$

where $K_i$ is a theoretical multiplier coefficient calculated from (J66).

The other 4 parameters are associated with $\chi_i$ in a similar way.

##### *J.11.7 Instant linear and angular ranges used as "analytical nanoscope"*

In our model, the stroke size ($\delta X_{Max}$) is a constant, so the slope $\chi_i$ depends only on the instant linear range $\delta X_{T,i}$ according to (J65). Relative to $e_i/e_0$ and $Y_i$, expressions (J73a) and (J73b) provide the following two equations:

$$\delta X_{T,i}(t) = 2 \cdot \left( \delta X_{Max} - \frac{e_0}{e_i(t)} \cdot \frac{K_i(t) \cdot pT0_i(t)}{\chi_0 \cdot K_0} \right) \qquad \text{(J74a)}$$

$$\delta X_{T,i}(t) = 2 \cdot \left( \delta X_{Max} - Y_i(t) \cdot K_i(t) \right) \qquad \text{(J74b)}$$

The instant linear and angular ranges $\delta X_{T,i}$ and $\delta\theta_{T,i}$ are linked by the expression (9) in Paper 4:

$$\delta X_{T,i}(t) = L_{S1b} \cdot R_{WS} \cdot \delta\theta_{T,i}(t) \qquad \text{(J75)}$$

The instantaneous angular range ($\delta\theta_{T,i}$) relative to $e_i/e_0$ on the one hand, and to $Y_i$ on the other hand, is equal by affine transformation to:

$$\delta\theta_{T,i}(t) = 2 \cdot \left( \delta\theta_{Max} - \frac{e_0}{e_i(t)} \cdot \frac{K_i(t) \cdot pT0_i(t)}{\chi_0 \cdot K_0 \cdot L_{S1b} \cdot R_{WS}} \right) \qquad \text{(J76a)}$$

$$\delta\theta_{T,i}(t) = 2 \cdot \left( \delta\theta_{Max} - Y_i(t) \cdot \frac{K_i(t)}{L_{S1b} \cdot R_{WS}} \right) \qquad \text{(J76b)}$$

It is easy with the formulas (J71a), (J71c), (J71d) and (J71f) to obtain the expressions of $\delta X_{T,i}$ and $\delta\theta_{T,i}$ for the 4 other parameters: $e_i$ , $e_i/T0_c$ , $C_i$ and $Y_i/Y_0$ .

**Note:** Equations (J62) to (J76b) also apply to time-independent experimental series, such as force step series or length step series at different intracellular concentrations of calcium, inorganic phosphate or a cross-bridge inhibitor. In this case, $T0_i$ is referred to as "i**ntermediate** tetanus tension".

We call back,

In a half-sarcomere on the right: $\delta\theta_{T,i}(t) \equiv \lfloor \theta_{T,i}(t); \theta_{up} \rfloor$ with $\theta_{down} \le \theta_{T,i}(t) \le \theta_{up}$

In a half-sarcomere on the left: $\delta\theta_{T,i}(t) \equiv \lfloor \theta_{up}; \theta_{T,i}(t) \rfloor$ with $\theta_{up} \le \theta_{T,i}(t) \le \theta_{down}$

Common to all hs, the instantaneous angular range $\delta\theta_{T,i}$ characterizes the instantaneous domain within which the θ orientation of the levers belonging to the WS myosin heads has as a first approximation a uniform density and outside which no lever exerts any action.

We are interested in the relationship of $\delta\theta_{T,i}$ as a function of the instantaneous relative tension ($pT0_i$) which varies between 0 and 1, i.e. between two equilibrium states :

**Equilibrium 1 with $pT0_i = 0$ and $\delta\theta_{T,i} = 0$** : these values characterize the **total relaxation** (or rest) where no cross-bridge is present

**Equilibrium 2 with $pT0_i = 1$ and $\delta\theta_{T,i} = \delta\theta_T$** : these values determine the conditions of the **isometric tetanus plateau**

We postulate that the passage between these two states of equilibrium is carried out continuously or discretly in 4 different ways:

1/ Exponential rise from 0 to $\delta\theta_T$ with the possibility of exceeding above $\delta\theta_T$:

$$\delta\theta_{T,i} = \delta\theta_T \cdot \left[1 - e^{-\lambda_1 \cdot pT0_i}\right] \cdot \left(1 + \omega_1 \cdot pT0_i\right) \tag{J77}$$

where $\lambda_1$ and $\omega_1$ are 2 constants specific to fiber and experimentation.

The relationship (J77) is representative of a variation in intracellular concentration, the presence of a cross-bridge disruptor or inhibitor: pCa, [Pi], [BTS], etc.

2/ Exponential descent from $\delta\theta_{Max}$ to $\delta\theta_T$ with the possibility of overtaking below $\delta\theta_T$:

$$\delta\theta_{T,i} = \delta\theta_T + \left(\delta\theta_{Max} - \delta\theta_T\right) \cdot \left[e^{-\lambda_2 \cdot pT0_i}\right] \cdot \left(1 - \omega_2 \cdot pT0_i\right) \tag{J78a}$$

where $\lambda_2$ and $\omega_2$ are 2 positive constants specific to fiber and experimentation.

The solution (J78a) is characteristic of the rise towards the isometric tetanus plateau with shortening of the fiber before the real isometry. At the beginning of stimulation (t=0 and $pT0_i$ =0), the range $\delta\theta_{T,i}$ instantly increases from 0 to $\delta\theta_{Max}$, the range singularizing the initiation of a WS.

Variant with a pseudo-periodic initial phase:

$$\delta\theta_{T,i} = \delta\theta_T + \left(\delta\theta_{Max} - \delta\theta_T\right) \cdot \left[e^{-\lambda_2 \cdot pT0_i}\right] \cdot \cos\left(\omega_2 \cdot pT0_i\right) \tag{J78b}$$

3/ Sigmoidal descent from $\delta\theta_T$ to 0:

$$\delta\theta_{T,i} = \frac{\delta\theta_T}{1+e^{-\lambda_3 \cdot (pT0_i - \omega_3)}} \quad \text{(J79)}$$

where $\lambda_3$ and $\omega_3$ are 2 positive constants specific to fiber and experimentation.

The answer (J79) is indicated to describe the relaxation phase following the isometric tetanus plateau after the stimulation has stopped.

4/ Sigmoidal rise from $\delta\theta_T$ to $\delta\theta_{Max}$

$$\delta\theta_{T,i} = \delta\theta_{Max} + \frac{\delta\theta_T - \delta\theta_{Max}}{1+e^{-\lambda_4 \cdot (pT0_i - \omega_4)}} \quad \text{(J80)}$$

where $\lambda_4$ and $\omega_4$ are 2 positive constants specific to fiber and experimentation.

The expression (J80) is suitable for a series of force steps used to establish the Force/Velocity relationship.

With equations (J77) to (J80), the typical hs of the fiber is interpreted as a servo-driven motor system that responds to a disturbance between the two states of equilibrium, total relaxation and the isometric tetanus plateau. The adequate response consists of a non-stationary step with critical regime followed by a stationary step with stable state representative of one of the two equilibrium states which acts as a setpoint to be respected.

With (J75), we obtain similar formulations for $\delta X_{T,i}$, the corresponding instantaneous linear range. Through $\delta\theta_{T,i}$ or $\delta X_{T,i}$, an analytical "nanoscope" is acquired to study the evolution of the uniform density of the $\theta$ angle of the levers belonging to the WS heads between the two equilibrium states.

#### J.12 Study of instant strain ($Y_i$) and instant angular range ($\delta\theta_{T,i}$) with an example

1/ After being isolated from a rabbit psoas muscle, a fiber prepared under standard conditions undergoes a series of length steps that serve as a control experiment. The data are displayed in the dark red column of Table 1 to Paper 4 with in particular according to (J23):

$$N_{hs} = 3300 \quad \text{and} \quad q_{c,z1} = 2 \quad \Rightarrow \quad K_{c,z1} = 1.31$$

Based on equation (J48), the $pT1_c$ plot appears as a **dark blue** line on Fig J11a.

2/ A "new" experiment is performed on the same fiber where the conditions are unchanged except for an increase in the concentration of inorganic phosphate (Pi): the tension of the associated isometric tetanus plateau ($T0_n$) decreases, so according to (J57) and (J59):

**Add 10 mM Pi** $\Rightarrow$ $pT0_n = 0.573$ $\Rightarrow$ $q_{n,z1} = 2 + Ln(0.573) / Ln(3300) = 1.935$ $\Rightarrow$ $K_{n,z1} = 1.51$

In Zone 1 Enlarged, the relative tension at the end of phase 1 ($pT1_n = T1_n/T0_c$) is calculated from (J58). The equations for the Mixed Zone and Zone 2 True are determined using (J60). The $pT1_n$ plot is represented by a **red** line in Fig J11a. The **blue** and **red** dots are from Fig 1C in [6]. A correct agreement is found between theoretical and experimental values ($r^2 > 97.5\%$).

3/ We consider an index (i) relating to the Pi concentration associated to the relative intermediate tetanus tension ($pT0_i$). The values of $q_i$ and $K_i$ for $pT0_i$ are calculated with (J63) and (J66). The parameter $Y_i$ is calculated according to (J71e) where the instant elastic slope ($\chi_i$) is a constant equal to $\chi_{z1}$, i.e. 0.113 $nm^{-1}$. The $Y_i$ plot appears as a **green** line in Fig J11b.
In the same figure, there are 6 round coloured points corresponding to 6 concentrations of Pi equal to to 0, **3**, **10**, and **25** mM ; these points come from Fig 1D in [6]. With respect to the two straight line segments in Zone 1 Enlarged in Fig J11a, $Y_i$ calculation provides from (J71e):

$$[Pi_c] = 0 \text{ mM} \quad \Rightarrow \quad pT0_c = 1 \quad \Rightarrow \quad Y_0 = 6.76 \text{ nm}$$

$$[Pi_1] = 10 \text{ mM} \quad \Rightarrow \quad pT0_1 = 0.57 \quad \Rightarrow \quad Y_1 = 5.85 \text{ nm}$$

These 2 values are represented by 2 crosses, **blue** and **red**, in Fig J11b.
We obtain a fair agreement between theoretical and experimental $Y_i$ with $r^2 \approx 93\%$.

4/ The angular range $\delta\theta_{T,i}$ is empirically modelled according to (J77):

$$\delta\theta_{T,i} \quad \delta\theta_{T,i} = \delta\theta_T \cdot \left[1 - e^{-20 \cdot pT0_i}\right] \qquad \text{(J81)}$$

Equation (J81) is represented by a **light blue** line in the inset of Fig J11b. The 6 round points and 2 crosses are determined according to (J76b) from the experimental values. The assumption of constancy of $\delta\theta_{T,i}$ equal to $\delta\theta_T$ is verified if $pT0_i > 15\%$, i.e. for Pi concentrations below 25 mM. Consequently with (J65), (J75) and (J76b), equality " $\chi_i = \chi_{z1} = 0.113$ $nm^{-1}$ " is justified.

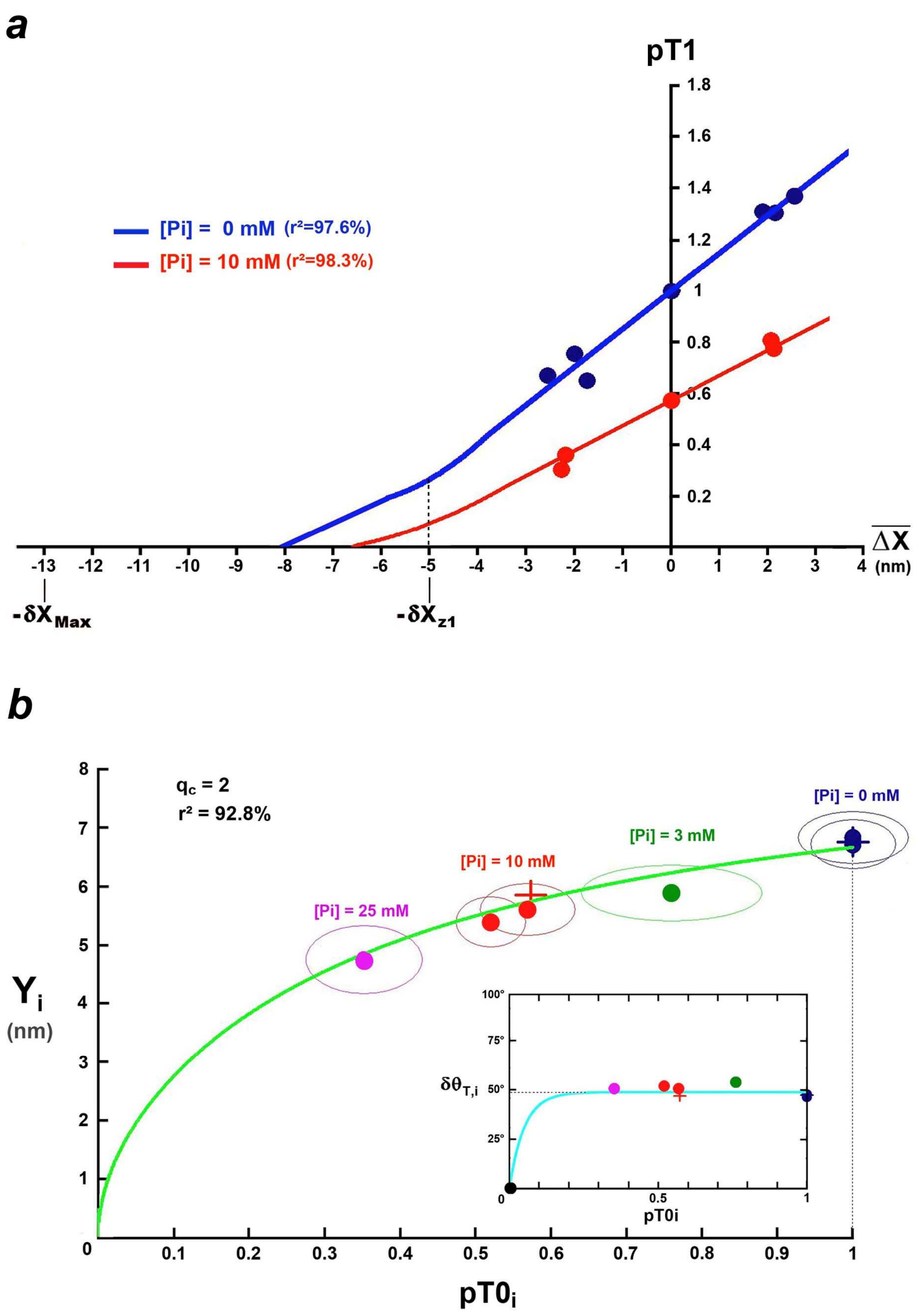


**Fig J11. (a) Relations of pT1 as a function of $\overline{\Delta X}$ according to the Pi concentration. The points are from Fig 1C in [6]. (b) Relation of $Y_i$ as a function of $pT0_i$ according to equation (J71e) of the model (green line). The 6 round coloured points come from Fig 1D in [6] and the 2 crosses result from the 2 values of $Y_i$ calculated from the 2 blue and red line segments in Zone 1 Enlarged of Fig J11a. In the inset, relationship of $\delta\theta_{T,i}$ (light blue line) as a function of $pT0_i$ according to (J81); the 6 round points and the 2 crosses are determined according to (J76b).**

#### J.13 Study of the instant linear range ($\delta X_{T,i}$) and instant normalized stiffness ($e_i/e_0$) during tension rise to the tetanus plateau according to three models

A fiber is extracted from the *lumbricalis″ digiti IV* muscle of *Rana Temporaria* and is subjected to a series of sinusoidal oscillations with an amplitude of 2 nm·hs$^{-1}$ peak to peak with a frequency of 4 kHz during the rise to the tetanus isometric plateau.

We pose: $\delta X_{Max}$ = 12 nm, $\delta X_T$ = 8 nm, $N_{hs}$ =2000 and $q_c$ = 1.88.

The value of $q_i$ is evaluated using (J63):

$$qi = 1.88 + 0.132 \cdot Ln\,(pT0_i)$$

The corresponding theoretical factor $K_i$ is determined according to (J66).

The linear range $\delta X_{T,i}$ is empirically modelled in three ways, twice according to (J78a) and once according to (J78b). After affine transformations with (J75):

$$\cdot\,\delta X_{T,i} = \delta X_T + (\delta X_{Max} - \delta X_T)\left[e^{-20 \cdot pT0_i}\right] \qquad \text{(J82a)}$$

$$\delta X_{T,i} = \delta X_T + (\delta X_{Max} - \delta X_T)\cdot\left[e^{-7.5 \cdot pT0_i}\right]\cdot(1 - 7 \cdot pT0_i) \qquad \text{(J82b)}$$

$$\delta X_{T,i} = \delta X_T + (\delta X_{Max} - \delta X_T)\cdot\left[e^{-9.5 \cdot pT0_i}\right]\cdot\cos(13.4 \cdot pT0_i) \qquad \text{(J82c)}$$

Equations (J82a), (J82b) and (J82c) are represented by a **light blue** line in Figs J12a, J12b and J12c, respectively. A **light blue** vertical line appears at $pT0_i$=0 to indicate that when the stimulation starts ($\delta X_{T,i}(t=0)$= 0), the WS initiation is instantaneous with θ distributed uniformly over the angular range $\delta\theta_{Max}$, corresponding to the linear range $\delta X_{T,i}$(0.1 ms) = $\delta X_{Max}$, after affine transformation with (J75). The 7 **red** points displayed identically in Figs J12a, J12b and J12c are determined according to (J74a) from the experimental values.

Knowing $\delta X_{T,i}$, the instant elastic slope $\chi_i$ is determined with (J65). On the basis of these elements, the parameter $e_i/e_0$ is calculated from (J71b) for each of the three models. The three respective plots appear in **red** on Figs J12a', J12b' and J12c'.

The 7 **red** dots, identical in Figs J12a', J12b' and J12c' come from Fig 10 in [7]. In all three cases, a good agreement is obtained between theoretical and experimental values ($r^2$ > 99.9%).

The three models are close and we find equality " $\delta X_{T,i}$= $\delta X_T$ " if $pT0_i$ > 15%.

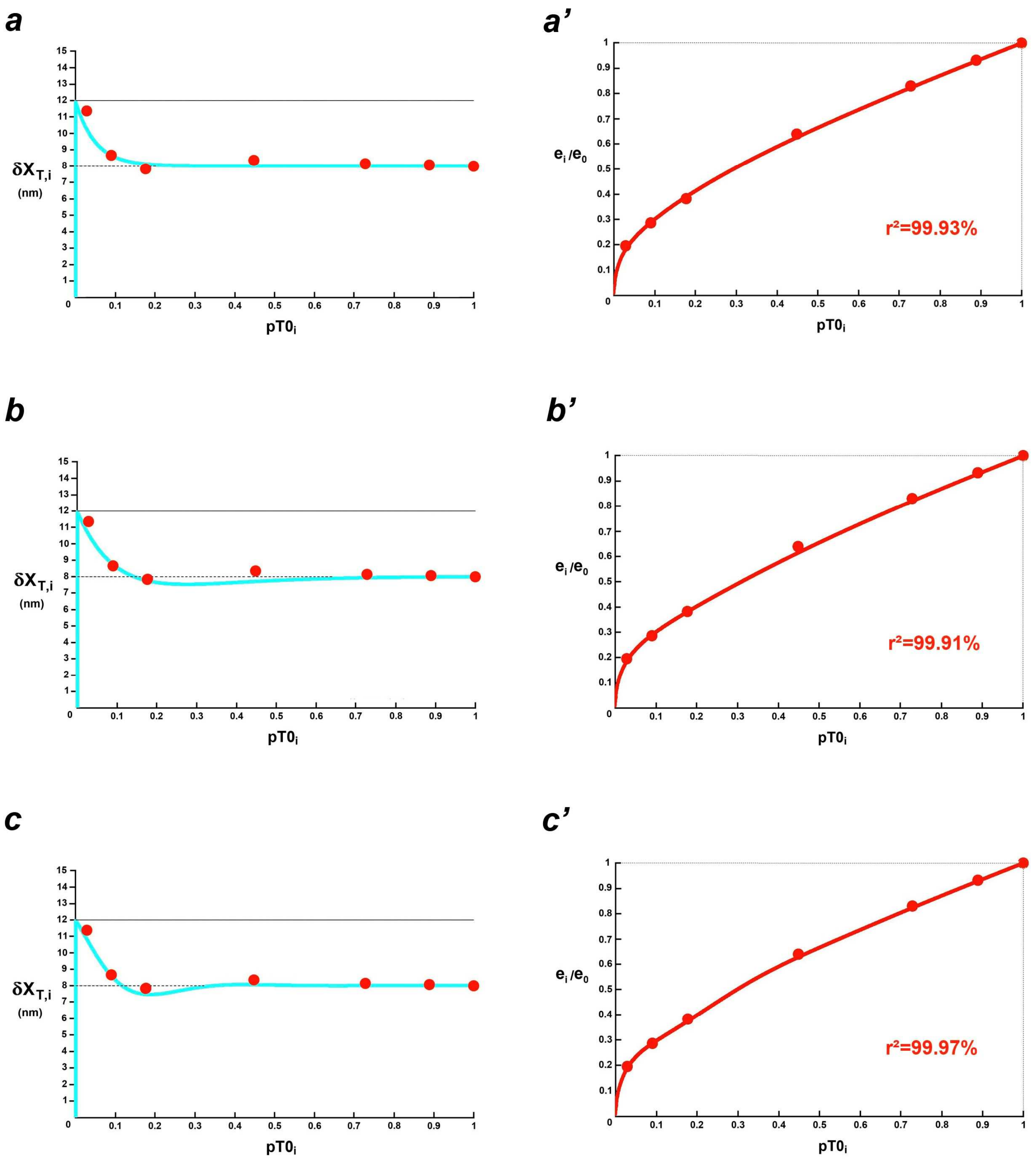


**Fig J12. (a), (b) and (c) Curves of $\delta X_{T,i}$ as a function of $pT0_i$ (blue line) according to the relationships (J82a), (J82b) and (J82c). The red points are calculated with (J74a) from the experimental values. (a'), (b') and (c') Corresponding curves of $e_i/e_0$ (red line) as defined with equation (J71b). The red dots are the experimental values during the isometric tetanus rise, represented by black circles on Fig 10 in [7].**

#### J.14 Study of the instant linear range ($\delta X_{T,i}$), the instant stiffness referred to $T0_c$ ($e_i/T0_c$) and the instant strain ($Y_i$) during tension rise to the tetanus plateau

Fibers are isolated from the *tibialis anterior* muscle of *Rana Esculenta*. During rise to the isometric tetanus plateau, each of the fibers is subjected to cycles of 10 sinusoids of 2 $nm \cdot hs^{-1}$ peak to peak amplitude at a frequency of 4 kHz, repeated cycles at 5 ms intervals. The corresponding data are displayed in the green column of Table 1 of Paper 4.

The $q_i$ value is evaluated according to (J63):

$$q_i = 1.835 + 0.118 \cdot Ln\,(pT0_i)$$

The corresponding theoretical factor $K_i$ is determined with (J66).

The linear range $\delta X_{T,i}$ is empirically modelled according to (J78a) after affine transformation using (J75):

$$\delta X_{T,i} = \delta X_T + (\delta X_{Max} - \delta X_T) \cdot \left[ e^{-10 \cdot pT0_i} \right] \cdot (1 - 25 \cdot pT0_i) \qquad \text{(J83)}$$

Equation (J83) is represented by a **blue line** in Fig J13a. The **purple** and **green** points are determined according to the reformatted equality (J74a) and according to (J74b) from the experimental values, respectively.

Knowing $\delta X_{T,i}$, we determine the instant elastic slope $\chi_i$ with (J65). The parameters $e_i / T0_c$ and $Y_i$ are calculated from (J71c) and (J71e). Their respective curves appear in continuous **purple** and **green** lines on Figs J13b and J13c.

The **purple** and **green** dots come, respectively, from Figs 3A and 3C in [8].

A good agreement between theoretical and experimental values is observed ($r^2 > 99.5\%$).

Under condition $pT0_i > 40\%$, equality " $\delta X_{T,i} = \delta X_T$ " is admissible and the instant slope of elastic origin ($\chi_i$) can be considered as constant. It is noteworthy that 40% is almost 3 times 15%, where 15%, is the value of the condition for the constancy of $\delta X_{T,i}$ found in the previous example (Figs J12a, J12b and J12c) also dedicated to the rise to the tetanus plateau. According to Fig 2B in [8], isometry requires about 50 ms during which the fiber is shortened by about 25 nm, a length that is twice as long as the stroke size of a myosin head ($\delta X_{Max}$). At time t=50ms, the instantaneous tetanus tension reaches about 50% of the tetanus plateau value. It is legitimate to speculate that the fiber shortening before effective isometry is the direct cause of the non-stationary critical phase present in Fig J13a between 0 to 50% $pT0_i$.

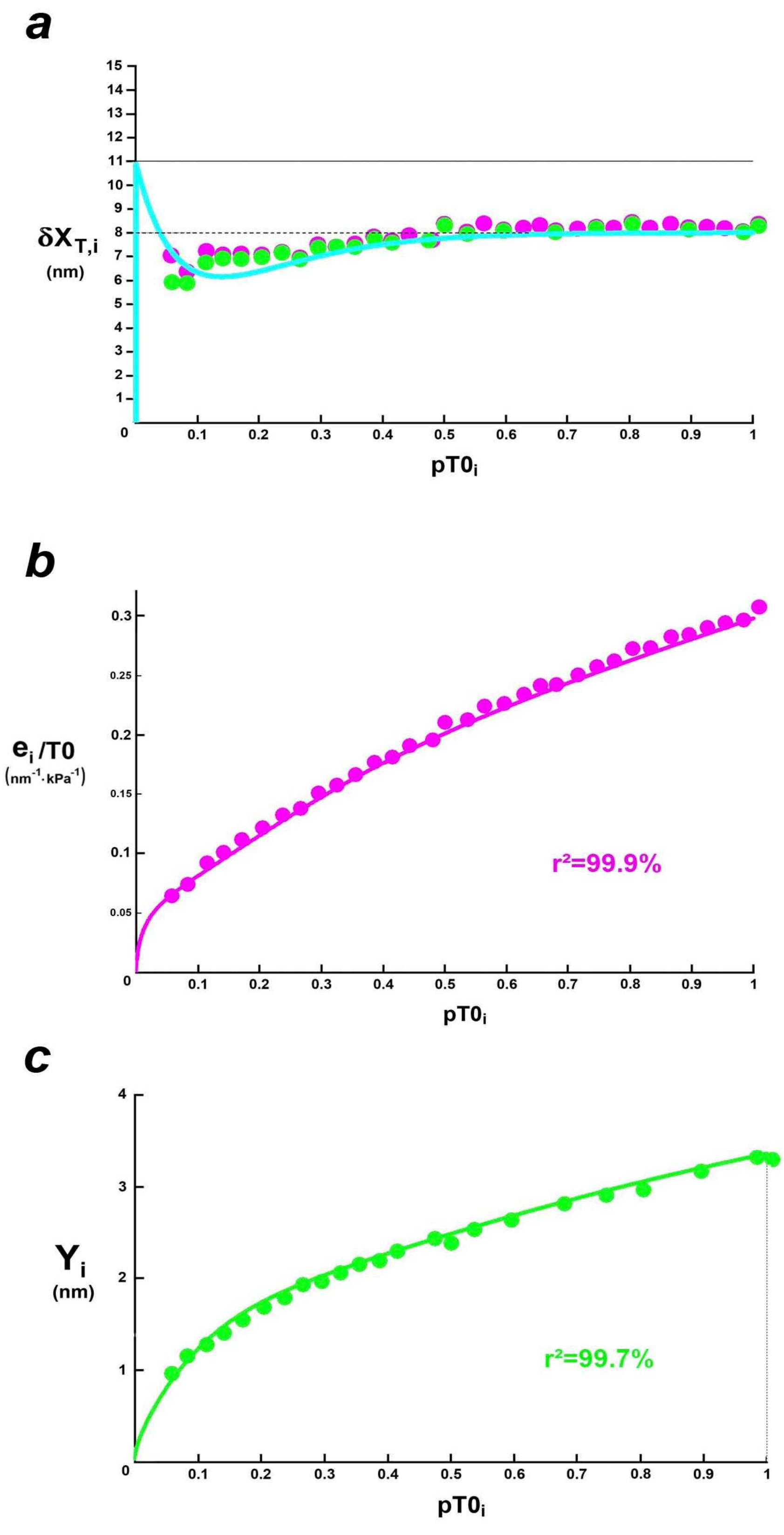


**Fig J13. (a) Relationship of $\delta X_{T,i}$ according to $pT0_i$ (blue line) according to (J83). The purple and green dots are derived from equations (J74a) and (J74b) after introducing the experimental values. (b) and (c) Relationships of $e_i/T0$ (purple line) and $Y_i$ (green line) as a function of $pT0_i$ according to equations (J71c) and (J71e). The purple and green experimental points come, respectively, from Figs 3A and 3C in [8].**

#### J.15 Study of the intermediate normalized stiffness ($e_i/e_0$) during phase 4 of a series of force steps with constant tension and shortening velocity

***J.15.1*** During phase 4 of the force steps, a fiber is tested using sinusoidal length oscillations (1.5 $nm{\cdot}hs^{-1}$ peak to peak) at a frequency of 4 kHz. From the data in the purple column of Table 1 of Paper 4, with the exception of $N_{hs}$ = 6450, the value of $q_i$ is evaluated with (J68):

$$q_i = \left[\left(1.99 + \frac{1}{Ln6450} \cdot \ln\frac{90\mu s}{200\mu s}\right) + \frac{1}{Ln6450} \cdot Ln(pT0_i)\right] = [1.9 + 0.114 \cdot Ln(pT0_i)]$$

The corresponding theoretical factor $K_i$ is determined according to (J66); it is noted that $K_0$=1.7.

The instant linear range ($\delta X_{T,i}$) is modelled by a sigmoid according to (J80) whose equation is provided in (36) in Paper 4 and reproduced below:

$$\delta X_{T,i} = \delta X_{Max} + \frac{\delta X_T - \delta X_{Max}}{1 + e^{-9 \cdot (pT0_i - 0.5)}} \quad \text{(J84a)}$$

The $\delta X_{T,i}$ curve according to (J84a) appears with a **light blue** line in the inset of Fig J14a. The **red** points of the insert are calculated from the expression (J74a) where the experimental values of $e_i/e_0$ collected on Fig 3A in [9] are entered.

Knowing $\delta X_{T,i}$ with (J84a), we determine the instant elastic slope ($\chi_i$) using (J65). The theoretical normalized slope ($e_i/e_0$) is calculated from (J71b) and its graphical representation appears with a solid **red** line in Fig J14a. The **red** dots are from Fig 3A in [9]. There is a good agreement between theoretical and experimental values.

***J.15.2*** Another fiber of the same type is used for another series of force steps under similar conditions. With $N_{hs}$ = 6900, the other data being identical, the value of $q_i$ is evaluated:

$$q_i = \left[1.9 + \frac{1}{Ln6900} \cdot Ln(pT0_i)\right] = [1.9 + 0.114 \cdot Ln(pT0_i)]$$

The corresponding theoretical factor $K_i$ is determined using (J66) and the instant linear range ($\delta X_{T,i}$) is modelled with an equation close to (J84a):

$$\delta X_{T,i} = \delta X_{Max} + \frac{\delta X_T - \delta X_{Max}}{1 + e^{-9 \cdot (pT0_i - 0.6)}} \quad \text{(J84b)}$$

Equation (J84b) is represented by a **light blue** line in the inset of Fig J14b. The **red-brown** points of the inset are determined according to (J74a) from the experimental values. Knowing $\delta X_{T,i}$ with (J84b), the instant elastic slope $\chi_i$ is determined and the parameter $e_i/e_0$ is calculated using (J71b); its curve appears in **red** in Fig J14b. The **red-brown** dots are from Fig 4A in [9]. There is a good agreement between theoretical and experimental values.

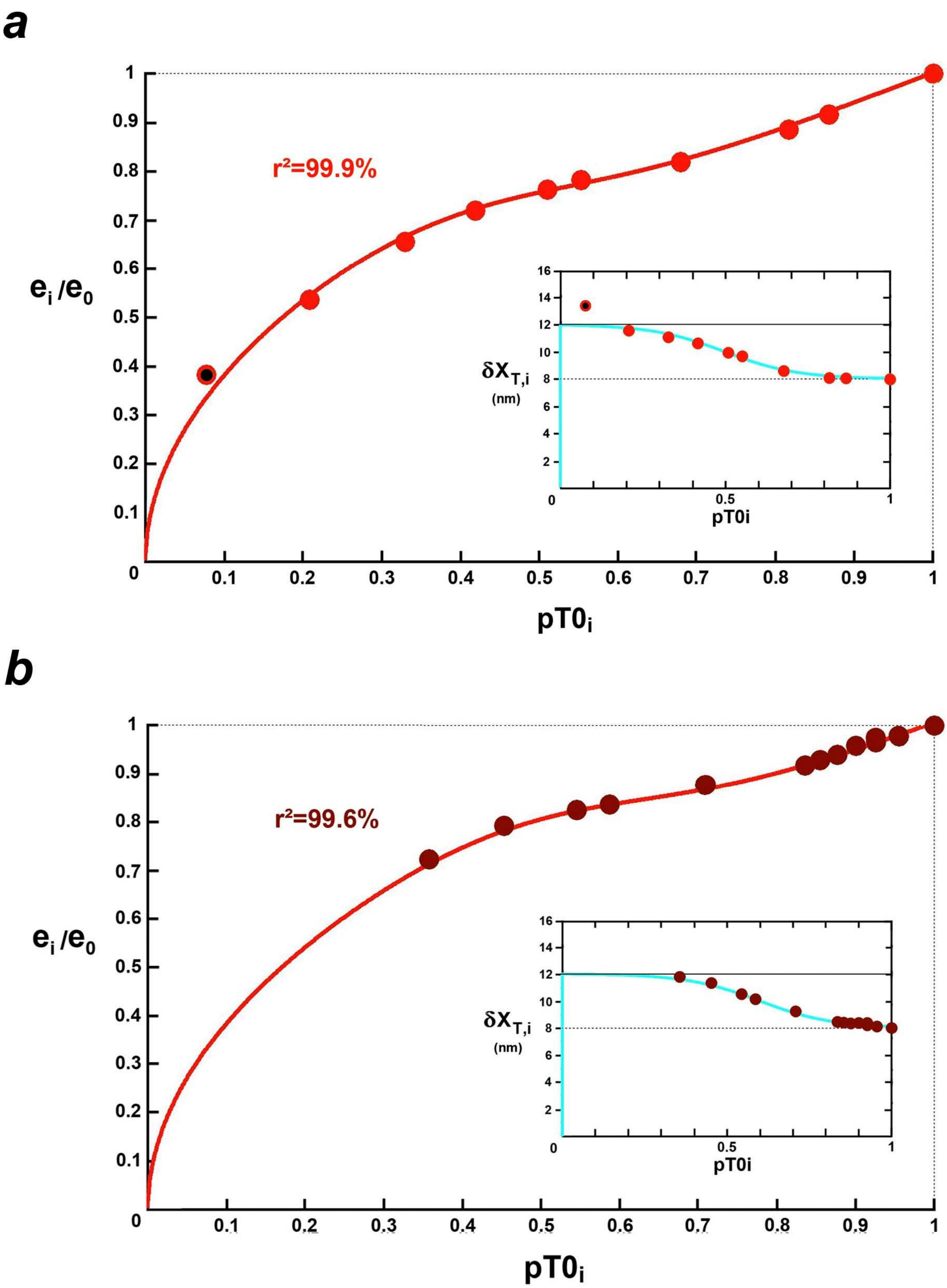


**Fig J14. (a) and (b) Relationships of $e_i/e_0$ (red solid line) as a function of $pT0_i$ according to (J71b). The red and red-brown dots are respectively from Figs 3A and 4A in [9].**
**In the insets, relations of $\delta X_{T,i}$ as a function of $pT0_i$ (light blue line) according to equations (J84a) and (J84b) for Figs J13a and J13b, respectively.**
**The two red and black dots in Fig J13a and in the inset correspond to a high hs shortening velocity (see explanations in text).**

***J.15.3*** Point n° 0 corresponds to the isometric tetanus plateau where $pT0_0 = (e_0/e_0) = 1$. The other points are numbered by increasing index according to the decreasing value of $pT0_i$.

In the inset of Fig J14a, we notice a value higher than $\delta X_{Max}$ : point n° 9 bicolor (**red** and **black**) corresponds to the lowest tension tested and to a high hs shortening velocity: $u_9 \approx 2\ nm{\cdot}ms^{-1}$ per hs according to Fig 3A in [9]. This apparent anomaly is due to the presence of viscosity which interferes with high shortening velocities (see accompanying Paper 1). The viscosity effects generated by $u_9$ are superimposed on the viscous forces generated by the oscillations.

In Fig J14a, point n° 9 bicolor (**red** and **black**) has the coordinates:

$$pT0_9 = 0.077 \quad \text{and} \quad e_{9,mes}/e_0 = 0.383$$

$K_{9,osc}$ is the multiplier coefficient determined with respect to point 9 when the actions of viscous origin come from oscillations alone.

The $q_9$ value come from sub-paragraph J.15.1:

$$pT0_9 = 0.077 \quad \Rightarrow \quad q_9 = [1.9 + 0.114 \cdot \mathrm{Ln}\,(0.077)] = 1.61$$

$K_{9,osc}$ is theoretically estimated with equality (J66):

$$K_{9,osc} = 6450^{0.2} \cdot \coth(6450^{0.2}) \approx 5.6$$

$K_{9,(osc+u9)}$ is the multiplier coefficient determined with respect to point n° 9 when the actions of viscosity are due to both forced oscillations and the high shortening speed ($u_9$).

The value of the linear range relative to point 9 ($\delta X_{T,9}$) is given by (J84a):

$$\delta X_{T,9} = 12 - \frac{4}{1 + e^{-9\cdot(0.077-0.5)}} \approx 11.91\ \mathrm{nm}$$

The original elastic slope ($\chi_9$) is evaluated with (J65):

$$\chi_9 = \frac{1}{12 - 11.91/2} = 0.165\ \mathrm{nm}^{-1}$$

With $\chi_0 = 0.125\ nm^{-1}$ and $K_0 = 1.7$, the factor $K_{9,(osc+u9)}$ is determined from the expression (J71b):

$$K_{9,(osc+u9)} = \left(\frac{e_{9,mes}}{e_0}\right) \cdot \left(\frac{\chi_0 \cdot K_0}{\chi_9 \cdot pT0_9}\right) = 0.383 \cdot \left(\frac{0.125 \cdot 1.7}{0.165 \cdot 0.071}\right) \approx 6.4$$

$K_{u9}$ is the multiplier coefficient induced only by the viscosity imposed by the high hs shortening velocity relative to point 9 ($u_9$).

The effects are added and the coefficients multiply: $K_{9,(osc+u9)}$ is the product of $K_{9,osc}$ and $K_{u9}$.

We deduce

$$K_{u9} = \frac{K_{9,(osc+u9)}}{K_{9,osc}} \approx 1.14$$

The figure is modest compared to $K_{9,osc}$.

*J.15.4* The sigmoid presented in the inset of Fig J13a essentially displays a three-phase shape:

- constancy if $0.8 \leq pT0_i \leq 1$ as already underlined by K.A. Edman [9]
- increasing linearity if $0.2 \leq pT0_i \leq 0.8$
- then again constant if $0 \leq pT0_i \leq 0.2$

#### J.16 Factors affecting viscosity forces

##### *J.16.1 Definition of* the *viscous origin slope [χ ·(K-1)]*

In paragraph (J.8), the calculations of the force generated by viscosity at the end of phase 1 of a length step lead to equations (J52a) and (J52b) which are formulated in a generic way:

$$pT1_{Visc} = [\chi \cdot (K-1)] \cdot \overline{\Delta X} \quad \text{(J85)}$$

It is the equation of a straight line passing through the origin with a slope equal to $[\chi \cdot (K-1)]$.

##### *J.16.2 Data for two frog species*

The impact by different factors on the equation (J85) is tested from data on the *tibialis anterior* muscle of two species of frogs for low experimental temperatures:

*rana Temporaria*

$T0_c = 250$ kPa $\quad N_{hs} = 5500 \quad \chi_{Z1} = 0.133$ nm$^{-1}$ $\quad \tau_{p1} = 0.2$ ms $\quad q_{Z1} = 1.99$

*rana Esculenta*

$T0_c = 200$ kPa $\quad N_{hs} = 4800 \quad \chi_{Z1} = 0.143$ nm$^{-1}$ $\quad \tau_{p1} = 0.12$ ms $\quad q_{Z1} = 1.88$

##### *J.16.3 Influence of the elastic origin slope*

The definition of equality (J15) requires:

$$q = \frac{Ln\chi - Ln\nu}{LnN_{hs}}$$

The previous expression is compared to (J23) which gives:

$$q = q_{z1} + \frac{1}{LnN_{hs}} \cdot Ln\left(\frac{\chi}{\chi_{z1}}\right)$$

The multiplier coefficient K associated with each value of χ is calculated using (J19):

$$K = N_{hs}^{1-q/2} \cdot \coth\left(N_{hs}^{1-q/2}\right) + N_{hs}^{1-q}$$

It is thus possible to calculate the original viscous origin slope $[\chi(K-1)]$ of the linear equation (J85) for each value of χ. The relationship of $[\chi \cdot (K-1)]$ to χ is plotted in Fig 15a where χ ranges from 0 to 0.2 nm$^{-1}$. It is noted that the elastic origin slope relating to *rana Temporaria* remains almost constant for $\chi \geq 0.05$ nm$^{-1}$, confirming the proximity of the values of $[\chi_{z1} \cdot (K_{z1}-1)]$ and $[\chi_{z2} \cdot (K_{z2}-1)]$ viewed in paragraph (J.8).

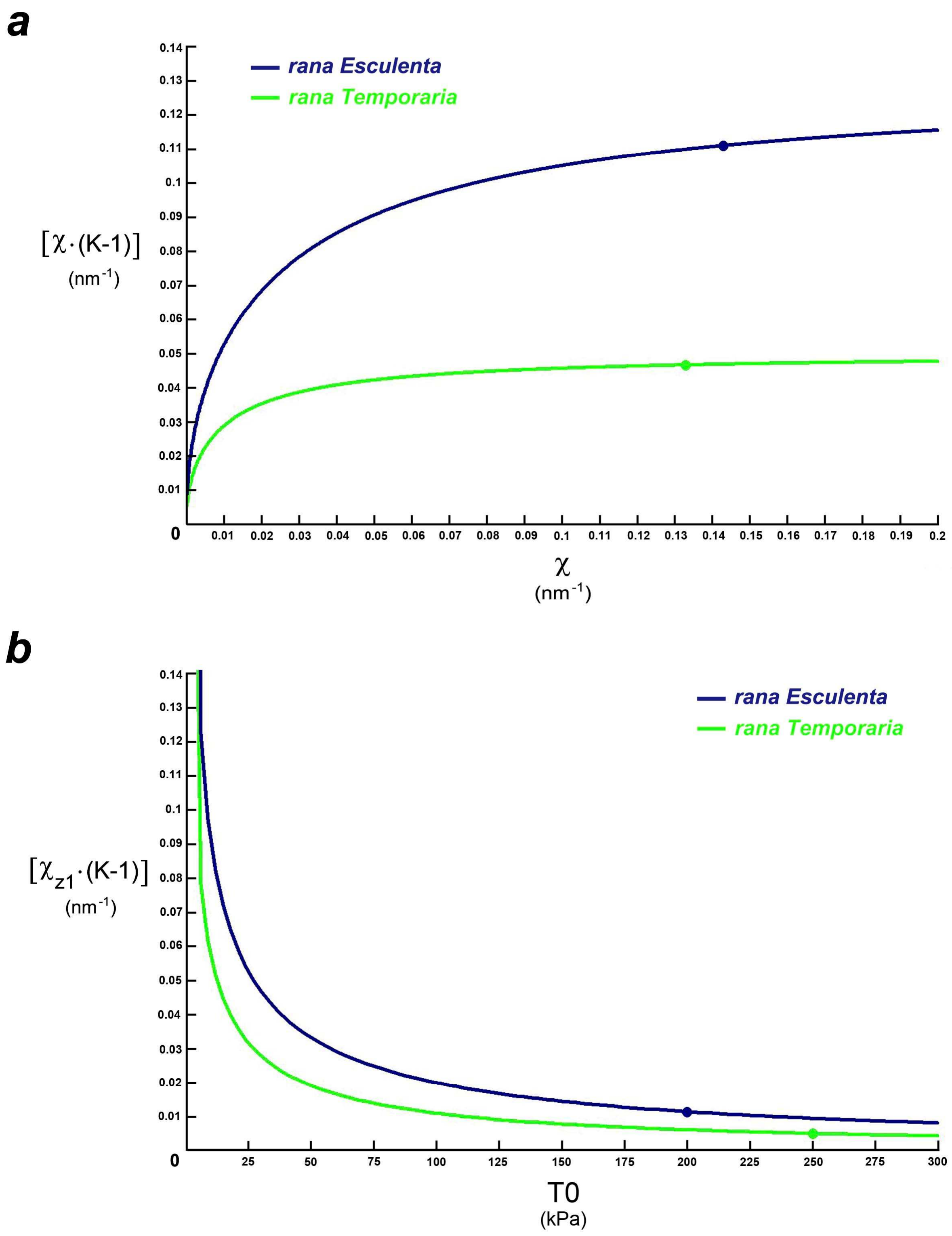


**Fig J15. (a) Relation of the viscous origin slope $[\chi \cdot (K\text{-}1)]$ as a function of the elastic origin slope ($\chi$). (b) Relation of $[\chi_{z1} \cdot (K\text{-}1)]$ as a function of isometric tetanus tension (T0).**
**The points on the plots correspond to the reference values for two fibers belonging to two frog species (see data in sub-paragraph J.16.2).**

##### *J.16.4 Influence of isometric tetanus tension (T0)*

For each new experiment associated with a particular value of T0, the parameter q is estimated using (J57):

$$q = q_c + \frac{1}{LnN_{hs}} \cdot Ln\left(\frac{T0}{T0_c}\right)$$

where $q_c$ and $T0_c$ are the reference values for the control experiment; see examples given in sub-paragraph J.16.2 for two frog species

The corresponding multiplier coefficient K is determined according to (J23). By setting the value of $\chi$ equal to $\chi_{z1}$, the viscous origin slope [$\chi_{z1}$·(K-1)] is calculated for each value of T0. The plot of $\chi_{z1}$·(K-1) as a function of T0 is shown in Fig 15b for two frog species.

As T0 decreases, the viscous origin slope increases in accordance with observations where viscosity is more significant for a fiber at rest (T0 ≈ 0) than stimulated.

##### *J.16.5 Influence of the duration of Phase 1*

As phase 1 of a length step is performed at constant velocity, we pose:

$$u = \frac{\overline{\Delta X}}{\tau_{p1}} \tag{J86}$$

where u is the average hs shortening velocity; $\tau_{p1}$ is the duration of phase 1 common to all length steps.

The introduction of (J86) in (J85) provides a linear relationship between the viscosity force ($pT_{Visc}$) and u:

$$pT_{Visc} = \varepsilon \cdot u \tag{J87}$$

where ε is a proportionality coefficient verifying:

$$\varepsilon = \lfloor \chi \cdot (K-1) \cdot \tau_{p1} \rfloor \tag{J88}$$

For each distinct experiment associated with a particular value of $\tau_{p1}$, the parameter q is estimated using (J54):

$$q = q_c + \frac{1}{LnN_{hs}} \cdot Ln\left(\frac{\tau_{p1}}{\tau_{p1,c}}\right)$$

where $q_c$ and $\tau_{p1,c}$ are the reference values for the control experiment; see examples given in sub-paragraph J.16.2 for two frog species.

The corresponding multiplier coefficient K is determined according to (J23). With the constant elastic origin slope ($\chi$) equal to $\chi_{z1}$, the coefficient ε is calculable with (J88) and the plot of ε as a function of $\tau_{p1}$ varying between 0 and 3 ms is presented in Fig 16 for the fibers relating to the two frog species.

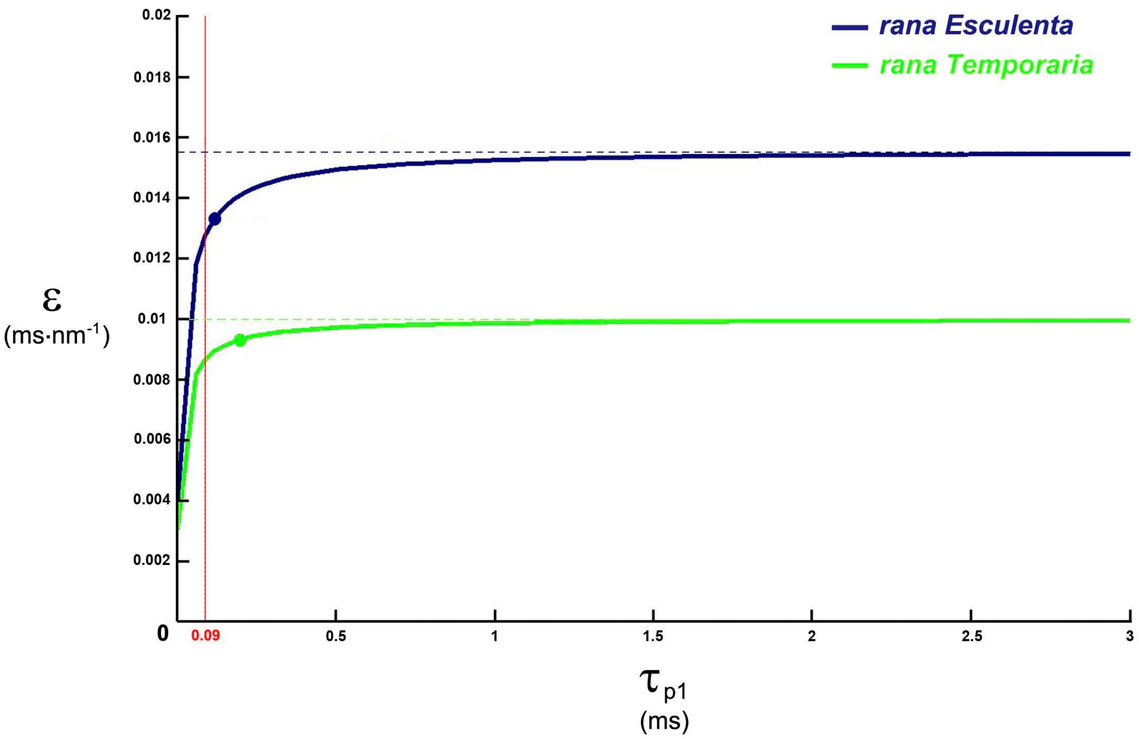


**Fig J16. Relation of the proportionality coefficient (ε) according to the duration of phase 1 of a length or force step ($\tau_{p1}$). The two points on the two curves correspond to the two reference values for the two types of fibers (see sub-paragraph J.16.2).**

It may seem paradoxical that $\tau_{p1}$ is a multiplicative factor of ε because it is noted in paragraph J.9 that the more $\tau_{p1}$ increases, the more the presence of viscosity decreases. In fact, the 2 parameters $\tau_{p1}$ and K are interdependent through equations (J54) and (J56): if one increases, the other decreases, and vice versa.

In accordance with (J46), the value " $\tau_{p1}$= 90 µs" corresponding to oscillations at 4 kHz is represented by a red vertical line.

If $\tau_{p1} \geq 200$ µs, the coefficient ε is in first approximation considered as constant and independent of the value of $\tau_{p1}$. For fibers isolated from *tibialis anterior* muscle of two frog species, we fix:

*rana Temporaria* : $\varepsilon \approx 0.01\ \text{ms}\cdot\text{nm}^{-1}$

*rana Esculenta* : $\varepsilon \approx 0.015\ \text{ms}\cdot\text{nm}^{-1}$

##### *J.16.6 Influence of temperature*

The fiber is tested at different intermediate temperatures ($\Gamma_i$ ). Assuming that the angular ranges $\delta\theta_{Max}$ and $\delta\theta_T$ remain unchanged, the elastic origin slope ($\chi_i$) associated with each value of $\Gamma_i$ is a constant equal to $\chi_{z1}$. Equality (J71e) brings:

$$K_i = \frac{1}{\chi_{z1} \cdot Y_i} \tag{J89}$$

With (J89), the viscous proportionality coefficient ($\varepsilon_i$) associated with $\Gamma_i$ checks using (J89):

$$\varepsilon_i = \left[\left(\frac{1}{Y_i} - \chi_{z1}\right) \cdot \tau_{p1}\right] \tag{J90}$$

From the values of $\Gamma_i$, $T0_i$ and $Y_i$ published for *rana Esculenta* [10,11], those of $K_i$ and $\varepsilon_i$ are determined using (J89) and (J90). The data are grouped in Table J1. The viscous coefficient $\varepsilon_i$ decreases with temperature [10,11].

The intermediate stiffness $e_i$ is evaluated according to (J71a) and its value is almost constant for the 8 temperatures, equal to or close to 40 $\text{kPa}\cdot\text{nm}^{-1}$ (Table J1). Consequently, in Zone 1 Enlarged, the line segments representing T1 as a function of ΔX appear parallel for different temperatures; this is verified experimentally. The decrease in the influence of viscosity via $K_i$ is compensated by the increase in $T0_i$ generated by an intensification of WS possibilities (see supplement S3.H of accompanying Paper 3) associated with an increase in enthalpies free of cross-bridge cycle reactions, especially those concerning the strong binding and working stroke.

An exponential regression between $\varepsilon_i$ and $\Gamma_i$ presented in Table J1 establishes the following relationship ($r^2$=99.4%):

$$\varepsilon_i = 0.0216 \cdot e^{-0.109 \cdot \Gamma_i} \tag{J91}$$

An evaluation of $\varepsilon_i$ corresponding to the example in Fig 6 of Paper 1 from [12] is provided in Table J2.

**Table J1. Calculation of the viscous proportionality coefficient ($\varepsilon_i$) and the stiffness ($e_i$) as a function of the experimental temperature ($\Gamma_i$) for fibers extracted from the *anterior tibialis* muscle in *rana Esculenta*.**

| | $\Gamma_i$ (°C) | $T0_i$ (kPa) | $Y_i$ | $K_i$ (1) | $\varepsilon_i$ (2) ($ms \cdot nm^{-1}$) | $e_i$ (3) ($kPa \cdot nm$) |
|---|---|---|---|---|---|---|
| Table 2 and Fig 3C in **Piazzesi 2003** | 2 | 139.5 | 3.5 (4) | 2 | 0.017 | 40 |
| | 5 | 165 | 4 (4) | 1.7 | 0.012 | 40 |
| | 10 | 198 | 4.75 (4) | 1.4 | 0.007 | 40 |
| | 17 | 233 | 5.9 (4) | 1.2 | 0.0035 | 40 |
| Table 1 and Fig 1B in **Decostre 2005** | 2 | 144 | 3.5 | 2 | 0.017 | 41 |
| | 5 | 172 | 4 | 1.75 | 0.012 | 43 |
| | 10 | 197 | 4.75 | 1.5 | 0.007 | 41.5 |
| | 17 | 226 | 5.9 | 1.2 | 0.0035 | 38 |

(1) calculated from (J89) with $\chi_{z1}$= 0.143 $nm^{-1}$
(2) calculated from (J90) with $\tau_{p1}$ = 0.12 ms
(3) calculated from (J71a) with $\chi_i = \chi_{z1}$
(4) calculated with $K_i = 0.025 \cdot T0_i$

**Table J2. Calculation of the viscous proportionality coefficient ($\varepsilon_i$) as a function of the experimental temperature ($\Gamma_i$).**

| | $\Gamma_i$ (°C) | $\varepsilon_i$ (1) ($ms \cdot nm^{-1}$) |
|---|---|---|
| Fig 1 in **Elongovan 2012** (Rana E – force steps) | 7 | 0.01 |
| | 15 | 0.004 |
| | 21 | 0.002 |

(1) calculated from (J91)

##### *J.16.7 Influence of inter-filament space*

The tonicity enrichment of the *Ringer* solution increased from 1*R* à 1.44 *R* osmotically compresses the cell fluid and reduces the inter-filament space.

**At normal tonicity (1*R*)**, the analysis of Fig 2A in [13] provides the following data relative to a fiber isolated from the *tibialis anterior* muscle of *rana Esculenta*:

$T0_{1R} = 210$ kPa  $N_{hs} = 4600$  $\tau_{p1} = 0.11$ ms  $\chi_{1R} = 0.143\ nm^{-1}$  $Y_{1R} = 4.3$ nm

Equality (J71e) delivers:

$$K_{1R} = K_0 = 1/(0.143 \cdot 4.3) = 1.63$$

The parameter $q_{1R}$ associated with $K_{1R}$ is calculated by par interpolation of the expression (J59):

$$q_{1R} = q_0 = 1.91$$

And equality (J89) brings with the previous data:

$$\varepsilon_{1R} = (0.143 \cdot 0.63 \cdot 0.11) = 0.01\ ms \cdot nm^{-1}$$

**At a hypertonicity (1.44 *R*)**, the study of Fig 2A in [13] gives:

$$T0_{1.44R} = 0.77 \cdot T0_{1R}$$
$$Y_{1.44R} = 3.5\ nm$$

In hypertonicity, the equation (J63) implies:

$$q_{1.44R} = q_{1R} + \frac{1}{Ln4600} \cdot Ln[0.77] = 1.88$$

The expression (J66) gives:

$$K_{1.44R} = 1.78$$

Equality (J74b) issues:

$$\begin{aligned}\delta X_{T,1.44R} &= 2 \cdot (\delta X_{Max} - Y_{1.44R} \cdot K_{1.44R}) \\ &= 2 \cdot (11 - 3.5 \cdot 1.78)\ nm \\ &= 9.5\ nm\end{aligned}$$

The original elastic slope ($\chi_{1.44R}$) is calculated according to (J65):

$$\chi_{1.44R} = \frac{1}{\delta X_{Max} - \delta X_{T,1.44R}/2} = 0.16\ nm^{-1}$$

And the viscous coefficient in hypertonicity is determined with (J89):

$$\varepsilon_{1.44R} = (0.16 \cdot 0.78 \cdot 0.11) = 0.014\ ms \cdot nm^{-1}$$

With respect to a hypertonicity of 1.44R, the angular range ($\delta\theta_{T,1.44R}$), linear range ($\delta X_{T,1.44R}$), elastic origin slope ($\chi_{1.44R}$) and viscous proportionality coefficient ($\varepsilon_{1.44R}$) have values higher than those of the corresponding parameters relatively to normal tonicity (1R), i.e. when the inter-filament distance decreases.

The data are those used for the Force-Velocity relationship plots in Fig 4 to Paper 1.

***J.16.8 Influence of the tension measurement method at the end of phase 1 of a length step***

The study is based on Fig 6B in [14]. The figure refers to the study of a fiber taken from the *semi-tendinosus* muscle of *Rana Temporaria* tested under different conditions. Under standard norms (2°C), we pose:

$T0 = 170$ kPa $\quad N_{hs} = 5500 \quad \chi_{z1} = 0.133$ nm$^{-1}$ $\quad \tau_{p1} = 0.14$ ms

If the line passes through the minimum tension value (T1), it intersects the abscissa axis in:

$$Y_{0,T1} = 4.4 \text{ nm}$$

The viscous proportionality coefficient is calculated with (J90) and the previous data:

$$\varepsilon = \left[\left(\frac{1}{Y_{0,T1}} - \chi_{z1}\right) \cdot \tau_{p1}\right] \approx 0.013 \text{ ms·nm-1}$$

With the instantaneous values plotted using squares, the intersection of the line with the abscissa axis gives:

$$Y_{0,inst} = 3.7 \text{ nm}$$

$$\varepsilon = \left[\left(\frac{1}{Y_{0,inst}} - \chi_{z1}\right) \cdot \tau_{p1}\right] \approx 0.019 \text{ ms·nm-1}$$

That is an increase for ε of nearly 50%. This allows to judge the importance of the choice that governs the collection of measurements.

We have retained a compromise value with $Y_0 = 4$ nm and $\varepsilon \approx 0.016$ ms·nm$^{-1}$ for the plots of the Force-Velocity relationships relative to Fig 5 of Paper 1.

#### Références du Supplément S4.J du PAPIER 4
